# The structure of the tetraploid sour cherry ‘Schattenmorelle’ (*Prunus cerasus* L.) genome reveals insights into its segmental allopolyploid nature

**DOI:** 10.1101/2023.03.28.534503

**Authors:** Thomas W. Wöhner, Ofere F. Emeriewen, Alexander H.J. Wittenberg, Koen Nijbroek, Rui Peng Wang, Evert-Jan Blom, Jens Keilwagen, Thomas Berner, Katharina J. Hoff, Lars Gabriel, Hannah Thierfeldt, Omar Almolla, Lorenzo Barchi, Mirko Schuster, Janne Lempe, Andreas Peil, Henryk Flachowsky

## Abstract

Sour cherry (*Prunus cerasus* L.) is an economically important allotetraploid cherry species believed to have evolved in the Caspian Sea and Black Sea regions. How, when and where exactly the evolution of this species took place is unclear. It resulted from a hybridization of the tetraploid ground cherry (*Prunus fruticosa* Pall.) and an unreduced (2n) pollen of the diploid ancestor sweet cherry (*P. avium* L.). Some indications implement that the genome of sour cherry is segmental allopolyploid, but how it is structured and to what extent is unknown. To get an insight, the genome of the sour cherry cultivar ‘Schattenmorelle’ was sequenced at ~400x using Illumina NovaSeq^TM^ short-read and Oxford Nanopore long-read technologies (ONT R9.4.1 PromethION). Additionally, the transcriptome of ‘Schattenmorelle’ was sequenced using PacBio Sequel II SMRT cell sequencing at ~300x. The final assembly resulted in a ~629 Mbp long pseudomolecule reference genome, which could be separated into two subgenomes each split into eight chromosomes. Subgenome *Pce*_S__a which originates from *P. avium* has a length of 269 Mbp, whereas subgenome *Pce*_S__f which originates from *P. fruticosa* has a length of 299.5 Mbp. The length of unassembled contigs was 60 Mbp. The genome of the sour cherry shows a size-reduction compared to the genomes of its ancestral species. It also shows traces of homoeologous sequence exchanges throughout the genome. Comparative positional sequence and protein analyses provided evidence that the genome of sour cherry is segmental allotetraploid and that it has evolved in a very recent event in the past.

## Background

Cherries include several species of the genus *Prunus*, which belong to the sub-family Spiraeoideae in the plant family Rosaceae (Potter et al. 2007). Although the fruits of several cherry species are used for consumption, only a few of them are grown and marketed on an economically significant scale (Quero-García et al. 2019). The two economically most important cherry species worldwide are the sweet cherry (*Prunus avium* L.) and the sour cherry (*Prunus cerasus* L.). Both species are thought to have originated in the Caspian Sea and Black Sea region (Quero-García et al. 2019).

Sour cherries commercial cultivation is mainly localized in Eastern and Central Europe, North America, and Central and Western Asia on an area of 217,960 ha. They are mainly cultivated for processing jams, juices, and whole fruits in preserved or dried forms. Furthermore, they are used in dairy products and baked goods. Their global production in 2021 was 1.51M tons of fruit and the cross production value in 2020 was 1.2 billion $US (https://www.fao.org/faostat/en/#data). The variability of morphological and fruit characteristics is very high in sour cherry, including fruit and juice colour, size, firmness, and fruit compounds (Schuster et al. 2017). Furthermore, tree growth varies from rather slender and upright to small and bushy types. Variation within ecotypes, differing e.g. in cold tolerance or growth habit, have been selected across Europe over time (Dirlewanger et al. 2007; Hancock 2008). However, just a small number of cultivars actually dominates the cultivation of sour cherry. ‘Schattenmorelle’ is the dominant cultivar (cv) in Middle Europe (Figure 1), whereas sour cherry production in the United States is still based on ‘Montmorency’ (Quero-Garcia et al. 2019).

**Figure 1.** Morphology of *P. cerasus* L. ‘Schattenmorelle’. (a) mature tree habitus, (b) leaves, (c) inflorescence, (d) fruits.

’Schattenmorelle’ was first described in France and today it is known in many countries with different names. In Poland, for example, it is called ‘Łutovka’ and in France ‘Griotte du Nord’ or ‘Griotte Noir Tardive’.

The sour cherry is an allotetraploid with 2n=4x=32 chromosomes. It originated as a hybrid of an unreduced 2n pollen grain of *P*. *avium* (2n=2x=16) and a 1n egg cell of the tetraploid ground cherry *P*. *fruticosa* (2n=4x=32) (Kobel 1927; Olden and Nybom 1968). Evidence of hybridization events between sweet and ground cherries has already been found several times in areas where both species occur simultaneously (Macková et al. 2018, Hrotkó et al. 2020). The resulting hybrids were usually triploid and were assigned to the secondary species *P.* ×*mohacsyana* Kárpáti. Natural occurrences of tetraploid sour cherries can be found in Eastern Turkey and the Caucasus region. There, they grow in forests and are used as wild forms for fruit production. The real area of origin is not known so far. Although *P. cerasus* can also be found in the wild in Europe, it is rather unlikely that those sour cherries are spontaneous hybrids. Since sour cherries are cultivated almost in many areas of the Northern hemisphere, they are often rather allochthonous individuals. The origin of the sour cherry thus seems to be based on a few hybridisation events. The results obtained by Olden and Nybom (1968) in experiments on the resynthesis of the species *P. cerasus* confirmed this hypothesis. The progeny from crosses between sweet and ground cherry showed the characteristic phenotype of the sour cherry. Studies based on chloroplast DNA markers strongly suggest that hybridisation between *P. avium* and *P. fruticosa* led to the emergence of *P. cerasus* at least twice (Dirlewanger et al. 2007). Furthermore, the hypothesis could also be confirmed by genomic *in situ* hybridisation (Schuster and Schreiber 2000) and transcriptome sequencing (Bird et al. 2022).

The sour cherry genome is presumed to have a genome size of 599 Mbp (Dirlewanger et al. 2007). It consists of the two subgenomes, each with eight chromosomes in the haploid set of chromosomes. One subgenome comes from the sweet cherry (*Pce*_a), while the other is from the ground cherry (*Pce*_f). However, the genome does not appear to be completely allopolyploid, since it has long been suspected that parts of the sour cherry genome might be segmental (Beaver and Lezzoni 1993; Olden and Nybom 1968; Raptopoulus 1941; Schuster and Wolfram 2005). The impact of hybridization and polyploidization between sweet and ground cherry has not been investigated so far, nor when this event originates. Cai et al. (2018) assumes that a mix of multi- and bivalent pairing led to imbalance segregation of chromosomes during meiosis in sour cherry, which in turn suggests that the sour cherry genome has not yet stabilized (Mason et al. 2020), or is still in the process of doing so. Recent advances in genome sequencing shed light into the complex structure and shape of polyploid genomes and their evolution (Zhang et al. 2021, Edger et al. 2019, Bertioli et al. 2019, Wu et al. 2021, Wang et al. 2019).

Here we report a high quality pseudo-chromosome-level genome assembly of the tetraploid sour cherry ‘Schattenmorelle’ (hereinafter referred to as *Pce*_S_) generated with a combination of Illumina NovaSeq short-read and Oxford Nanopore long-read sequencing technology. The sequences were scaffolded to chromosomes by Hi-C. In parallel, a full-length transcriptome of ‘Schattenmorelle’ was generated with the Pac-Bio Sequel II SMRT cell long-read technology. Comparative sequence and amino acid analyses between data sets of *Prunus avium* cv ‘Tieton’ (hereinafter referred to as *Pa*_T_) and *Prunus fruticosa* ecotype Hármashatárhegy (hereinafter referred to as *Pf*_eH_) as representatives of the two ancestral species (Wang et al. 2019, Wöhner et el. 2021) and the two subgenomes of ‘Schattenmorelle’ *Pce*_S__a and *Pce*_S__f shed light into the evolution of sour cherry. Finally, possible HE within the sub-genomic structure of sour cherry were spotted, explaining the sour cherry genome segmental allopolyploidy.

## Results

### De novo assembly and scaffolding

A total of 68 Gb of paired-end Illumina sequencing data was obtained, corresponding to ~114x coverage of the estimated genome size of 599 Mbp. Using two PromethION flow cells, a total of 178 Gb was produced (~300x coverage). The longest ONT reads that together resulted in a 20x coverage were selected for assembly, having a minimum read length of 64,214 bp. Table S1 summarizes the properties of the 20-WGS-PCE.1.0 assembly after polishing. The *Prunus avium* and *Prunus fruticosa* contigs were then separated successfully by read mapping and contig selection that fit the hypothesis of 1 or more clear coverage peaks from the 20-WGS-PCE.1.0 assembly. The resulting two datasets, representing the subgenomes *Pce*_S__a and *Pce*_S__f, were purged and used for scaffolding using HI-C. After manual curation of both datasets, the final consensus genome assembly was scaffolded from 935 and 865 contigs of the *Pce*_S__a and *Pce*_S__f subgenomes, respectively. Eight clusters ideally representing the eight chromosomes were obtained for each subgenome (Figure S1). The final genome sequence is 628.5 Mbp long and consists of eight chromosomes for each subgenome (Figure 2). A total of 269 Mbp were assigned to subgenome *Pce*_S__a (N50 of 31.5 Mbp) and 299.5 Mbp (N50 of 39.4 Mbp) to *Pce*_S__f. Eighty-six and 134 unassembled contigs were unassigned to chromosomes for *Pce*_S__a (22.7 Mbp) and *Pce*_S__f (37.3 Mbp), respectively.

**Figure 2.** The genome of *P.cerasus* ‘Schattenmorelle’. Circos plot of 16 pseudomolecules of the subgenomes of *Pce_S_*_a and *Pce_S_*_f. (a) chromosome length (Mb); (b) gene density in blocks of 250k: (c) distribution of repetitive sequences in blocks of 250k (d) Gypsy elements in blocks of 250k; (e) Copia elements in block of 250k; (f) GC content in blocks of 1 Mb. (g) The inner ring shows markers from the 6+9k SNP array located on both subgenomes.

The longest scaffold from *Pce*_S__a is 52.8 Mbp and 53.5 Mbp from subgenome *Pce*_S__f (Table S2). Except for chromosome five, all scaffolds obtained from subgenome *Pce*_S__f are longer compared to the corresponding chromosome of subgenome *Pce*_S__a. The chloroplast sequence obtained was 158,178 bp and the mitochondrial sequence was 343,516 bp long (Figure S2).

### Transcriptome sequencing, isoseq-analysis, structural and functional annotation

The total repeat content of the entire *Pce*_S_ genome sequence was 49.7%. The total repeat content of subgenome *Pce*_S__a was 48.3% and of subgenome *Pce*_S__f 50.9%, respectively (Table 1). The class I elements Gypsy comprised the largest fractions of repetitive elements in the *Pce*_S_ genome sequence. A quantitative reduction between elements of this family was also detected in the *Pce*_S__f subgenome with a difference of 10.7%. Several elements could only be detected in one genotype of the two ancestral species. The TAD1 class I element only occurred in *Pf*_eH_, while class II, order TIR - IS3EU, P, and Sola-3 were specifically detected in the genome of *Pa*_T_. No element was found, which was only present in one of the two subgenomes of *Pce*_S_. Several elements occurred in both subgenomes (class I, LINE – R1-LOA, RTE-X, SINE – tRNA-DEU-L2, class II, TIR – TcMar-Mariner, and DADA elements) but were not detected in *Pf*_eH_ and *Pa*_T_. The class I elements of the order LTR (ERV1, Pao) and Academ/-2 were only detected in one of the two genomes representing the ancestral species and in *Pce*_S__a and *Pce*_S__f. Iso-Seq results are summarized in Table S3. In total 248,218 high quality isoforms have been identified. Both, the high and the low quality isoforms have been used for genome annotation where each gene might be represented by multiple isoforms. A total of 107,508 transcripts (*Pce*_S__a: 53,497; *Pce*_S__f: 54,011) were predicted from the 60,123 gene models (*Pce*_S__a: 29,069; *Pce*_S__f: 31,054) obtained by structural annotation procedures (Table S4). Interproscan analysis detected 1,381,841 functional annotations (*Pce*_S__a: 649,310; *Pce*_S__f: 687,531) using 16 databases. Two third (71,870) of the transcripts were assigned with GO terms and 9,114 were found to be involved in annotated pathways.

**Table 1.** Characterization of repetitive sequences of P. fruticosa ecotype Hármashatárhegy (Pf_eH_) compared to P. avium ‘Tieton’ (Pa_T_), P. persica ‘Lovell’, and the two subgenomes of P. cerasus ‘Schattenmorelle’ Pce_S__a and Pce_S__f

### Completeness and quality of the genome and transcriptome

BUSCO completeness of the *Pce*_S_ genome was 99.0% (S: 16.7%, D: 82.3%, F: 0.4%, M: 0.6%, n: 1,614) respectively and comparable with *P. persica ‘*Lovell’ (99.3%) and *P. avium ‘*Tieton’ (98.3%, Figure S3). Completeness of subgenome *Pce*_S__a was higher (C: 89.4%, S: 84.8%, D: 4.6%, F: 1.5%, M: 9.1%, n: 1614) compared to subgenome *Pce*_S__f (C: 87.1%, S: 80.9%, D: 6.2%, F: 1.2%, M: 11.7%, n: 1,614). The calculated LAI index was 6.3 and low in comparison to other genomes (*Pp*_L_: 17.6, *Pa*_T_: 10.3, *Pf*_eH_: 13.1). The LAI index for subgenome *Pce*_S__a was 7.1. The LAI index for subgenome *Pce*_S__f was 5.6 (Figure S4). The nucleotide heterozygosity rates were 94.9% for aaaa, 2.39 for aaab, 2.4 for aabb; 0.001 for aabc and 0.308 for abcd (Figure 3). The comparison of genetic position and physical position of up to 1,856 markers of the five genetic sour cherry maps (Table S5a) showed a good co-linearity to the genome sequence (Figure S5). Busco evaluation on completeness of the annotated proteins resulted in 99.2% [C: 99.2% (S: 8.4%, D: 90.8%), F: 0.4%, M: 0.4%, n: 1,614]. The chloroplast sequence obtained contained 427 genes, 21 rRNAs and 136 tRNAs, whereas the mitochondrial sequence contained 188 genes, 3 rRNAs and 152 tRNAs (Figure S2). An ab-initio and homology-based gene prediction with 14 reference species was performed (IAA). Based on the homology prediction, thirty-four percent of the proteins showed the highest IAA towards *Prunus fruticosa* and 17.9% towards *P. avium*. Only 5.2% of the proteins showed no IAA to any of the used reference datasets used, which was due to ab-initio prediction. The data is summarized in Figure S6.

**Figure 3.** GenomeScope (Galaxy Version 2.0) estimation of the *P. cerasus* genome size by k-mer counts obtained from the software Meryl (Galaxy Version 1.3+galaxy2). Both programs are integrated on the GalaxyServerEurope. The k-mer-peaks indicate that k-mers with a length of 19 bp occur in heterozygote (100x depth, 200x depth, 300x depth) and homozygote (400x depth) constitution within the genome. Coverage depth of individual k-mers is assigned as coverage.

A comparison of transcripts of *Pce*_S_ and the annotation datasets of *Pf*_eH_ and *Pa*_T_ enabled a quantitative comparison of shared transcripts within the datasets (Table 2). A total number of 26,532 shared transcripts were found between the two subgenomes *Pce*_S__a and *Pce*_S__f and the genomes of *Pf*_eH_ and *Pa*_T_. Thirty-eight percent of the *P. cerasus* proteins had a greater IAA to *Pf*_eH_, whereas 54% showed a greater IAA to *Pa*_T_. Eight percent showed an identical IAA to both ancestral species. A larger number of transcripts of both sour cherry subgenomes (Table 2) were assigned to the annotation data set of *Pf*_eH_. A total of 13,425 transcripts from the *Pce*_S__a subgenome and 13,107 from the *Pce*_S__f subgenome were found in the genome sequences of *Pf*_eH_ and *Pa*_T_. Seventy-five percent of the pool from the *Pce*_S__a subgenome showed a higher IAA to *Pa*_T_ and 17% to *Pf*_eH_, while 59% from the pool originating from the *Pce*_S__f subgenome showed a higher IAA to *Pf*_eH_ and 32% to *Pa*_T_.

**Table 2.** Comparison between the number of transcripts and %-IAA obtained from P. fruticosa ecotype Hármashatárhegy (Pf_eH_) and P. avium cv ‘Tieton’ (Pa_T_) representing the two ancestral species of P. cerasus

### Identification of syntenic regions and inversions

The sequences of the two subgenomes *Pce*_S__a and *Pce*_S__f and the genotypes *Pa*_T_ and *Pf*_eH_ of the two ancestral species *P. avium* and *P. fruticosa* were screened for duplicated regions using DAGchainer as previously published for peach (Verde et al. 2012). The seven major triplicated regions were found nearly one to one in *P. avium* but not in *P. fruticosa,* which lacked regions 4 and 7 corresponding to Verde et al. (2012). *P. avium* and *P. fruticosa* seem to derive from the same paleohexaploid event like peach, but with a loss of the fourth and seventh paleoset of paralogs in *P. fruticosa.* The graphical analysis is summarized in figure S7).

Thirteen inversions were detected through positional co-linearity comparison between the two subgenomes using the molecular markers from the 9+6k SNP array (Figure S8). Five inversions were found between subgenome *Pce*_S__a and the genome sequence of *Pa*_T_. Eleven inversions were found between sugenome *Pce*_S__f and *Pf*_eH_ (Table S6). By comparing the position of amino acid sequences of orthologous proteins (synteny), we found 21 inversions when comparing *Pce*_S__f with *Pf*_eH_. Only seven were found between *Pce*_S__a and *Pa*_T_ and 16 between both subgenomes *Pce*_S__a and *Pce*_S__f (Figure S9).

### Detection of de novo homoeologous exchanges

For the detection of de novo homoeologous exchanges we used three approaches by comparing inter- and intraspecific %-covered bases (genomic and transcriptomic) and %-IAA between proteins of *Pce*_S_ to *Pa*_T_ and *Pf*_eH_ (Figure 4, Figure S10, S11). *Pce*_S_ short-reads were mapped against *Pa*_T_ and *Pf*_eH_ and only species specific reads (*Pa*_T_ and *Pf*_eH_) were filtered into read-subsets. The obtained read-subsets were re-mapped against *Pce*_S__a and *Pce*_S__f and base coverage was calculated. A total of 1024 regions (100k window) were intraspecific %-covered bases from mapped reads (*Pce*_S__a to *Pa*_T_, *Pce*_S__f to *Pf*_eH_) was < than interspecific %-covered bases from mapped reads (*Pce*_S__a to *Pf*_eH_, *Pce*_S__f to *Pa*_T_) were discovered. In a second approach, translocations between the two subgenomes were localized by a short-read mapping analyses. Short-reads (RNAseq) from *P. cerasus, P. avium* and *P. fruticosa* obtained from Bird et al. (2022) were mapped on *Pce*_S_. A total of 148 regions were intraspecific difference of %-covered bases from obtained RNAseq reads (*Pa* and *Pce*_S__a, *Pf* and *Pce*_S__f) > than interspecific difference of %-covered bases from obtained RNAseq reads (*Pf* and *Pce*_S__a, *Pa* and *Pce*_S__f) indicated homoeologous exchanges between the two subgenomes. Finally, 367 regions were the proportion of transcripts with intraspecific amino acid identity (*Pa*_T_ and *Pce*_S__a, *Pf*_eH_ and *Pce*_S__f) < than the proportion of transcripts with interspecific amino acid identity (*Pf*_eH_ and *Pce*_S__a, *Pa*_T_ and *Pce*_S__f) were identified (Figure 4). Several regions were confirmed by calculating the 70% quantile of the IAA value within a window of 1 Mbp windows (Note S2). This confirms that there are transcripts in the *Pce_S_* genome whose IAA to the homoeologous representative genome (*Pf*_eH_) is greater than to the homologous (*Pa*_T_) representative. A total of 16 in *Pce*_S__a and 12 in *Pce*_S__f regions spanning 49 250k-windows were finally identified that match all three criteria indicating denovo homoelogous exchanges within the subgenomes (Figure 4, Figure S10). No evidence for an introgression of other *Prunus* species was found (Note S2, Note S3).

**Fig. 4.** Detected regions of homoeologous exchanges in the genome of *P. cerasus* ‘Schattenmorelle’. Circos plot of 16 pseudomolecules of the subgenomes of *Pce*_S__a and *Pce*_S__f. (a) chromosome length (Mb); (b) 16 in *Pce_S_*_a and 12 in *Pce*_S__f detected regions that match all three following analysis methods: (c) 1024 regions (100k window) were intraspecific %-covered bases from mapped reads (*Pce*_S__a to *Pa*_T_, *Pce*_S__f to *Pf*_eH_) was < than interspecific %-covered bases from mapped reads (*Pce*_S__a to *Pf*_eH_, *Pce*_S__f to *Pa*_T_); (d) 148 regions were intraspecific difference of %-covered bases from obtained RNAseq reads (*Pa* and *Pce*_S__a, *Pf* and *Pce*_S__f) > than interspecific difference of %-covered bases from obtained RNAseq reads (*Pf* and *Pce*_S__a, *Pa* and *Pce*_S__f); (e) 367 regions were the proportion of transcripts with intraspecific amino acid identity (*Pa*_T_ and *Pce*_S__a, *Pf*_eH_ and *Pce*_S__f) < than the proportion of transcripts with interspecific amino acid identity (*Pf*_eH_ and *Pce*_S__a, *Pa*_T_ and *Pce*_S__f).

### LTR dating and divergence of time estimation

Left and right LTR identity of a subset of 2,385 (*Pce*_S__a), 3,028 (*Pce*_S__f), 3,130 (*Pa*_T_) and 3,992 (*Pf*_eH_) LTRs was analysed. The homologous genomes shared 200 (*Pce*_S__a versus *Pa*_T_) and 100 (*Pce*_S__f versus *Pf*_eH_) LTRs whereas 12 LTRs were shared by *Pce*_S__a versus *Pf*_eH_ and *Pce*_S__f versus *Pa*_T_. Only five common LTRs were found between *Pce*_S__a and *Pce*_S__f and 13 between *Pa*_T_ and *Pf*_eH_. A summary of the LTRs insertion time is shown in figure 5A.

**Fig. 5.** Investigation on the evolution of the genome of *P. cerasus* ‘Schattenmorelle’. **A** Determination of insertion time from shared long terminal repeats (LTRs) in *P. cerasus* subgenome *avium* (*Pce*_S__a) and *P. cerasus* subgenome *fruticosa* (*Pce*_S__f) compared to *P. avium* ‘Tieton’ (*Pa*_T_) and *P. fruticosa* ecotype Hármashatárhegy (*Pf*_eH_). **B** Estimation of divergence of time (Mya) of *P. cerasus* subgenomes *Pce*_S__a and *Pce*_S__f compared to the donor species *P. avium* (Pa) and *P. fruticosa* (Pf). *Prunus yedonensis* (Pyn); *Prunus avium* (Pa); *Prunus persica* (Pp); *Prunus mume* (Pm); *Malus domestica* (Md); Paleocene (PAL); Eocene (EOC); Oligocene (OLI); Miocene (MIO); Pliocene (PlI); Pleistocene (PLEI).

The youngest shared LTRs between *Pce*_S__a and *Pa*_T_ were calculated with 103,896.1 generations and between *Pce*_S__f and *Pf*_eH_ with 97,402.6 generations. When comparing the homeologous chromosomes, the youngest shared LTRs between *Pa*_T_ and *Pf*_eH_ was calculated with 116,883.1 generations, between *Pce*_S__a and *Pce*_S__f with 194,805.2 generations. LTRs of *Pa*_T_ were also found in subgenome *Pce*_S__f and calculated with 207,792.2 generations. LTRs of *Pf*_eH_ were detected in subgenome of *Pce*_S__a and calculated with 149,350.6 generations. This indicates an exchange of LTRs between the two subgenomes.

A total of 834 single copy orthogroups among nine genomes were found and used for single protein alignments. Single alignments were concatenated and a final alignment with nine amino acid sequences representing each species with 419,586 amino acid positions was used for phylogenetic tree construction. Using the RelTime method, the estimated divergence time between the genera *Malus* and *Prunus* was 50.4 Mya. The species groups *P. persica* and *P. mume* diverged from the *P. yedonensis/P.avium/P.fruticosa* group 11.6 Mya. Based on this model, the divergence of the two subgenomes of *P. cerasus* compared to the genome sequences of *Pa*_T_ and *Pf*_eH_ was estimated with 2.93 Mya and 5.5 Mya respectively (Figure 5B).

## Discussion

The genome of the economically most important sour cherry ‘Schattenmorelle’ in Europe was sequenced using a combination of Oxford Nanopore R9.4.1 PromethION long read technology and Illumina NovaSeq^TM^ short read technology. After assignment of the long-reads to the two subgenomes and Hi-C analysis, the final assembly was 629 Mbp.

This sequence was used to study structural changes present in the allotetraploid sour cherry genome after its emergence. Therefore, the sour cherry genome sequence was compared to the published genome sequences of *Prunus avium* ‘Tieton’ (*Pa*_T_, Wang et al. 2019) and *Prunus fruticosa* ecotype Hármashatárhegy (*Pf*_eH_, Wöhner et al. 2021a) representing genotypes of the two ancestral species. The size of the subgenome *Pce*_S_ originating from *P. avium* was 269 Mbp. A similar genome size (271 Mbp) is described for the *Prunus avium* ‘Big Star’ (Pinosio et al. 2020) and ‘Sato Nishiki’ (Shirarsawa et al. 2017). Larger differences were found to ‘Regina’ with 279 Mbp and ‘Tieton’ with 344 Mbp (Wang et al. 2020, Le Dantec et al. 2019). Differences were also found between the size of subgenome *Pce*_S__f (299 Mbp) and the genome of the ground cherry genotype *P. fruticosa* ecotype Hármashatárhegy (366 Mbp, Wöhner et la. 2021a). These differences indicate a reduction of the subgenome *Pce*_S__a by 0.49%-21%, whereas for subgenome *Pce*_S__f a reduction of 18.29% was found. The reduction in genome size for allotetraploid species in comparison to their ancestral genomes is common, as already reported for *Nicotiana tabacum* (1.9%-14.3%) and *Gossypium* species (Leitch et al., 2008; Hawkins et al., 2009; Renny-Byfield et al., 2011), with an overall downsizing rate for angiosperms calculated as 0%-30% (Zenil-Ferguson et al. 2016). Genome downsizing in repose to a genome hybridisation event can be explained with evolutionary advantages, which give these species with smaller genomes a selection advantage in the long-term (Knight et al. 2005, Zenil-Ferguson et al. 2016).

Although downsizing of the *P. cerasus* subgenomes is most probable, enlargement and expansion of the genomes of ancestral species during evolution would be another possibility. However, an increase in genome size during the evolution of a species has only rarely been documented (Leitch et al. 2008, Jackobs et al. 2004, Kim et al. 2014). BUSCO analysis provides additional evidence for the reduction of genome size. Although the number of genes does not correlate with genome size in eukaryotes (Pierce 2012), differences between the ancestral genomes and sour cherry could be observed when looking at BUSCO-completeness. Considering both subgenomes and the genomes of *Pa*_T_ and *Pf*_eH_, a completeness of > 96.4% was obtained. However, the completeness of the single subgenomes was only 89.4% for *Pce*_S__a and 87.1% for *Pce*_S__f. Structural differences between the *P. cerasus* subgenomes and the genomes of *Pa*_T_ and *Pf*_eH_ were also found by comparing the number of repetitive elements. While the content of repetitive elements differs by only 0.86% between *Pf*_eH_ and *Pce*_S__f, it is 17.8% between *Pa*_T_ and *Pce*_S__a. Whether this is a consequence of hybridisation remains speculative and would deserve further studies. An increase of class I elements Gypsy from six percent in the *Pa*_T_ genome to 7.3% in subgenome *Pce*_S__a indicates an expansion of this class following the formation of the sour cherry genome or a possible reduction of non-repetitive sequences in the corresponding subgenome resulting in a smaller genome size.

A comparison of syntenic regions showed a high degree of collinearity between *Pce*_S_, *Pa*_T_ and *Pf*_eH_ genomes (Figure 2, Figure S7-9), with single inversions between the respective chromosome pairs. Using the genome of *P. persica,* seven triplicated regions were detected in *Pce*_S__a, confirming that these genomes descend from a palaeohexploid ancestor. However, the triplicated regions 4 and 7 in *Pce*_S__f were only detected in highly fragmented form or have been lost. Hao et al. (2022), who described a rapid loss of homoeologs immediately after polyploidy events, described similar finding.

Based on Ranallo-Benavidez (2020) the results from the k-mer analysis confirm that the genome of sour cherry can be considered as highly heterozygous and segmental allotetraploid. Furthermore, genomes of segmental allopolyploids may possess a mix of auto- and allopolyploid segments through duplication-deletion events as a result of homoeologous exchanges leading to either reciprocal translocations or homoeologous non-reciprocal translocations (Mason and Wendel, 2020). Whereas autotetraploids have an aaab > aabb rate, allotetraploids are considered to have aaab < aabb. The near identical rate between aaab and aabb in *Pce*_S_ provides strong evidence that the sour cherry is a segmental mix of auto- and allotetraploidy.

Due to this assumption of segmental allotetraploidy, homoeologous recombination between homoeologous chromosomes is very likely. This is confirmed by the coverage and amino acid identity analyses. Homoeologous exchange events between the chromosomes of subgenome *Pce*_S__a and *Pce*_S__f were detected (Figure 4, Figure S10). These exchanges are not balanced but probably a product of a duplication/deletion event as described by Mason et al. (2020), generating the proposed mosaic of genomic regions representing one or the other subgenome.

Additional mapping of transcriptomes from six other *Prunus* species indicates that no major introgression from one of these species occurred. Using 14 reference species, 60,123 gene models were annotated. Almost the same number was assigned to the two *P. cerasus* subgenomes (Table 2). No evidence was found for large introgressions from any of the reference species (Note S2, S3). By comparing the amino acid identity of the proteins of *Pa*_T_ and *Pf*_eH_ with the respective sour cherry subgenome, the identified translocations via read mapping could be confirmed. The majority of the transcripts (51%) could be assigned to the genotypes *Pa*_T_ and *Pf*_eH_ of the two ancestral species *P. avium* and *P. fruticosa* (Figure S6). 5.2% of the transcripts could not be assigned to any of the reference species. Only < 1% of the transcripts could not be assigned to one of the ancestral species. They showed equivalent matches to both species and are probably a product of ab-initio prediction. A total of 49,698 proteins in subgenome *Pce*_S__a and 48,576 proteins in *Pce*_S__f shared only 13,435 and 13,107 proteins with *Pa*_T_ and *Pf*_eH_, respectively. A total of 75% of the proteins of subgenome *Pce*_S__a matched better to *Pa*_T_ compared to *Pf*_eH_, whereas only 59% of *Pce*_S__f mapped better to *Pf*_eH_ than to *Pa*_T_ (Table 2). Subgenome *Pce*_S__a seems to be closer to *Pa*_T_ than *Pce*_S__f to *Pf*_eH_. This was confirmed by an evolutionary approach that calculated the separation of the subgenome *Pce*_S__f from *P. fruticosa* 5.5 Mya, while subgenome *Pce*_S__a separated from *P. avium* 2.93 Mya (Figure 5B).

To validate these results, the insertion events of long terminal repeats between *P. avium, P. fruticosa* and the subgenomes were calculated. Assuming a *Prunus*-specific rate of 7.7*10^-9^ mutations per generation (Xie et al. 2016), LTRs of the same type with the same insertion time were identified in the same positional order in the different (sub)genomes. The most recent co-occurring LTRs between the genomes of *Pa*_T_ and *Pf*_eH_ could be dated at 116,883.1 generations. Exact data on the duration of the generation time of *Prunus* species in natural habitats do not exist. Although the juvenile phase of many *Prunus* species is usually completed after 5 years (Besford et al. 1996), it can be assumed that the times for a generation are considerably higher. Many fruit species are hardly or not at all able to rejuvenate by seeds under natural conditions (Coart et al. 2003), or they rejuvenate mainly by root suckers (Li et al. 2022). Other studies on *Prunus* therefore assume a duration of 10 years per generation (Wang et al. 2021), although even this seems rather too little.

Assuming that a generation change is to be expected after 10 to 60 years (Besford et al. 1996), this would correspond to a time period of ~ 1 to 6 Mya. The youngest co-occurring LTR could be estimated at 1.9 Mya. This suggests that *P. fruticosa* and *P. avium* probably shared a gene pool between ~1 Mya and 2 Mya. It should be noted that this estimate can vary greatly depending on the number of years per generation used in the calculation (Figure 5A). Based on the results of the protein dating a generation time of 30 years is more likely for *P. avium*. For *P. fruticosa*, which occurs less frequently in natural habitats and reproduces mainly via root suckers, the generation time seems to be somewhat longer at 55 to 60 years. Some LTRs present in *Pa*_T_, but absent in *Pf*_eH_ were found in subgenome *Pce*_S__f only and vice versa. Other class I elements (LTR - ERV1, Pao) and Academ/-2 specifically detected in one of the two genotypes *Pa*_T_ and *Pf*_eH_ representing the two ancestral species of sour cherry and in *Pce*_S__a and *Pce*_S__f, which indicates a transfer of these elements between the two subgenomes following the formation of the allotetraploid *P. cerasus* genome. This is a further indication for a segmental exchange between the two sour cherry subgenomes. Mason et al. (2020) speculated that uni-directional homeologous exchange was observed in recent or synthetic allopolyploids. However, our results confirm this hypothesis by the evidence that sour cherry is a recent allopolyploid with autopolyploid segments derived from uni-directional homoeologous exchanges.

## Conclusion

After sequencing of the genome of the sour cherry ‘Schattenmorelle’, the following can be concluded. The genome of sour cherry is segmental allotetrayploid. It consists of two subgenomes, one derived from the sweet cherry *P. avium* and one from the ground cherry *P. fruticosa*. DNA sequences have been repeatedly exchanged between the two subgenomes. At the same time, a reduction in genome size has taken place. Other *Prunus* species have not contributed to the evolution of this species. No evidence was found for introgressions in the sour cherry genome derived from *Prunus* species other than *P. avium* and *P. fruticosa*. Sour cherry is a very young *Prunus* species. The origin of this species is estimated about 1 mya earliest.

## Material and Methods

### Plant Material, DNA and RNA extraction, sequencing and iso-seq analysis

Snap frozen *Prunus cerasus* L. ‘Schattenmorelle’ (accession KIZC99-2, Figure 1, supplements 1.1) young leaf material was sent to KeyGene N.V. (Wageningen, The Netherlands). High molecular weight extracted DNA (Wöhner et al. 2021a) was used to generate 1D ligation (SQK-LSK109) libraries which were subsequently sequenced on two Oxford Nanopore Technologies (ONT) R9.4.1 PromethION flow cells. The same material was used to generate an Illumina PCR free paired-end library (insert size of ~550 bp) which was sequenced on a HiSeq 4000^TM^ platform using 150bp and 125bp paired-end sequencing.

Snap frozen tissues from buds, flowers, leaves and fruits were collected and in the field and total RNA was extracted with Maxwell® RSC Plant RNA Kit (Promega). Two pools were generated and used for PacBio Iso-Seq library preparation (Procedure & Checklist – Iso-Seq™ Express Template Preparation for Sequel® and Sequel II Systems, PN 101-763-800 Version 02). Each library pool was sequenced on a single 8M ZMW PacBio Sequel II SMRT cell (supplements 1.1). Obtained full-length reads with 5’-end primer, the 3’-end primer and the poly-A tail were filtered and these sequences were trimmed off. Transcripts containing (artificial) concatemers were completely discarded. Isoforms (consensus sequence) generated by full-length read clustering (based on sequence similarity), were finally polished with non-full-length reads using Arrow (SMRT Link v7.0.0, https://www.pacb.com/wp-content/uploads/SMRT_Tools_Reference_Guide_v600.pdf).

### De novo assembly and scaffolding

The aligner Minimap2 (v2.16-r922, Li et al. 2018) and assembler Miniasm (v0.2-r137-dirty, Li et al. 2016) were used for raw data assembly generation. Racon (vv1.4.10, Vaser et al. 2017) and Pilon (v1.22, Walker et al. 2014) were used for base-quality improvement with raw ONT and Illumina read data. Chromosome-scale scaffolding was performed by Phase Genomics (Seattle, Washington, USA) with Proximo Hi-C (supplements 1.2). The resulting assembly was designated as 20-WGS-PCE_<Avium|Fruticosa>.2.0 _<Contig|Scaffold>.

*Correctness, completeness and contiguity of the Prunus cerasus genome sequence* The BUSCO (Benchmark Universal Single-Copy Orthologs - Galaxy Version 4.1.4) software was used for quantitative and quality assessment of the genome assemblies based on near-universal single-copy orthologs. The long terminal repeat (LTR) assembly Index (LAI) Ou et al. (2018) was calculated with LTR_retriever 2.9.0 (https://github.com/oushujun/LTR_retriever) to evaluate the assembly continuity between the final genome sequence of *P. cerasus* ‘Schattenmorelle’ and *Prunus fruticosa* ecotype Hármashatárhegy (Wöhner et al. 2021a, Pf_1.0), *P. avium* ‘Tieton’ (Wang et al. 2020) and *P. persica* ‘Lovell’ (Verde et al. 2017) respectively. LTR_harvest (genometools 1.6.1 implementation) was used to obtain LTR-RT candidates. The genome size was also estimated by k-mer analysis (supplements 1.3) using Illumina short read data. The merged datasets were subsequently used to generate a histogram dataset representing the k-mers from all datasets. GenomeScope (Galaxy Version 2.0, Ranallo-Benavidez 2020) was used to generate a histogram plot of k-mer frequency of different coverage depths using the tetraploid ploidy level (k-mer length 19). Marker sequences and genetic positions from five available genetic sour cherry maps (M172x25-F1, US-F1, 25x25-F1, Montx25-F1, RE-F1) and 14,644 SNP markers (9+6k array) were downloaded from the Genome Database for Rosaceae (GDR, https://www.rosaceae.org/). The marker sequences were mapped on the chromosome sequences using the mapping software bowtie2 (Galaxy Version 2.5.0+galaxy0, Langmead et al. 2012) implementation on the Galaxy server (https://usegalaxy.org) with standard settings.

### Structural and functional annotation

For an inter species repeat comparison, a species-specific repeat library was generated with RepeatModeler open-1.0.11, and the genome was subsequently masked with RepeatMasker open-4.0.7. For structural genome annotation, another species-specific repeat library for PCE_1.0 was generated with RepeatModeler2 (Flynn et al., 2020) version 2.0.2, and the genome was subsequently masked with RepeatMasker 4.1.2. (Further details on the repeat masking software configuration are available in supplements 1.4.1.).

To generate extrinsic evidence for structural annotation of protein coding genes, short read RNA-Seq library SRR2290965 was aligned to the genome using HiSat2 version 2.1.0 (Kim et al., 2019). The output SAM file was converted to BAM format using SAMtools (Li et al, 2009). The resulting alignment file was further used by both BRAKER1 (Hoff et al., 2016; Hoff et al., 2019) and GeMoMa (Keilwagen et al. 2016). Further, a custom protocol was used for integrating long read RNA-Seq data into genome annotation (supplements 1.4.2). In short, protein coding genes were called in Cupcake transcripts using GeneMarkS-T (Tang et al., 2015) and these predictions were converted to hints for BRAKER1. In addition, intron coverage information from long read to genome spliced alignment with Minimap2 (Li et al. 2018) was provided to BRAKER1.

A combination of BRAKER1 (Hoff et al., 2016, Hoff et al., 2019), BRAKER2 (Bruna et al., 2021), TSEBRA (Gabriel et al., 2021), and GeMoMa (Keilwagen et al. 2016) was used for the final annotation of protein coding genes. BRAKER pipelines use a combination of evidence-supported self-training GeneMark-ET/EP (Lomsadze et al., 2014; Bruna et al., 2020) (here version 4.68) to generate a training gene set for the gene prediction tool AUGUSTUS (Stanke et al., 2008; here version 3.3.2). BRAKER1 version 2.1.6 was here provided with BAM-files of from short and long read RNA-Seq to genome alignments, and with gene structure information derived from Cupcake transcripts using GeneMarkS-T. This generated a gene set that consists of ab initio and evidence supported predictions. A separate gene set was generated with BRAKER2, which uses protein to generate a gene set. We used the OrthoDB version 10 (Kriventseva, 2019) partion of plants in combination with the full protein sets of *Prunus fruticosa* (Wöhner et al., 2021b), *Prunus armeniaca* (GCA 903112645.1), *Prunus avium* (GCF_002207925.1), *Prunus dulcis* (GCF_902201215.1), *Prunus mume* (GCF_000346735.1), *Prunus persica* (GCF_000346465.2) as input for BRAKER2. Both the BRAKER1 and BRAKER2 AUGUSTUS gene sets were combined with a GeneMarkS-T derived gene set using TSEBRA (Gabriel et al., 2021) from the long_reads branch on GitHub with a custom configuration file (supplements 1.4.3.) incorporating evidence from BRAKER1 and BRAKER2.

GeMoMa was run on the genome assembly of Schattenmorelle using 14 reference species and experimental transcript evidence (supplements 1.4.4). GeMoMa gene predictions of each reference species were combined with TSEBRA predictions using the GeMoMa module GAF and subsequently, UTRs were predicted in a two-step process based on mapped Iso-seq and RNA-seq data using the GeMoMa module AnnotationFinalizer (supplements 1.4.5). First, UTRs were predicted based on Iso-seq data. Second, UTRs were predicted based on RNA-seq data for gene predictions without UTR prediction from the first step. An assembly hub for visualization of the *Prunus cerasus* genome with structural annotation was generated using MakeHub (Hoff, 2019; supplements 2). The functional annotation was performed with the Galaxy Europe implementation of InterProScan (Galaxy Version 5.59-91.0+galaxy3, Jones et al. 2014, Cock et al 2013, Zdobnov et al. 2001, Quevillon et al. 2005, Hunter et al. 2009). The chloroplast and mitochondria sequences were annotated with GeSeq (Tillich et al. 2017, supplements 1.4.5).

### Identification of syntenic regions

Structural comparison of orthologous loci between the subgenomes *Pce*_S__a and *Pce*_S__f of *Prunus cerasus* and the two genotypes *Pa*_T_ and *Pf*_eH_ as representatives of the two genome donor species *P. avium* and *P. fruticosa* was calculated with the final annotations using SynMap2 (Haug-Baltzell et al. 2017) available at the CoGe platform (https://genomevolution.org/coge/). Analysis on triplication events were performed with standard settings and Last as Blast algorithm at a ratio coverage depth 3:3 in SynMap2 (Haug-Baltzell et al. 2017).

### Identification of homoeologous exchange regions

Homoeologous exchanges were identified on the amino acid, transcript and genomic level.

### Calculation of amino acid identity

Identity of amino acids (IAA) between all reference annotations homology-based gene prediction was calculated by GeMoMa using the default parameters. Subsequently, the *Pce*_S_ genome was divided into 250k windows, and the percentage of proteins showing a higher IAA between *Pf_eH_* (Wöhner et al. 2021b) and *Pa_T_* (Wang et al. 2020) to the respective subgenome (*Pce*_S__a and *Pce*_S__f) was determined. The percentage of proteins in this window, which were more similar to *Pa*_T_ was finally subtracted from the percentage of proteins which were more similar to *Pf*_eH_. A proportion of transcripts with higher intraspecific amino acid identity (between *Pa_T_* and *Pce_S_*_a or *Pf_eH_* and *Pce_S_*_f) is expected compared to the proportion of transcripts with interspecific amino acid identity match (*Pf_eH_* and *Pce_S_*_a or *Pa_T_* and *Pce_S_*_f). Opposite cases, indicate potential translocations between the two subgenomes *Pce*_S__a and *Pce*_S__f and were plotted into a circus plot (Figure S11).

### Read mapping and coverage analysis

RNAseq raw data published by Bird et al. (2022) was obtained from NCBI sequence read archive (SRA) for the following species: *P. cerasus* (SRX14816146, SRX14816142, SRX14816138), *P. fruticosa* (SRX14816141), *P. avium* (SRX14816143), *P. canescens* (SRX14816137), *P. serrulata* (SRX14816136), *P. mahaleb* (SRX14816140), *P. pensylvanica* (SRX14816144), *P. maackii* (SRX14816139), *P. subhirtella* (SRX14816145). Reads were adapter- and quality trimmed using the software Trim Galore (version 0.6.3, parameters --quality 30 –length 50). Trimmed reads were mapped against the *P. cerasus* subgenomes *Pce*_S__a and *Pce*_S__f using STAR (version 2.7.8a, parameter --twopassMode Basic). The subsequent analysis was performed in accordance to Keilwagen et al. (2022). The *Pce*_S_ genome was divided into 250k windows. The percentage of covered bases using RNAseq data of *P. cerasus* (SRX14816146, SRX14816142, SRX14816138) was estimated at a depth of 1 for each window. The same was done with all other RNAseq data sets. The percentage of covered bases from *P. avium* (SRX14816143) was subtracted from the percentage of covered bases from *P. cerasus* (SRX14816146, SRX14816142, SRX14816138). The same was done using the reads of *P. fruticosa* (SRX14816141). For subgenome *Pce*_S__a it is expected that the intraspecific difference for transcripts of data set *P. avium* (SRX14816143) is lower (close to 0) than the interspecific difference for transcripts of data set *P. fruticosa* (SRX14816141) and vice versa. Opposite cases indicate potential homoeologous exchanges between the two subgenomes *Pce*_S__a and *Pce*_S__f and were plotted into a circos plot (Figure S11).

The nucleotide short reads from *Pce_S_* were mapped against the genomes of the two ancestral species *Pa*_T_ and *Pf*_eH._ Subsequently, the mapped reads were filtered using samtools for mapped reads in proper pair (-f 3) and primary alignments and not supplementary alignment (-F 2304). Those reads were divided into four groups according to the following criteria: 1. unique match to *Pa*_T_: 2. unique match to *Pf*_eH_; 3. match to *Pa*_T_ and *Pf*_eH_; 4. no match to *Pa*_T_ and *Pf*_eH_ (unique to *Pce_S_*). The first two separated read sets were then re-mapped against the subgenomes *Pce*_S__a and *Pce*_S__f. The percentage of covered bases was calculated for a 100 k window. For the subgenomes of *Pce*_s_, the percentage of intraspecific covered bases (*Pce_S_*_a to *Pa_T_*, *Pce_S_*_f to *Pf_eH_*) should be higher compared to the percentage of interspecific covered bases (*Pce_S_*_a to *Pf_eH_*, *Pce_S_*_f to *Pa_T_*). The opposite case indicates possible translocations and were plotted into a circus plot (Figure S11). Additionally, regions of the Schattenmorelle genome assembly were determined that are uniquely covered by *Pa*_T_ and *Pf*_eH_ filtered read sets.

### LTR insertion estimation

The difference (identity) of left and right LTR was calculated using the script EDTA_raw.pl from the software EDTA version 1.9 (https://github.com/oushujun/EDTA, Ou et al. 2019). As input files we used the genome sequences of *P. cerasus* (*Pce*_S__a and *Pce*_S__f), *Pa*_T_ (NCBI BioProject acc. no. PRJNA596862), *Pf*_eH_ (NCBI BioProject acc. no. PRJNA727075) and a curated library of representative transposable elements from *Viridiplantae* (https://www.girinst.org/repbase/). Because trees are not annual plants, the identity obtained from the resulting .pass.list file was used for the estimation of generation time after LTR insertion using the formula T=K/2µ (K is the divergence of the LTR = 1 – identity) assuming a *Prunus* specific mutation frequency of µ=7.7 * 10^-9^ (Xie et al. 2016) per generation.

### Protein clustering, multiple sequence alignment and divergence of time estimation

The protein datasets from *Pce*_S_*_a* and *Pce*_S_*_f, Pa*_T_, *Pf*_eH_, *Pp* (*Prunus persica* Whole Genome Assembly v2.0, v2.0.a1), *Pm* (*Prunus mume* Tortuosa Genome v1.0), *Py* (*Prunus yedoensis* var. *nudiflora* Genome v1.0), *Md* (*Malus* x *domestica* HFTH1 Whole Genome v1.0) and *At* (TAIR10.1, RefSeq GCF_000001735.4) from the annotation step were uploaded to Galaxy_Europe server as .fasta. The Proteinortho (Galaxy Version 6.0.32+galaxy0) was used to find orthologous proteins within the datasets. MAFFT (Galaxy Version 7.505+galaxy0) was used to align the obtained single copy orthogroups. The final alignments were merged with the Merge.files function (Galaxy Version 1.39.5.0). Finally, the alignments were concatenated into a super protein and the final sequences were aligned with MAFFT. A phyogenetic tree was reconstructed with RAxML (maximum likelihood based inference of large phylogenetic trees, Galaxy Version 8.2.4+galaxy3) and the obtained .nhx file was reformatted as .nwk file for further processing using CLC Mainworkbench (21.0.1, QIAGEN Aarhus A/S). Evolutionary analyses were conducted in MEGA X (Kumar et al. 2018). Estimation of pairwise divergence time was performed according to Shirasawa et al. (2019) with a divergence time from the reference species peach and apple (34-67 Mya, www.timetree.org). Specific parameters for the calculation are listed in supplements.

## Data Availability

Data supporting the findings of this study are deposited into the Open Agrar repository (https://doi.org/10.5073/20230324-105730-0, Wöhner et al. (2023)) and on personal request to the corresponding author. An assembly hub for genome and annotation visualization is permanently hosted at http://bioinf.uni-greifswald.de/private-hubs/pcer/hub.txt.

## Author contributions

TWW, OFE, HF wrote the manuscript. AHJW, KN, RPW and EJB performed DNA isolation, sequencing, genome assembly and scaffolding. KJH, JK, LG, HT and TB performed masking and annotation of the dataset. OA and LB performed the LTR-dating and TW did the corresponding analysis and results compilation. JK performed BUSCO analysis, read mapping and coverage analysis. TW the did the k-mer analysis, interproscan, synteny analysis, phylogenetic analysis and circus plots. TW and JL performed the Lai-Index analysis. HF, MS and AP conceived the study and made substantial contributions to its design, acquisition, analysis and interpretation of data. All authors contributed equally to the finalization of the manuscript.

## Acknowledgments

We would acknowledge the Galaxy Europe, Galaxy USA server administration for support and provision of resources. Special thanks to Eric Lyon for support with Synmap2.

## Supplemental information (SI)

Document S1 - Supplemental material and methods

## Supplemental Figures

**Figure S1** Hi-C heatmap post-scaffolding for the subgenomes *Pce*_S__a (left) and *Pce*_S__f (right) of *P. cerasus* cv ‘Schattenmorelle’. The heatmap indicates the density of paired Hi-C reads which interact to each other in close proximity. High intense colour indicates high interaction.

**Figure S2** The chloroplast (a) and mitochondrial (b) sequences of *P. cerasus* L. cv ‘Schattenmorelle’.

**Figure S3** Analysis of completeness of the *P. cerasus* cv. Schattenmorelle subgenomes *P. cerasus* cv ‘Schattenmorelle’ subgenome *avium* (*Pce*_S__a) and *P. cerasus* cv ‘Schattenmorelle’ subgenome *fruticosa* (*Pce*_S__f) and combined datasets compared to *P. avium* cv. ‘Tieton’ (*Pa*_T_) and *P. persica* cv. Lovell (*Pp*_L_) by mapping of a set of universal single-copy orthologs using BUSCO. The bar charts indicate complete **<u>s</u>**ingle copy (orange), complete **<u>d</u>**uplicated (gray), **f**ragmented (yellow) and **<u>m</u>**issing (blue) genes. For evaluation the embryophyta_odb10 BUSCO dataset (n=1614) was used. *P. cerasus* cv. Schattenmorelle show a 99 % completeness (S: 16.7 %, D: 82.3 %, F: 0.4 %, M: 0.6 %, n: 1614) which reaches the completeness of *P. avium* cv. ‘Tieton’ (C: 98.3 %, S: 95.6 %, D: 2.7 %, F: 0.5 %, M:1.5 %, n:1614) and *P. persica* ‘Lovell’ (C: 99.3 %, S: 97.5 %, D: 1.8%, F: 0.1 %, M: 0.6 %, n:1614).

**Figure S4** Assessing the quality of repetitive sequences between the chromosome sequences of *P. avium* ‘Tieton’ (A), *P. fruticosa* ecotype Hármashatárhegy (B), *P. perisca* ‘Lovell’, and (C) *P. cerasus* subgenome *avium* and *P. cerasus* subgenome *fruticosa* using the LAI index. The genomes *P. cerasus* [this study] and *P. avium* were sequenced with ONT 9.4.1 and Illumina (Wang wet al. 2020), *P. fruticosa* with ONT 10.3 (Wöhner et al. 2021) and *P. persica* with Illumina and Sanger sequencing of fosmid and BAC clones (Verde et al. 2017).

**Figure S5** Collinearity plots between the five published genetic maps of sour cherry (M172x25-F1, US-F1, 25x25-F1, Montx25-F1, RE-F1) and the *P. cerasus* cv ‘Schattenmorelle’ subgenome *avium* (*Pce*_S__a) and *fruticosa* (*Pce*_S__f).X-axis represents the genetic position of a marker in the genetic linkage map given in centi Morgan (cM). Y-axis represents the physical position of a marker sequence within the genome sequence of the respective subgenome given in Mega base pairs (Mbp).

**Figure S6** Percentage of *P. cerasus* (*Pce*) proteins by IAA compared with 15 reference species. *P. fruticosa* ecotype Hármashatárhegy (*P_f_*_eH_), *P. avium* ‘Tieton’ (*Pa*_T_), *P. yedonensis* (*Pyed*), *P. domestica* (*Pd*), *P. armeniaca* (*Par*), *P. persica* (*Pp*), *Pyrus communis* (*Pyrco*), *Populus trichocarpa* (*Poptri*), *Vitis vinifera* (*Vv*), *Arabidopsis thaliana* (*At*), *Malus domestica* (*Md*).

**Figure S7** Synmap2 plots of self-self comparisons between (A) *Prunus persica* ‘Lovell’ (*Pp*_L_) and *P. avium* ‘Sato Nishiki’ (*Pa*_S_), (B) *P. fruticosa* ‘Hármashatárhegy’ (*Pf*_eH_), (C) *P. cerasus_avium* ‘Schattenmorelle’ (*Pce*_S__a), (D) *P. cerasus_fruticosa* ‘Schattenmorelle’ (*Pce*_S__f) for the identification of triplicated regions (TR) 1-7.

**Figure S8** Positional co-linearity comparison between the two subgenomes *P. cerasus_avium* ‘Schattenmorelle’ (*Pce*_S__a), *P. cerasus_fruticosa* ‘Schattenmorelle’ (*Pce*_S__f) and *P. avium* ‘Tieton’ (*Pa*_T_), *P. fruticosa* ‘Hármashatárhegy’ (*Pf*_eH_), using the molecular markers from the 9+6k SNP array. The plots were generated using the R-software package chromoMap v0.4.1.

**Figure S9** Synteny between (a) *P. cerasus* subgenome _*avium* (*Pce*_S__a) and *P. cerasus* subgenome _*fruticosa* (*Pce*_S__f), and (b) the subgenomes and the genotypes *Prunus avium* ‘Tieton’ (*Pa*_T_) and *P. fruticosa* (*Pf*_eH_) of the ancestral species *P. avium* and *P. fruticosa.* Blue arrows indicate positions where inversions occurred.

**Figure S10** Detected regions of homoeologous exchanges in the genome of *P. cerasus* ‘Schattenmorelle’. Circos plot of 16 single pseudomolecules 1 to 8 (see continuating plots) of the subgenomes of *Pce*_S__a and *Pce*_S__f. (a) chromosome length (Mb); (b) region in *Pce_S_*_a and in *Pce*_S__f detected that match all three following analysis methods: (c) regions (100k window) were intraspecific %-covered bases from mapped reads (*Pce*_S__a to *Pa*_T_, *Pce*_S__f to *Pf*_eH_) was < than interspecific %-covered bases from mapped reads (*Pce*_S__a to *Pf*_eH_, *Pce*_S__f to *Pa*_T_); (d) regions were intraspecific difference of %-covered bases from obtained RNAseq reads (*Pa* and *Pce*_S__a, *Pf* and *Pce*_S__f) > than interspecific difference of %-covered bases from obtained RNAseq reads (*Pf* and *Pce*_S__a, *Pa* and *Pce*_S__f); (e) regions were the proportion of transcripts with intraspecific amino acid identity (*Pa*_T_ and *Pce*_S__a, *Pf*_eH_ and *Pce*_S__f) < than the proportion of transcripts with interspecific amino acid identity (*Pf*_eH_ and *Pce*_S__a, *Pa*_T_ and *Pce*_S__f).

## Supplemental Tables

**Table S1** Statistics of different assemblies for *P. cerasus* cv ‘Schattenmorelle’ (*Pce*_S_) and the subgenomes *Pce*_S__a and *Pce*_S__f

**Table S2** Pseudomolecule statistics for *Pce*_S_

**Table S3** Iso-Seq results

**Table S4** Functional annotation results generated by interproscan using BRAKER & GeMoMa combination of ab-initio and homology-based structural gene annotation and statistics

**Table S5** Mapping of marker sequences to *Prunus cerasus* ‘Schattenmorelle’ (*Pce*_S_) genome

**Table S6** Position of inversions within the subgenomes of *Pce*_S_ indicated by collinearity of marker positions.

## Supplemental Notes

**Note S1** Access to the assembly hub for genome and annotation visualization at UCSC Genome Browser.

**Note S2** Calculation of the 70% quantile from IAA.

**Note S3** Calculation of %-covered bases obtained from RNAseq data.

