## supplements for "The structure of the tetraploid sour cherry ‘Schattenmorelle’ (*Prunus cerasus* L.) genome reveals insights into its segmental allopolyploid nature"

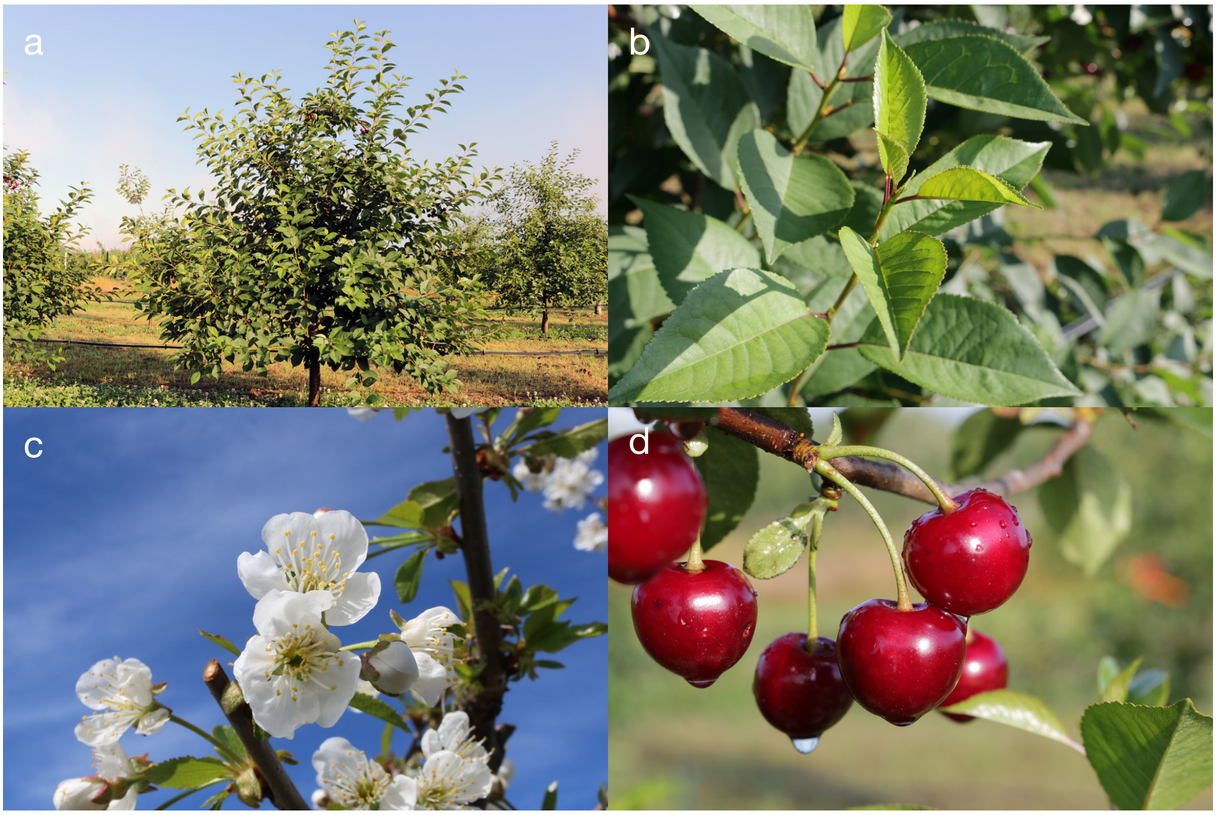


**Figure 1** Morphology of *P. cerasus* L. 'Schattenmorelle'. (a) mature tree habitus, (b) leaves, (c) inflorescence, (d) fruits.

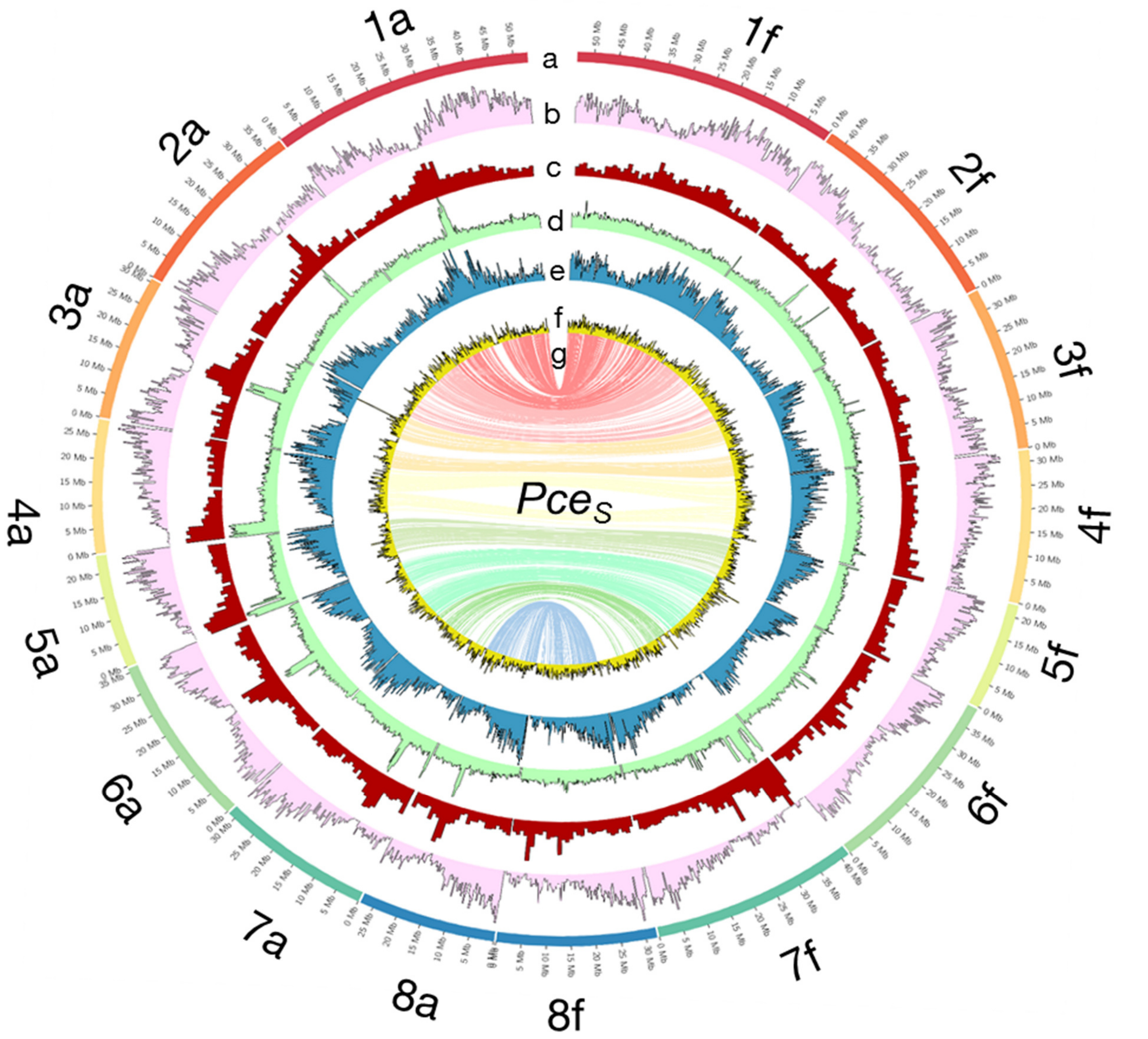


**Figure 2** The genome of *P.cerasus* 'Schattenmorelle'. Circos plot of 16 pseudomolecules of the subgenomes of *Pce*$_S$_a and *Pce*$_S$_f. (a) chromosome length (Mb); (b) gene density in blocks of 250k: (c) distribution of repetitive sequences in blocks of 250k (d) Gypsy elements in blocks of 250k; (e) Copia elements in block of 250k; (f) GC content in blocks of 1 Mb. (g) The inner ring shows markers from the 6+9k SNP array located on both subgenomes.

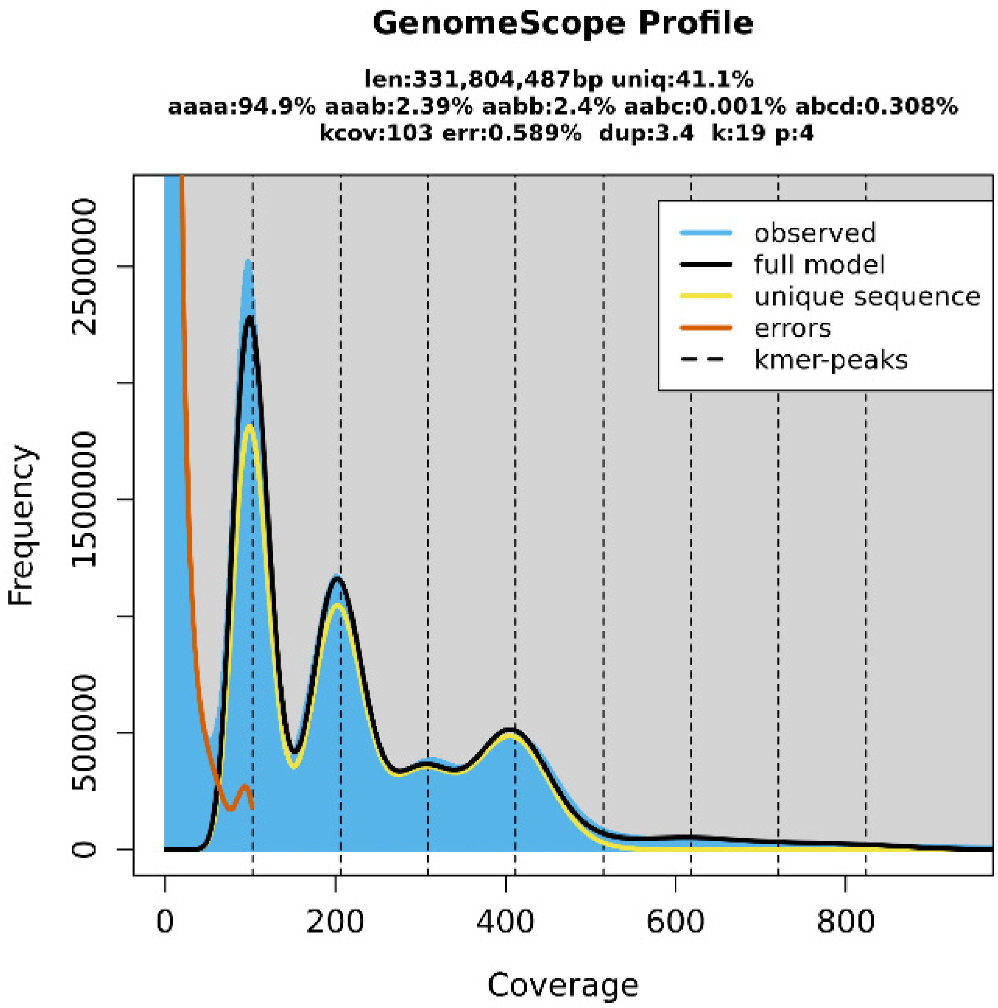


**Figure 3** GenomeScope (Galaxy Version 2.0) estimation of the *P. cerasus* genome size by k-mer counts obtained from the software Meryl (Galaxy Version 1.3+galaxy2). Both programs are integrated on the GalaxyServerEurope. The k-mer-peaks indicate that k-mers with a length of 19 bp occur in heterozygote (100x depth, 200x depth, 300x depth) and homozygote (400x depth) constitution within the genome. Coverage depth of individual k-mers is assigned as coverage.

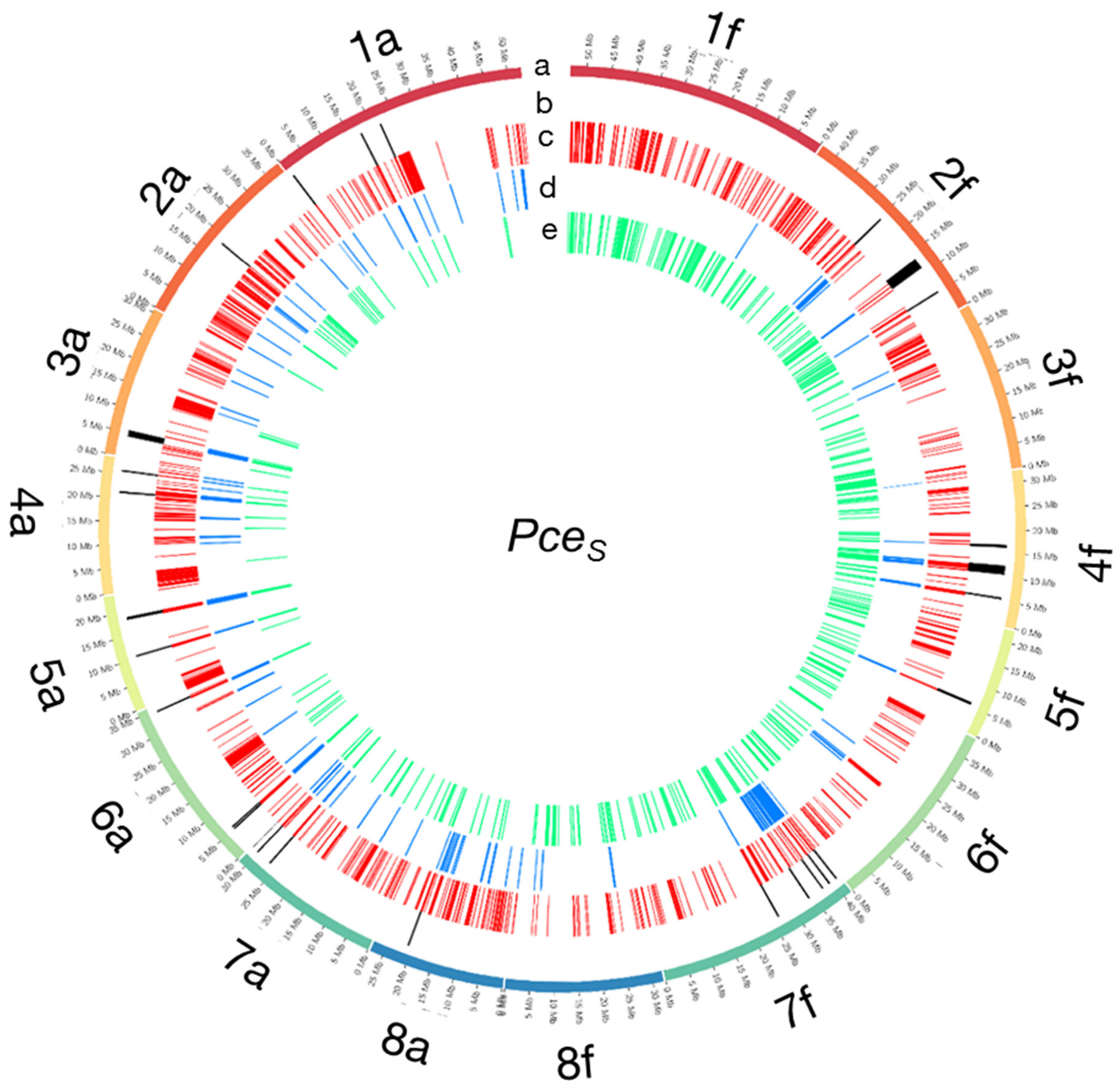


**Figure 4** Detected regions of homoeologous exchanges in the genome of *P. cerasus* 'Schattenmorelle'. Circos plot of 16 pseudomolecules of the subgenomes of $Pce_S$_a and $Pce_S$_f. (a) chromosome length (Mb); (b) 16 in $Pce_S$_a and 12 in $Pce_S$_f detected regions that match all three following analysis methods: (c) 1024 regions (100k window) were intraspecific %-covered bases from mapped reads ($Pce_S$_a to $Pa_T$, $Pce_S$_f to $Pf_{eH}$) was < than interspecific %-covered bases from mapped reads ($Pce_S$_a to $Pf_{eH}$, $Pce_S$_f to $Pa_T$); (d) 148 regions were intraspecific difference of %-covered bases from obtained RNAseq reads (*Pa* and $Pce_S$_a, *Pf* and $Pce_S$_f) > than interspecific difference of %-covered bases from obtained RNAseq reads (*Pf* and $Pce_S$_a, *Pa* and $Pce_S$_f); (e) 367 regions were the proportion of transcripts with intraspecific amino acid identity ($Pa_T$ and $Pce_S$_a, $Pf_{eH}$ and $Pce_S$_f) < than the proportion of transcripts with interspecific amino acid identity ($Pf_{eH}$ and $Pce_S$_a , $Pa_T$ and $Pce_S$_f).

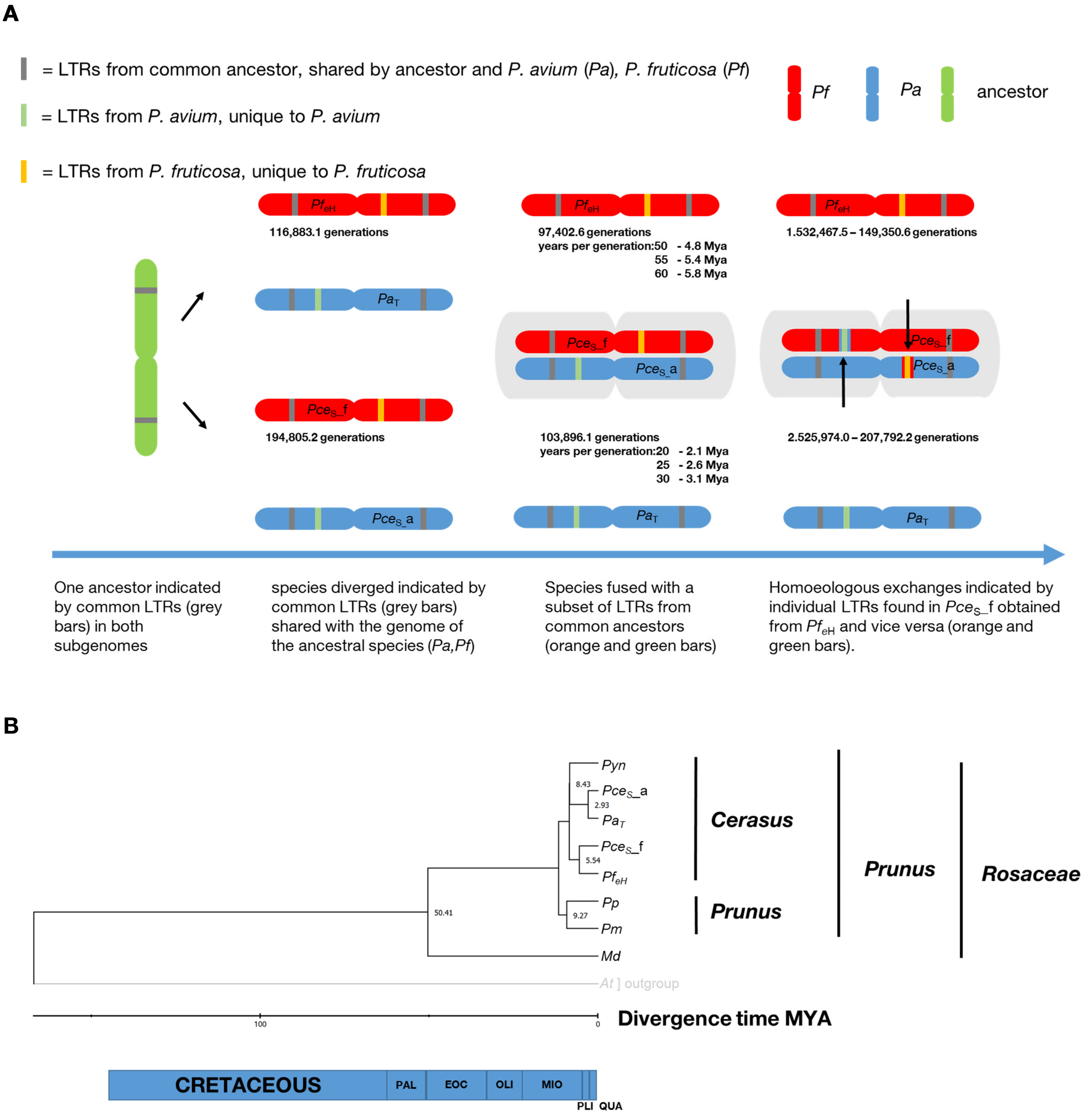


**Fig. 5** Investigation on the evolution of the genome of *P. cerasus* 'Schattenmorelle'. **A** Determination of insertion time from shared long terminal repeats (LTRs) in *P. cerasus* subgenome *avium* (*Pce*$_S$_a) and *P. cerasus* subgenome *fruticosa* (*Pce*$_S$_f) compared to *P. avium* 'Tieton' (*Pa*$_T$) and *P. fruticosa* ecotype Hármashatárhegy (*Pf*$_{eH}$). **B** Estimation of divergence of time (Mya) of *P. cerasus* subgenomes *Pce*$_S$_a and *Pce*$_S$_f compared to the donor species *P. avium* (Pa) and *P. fruticosa* (Pf). *Prunus yedonensis* (Pyn); *Prunus avium* (Pa); *Prunus persica* (Pp); *Prunus mume* (Pm); *Malus domestica* (Md); Paleocene (PAL); Eocene (EOC); Oligocene (OLI); Miocene (MIO); Pliocene (PlI); Pleistocene (PLEI).

**Table 1** Characterization of repetitive sequences of *P. fruticosa* ecotype Hármashatárhegy ($Pf_{eH}$) compared to *P. avium* 'Tieton' ($Pa_T$), *P. persica* 'Lovell', and the two subgenomes of *P. cerasus* 'Schattenmorelle' $Pce_S$_a and $Pce_S$_f

| Class | Order | Family | No. of elements | | | | | Length (bp) | | | | | Percentage of the genome (%) | | | | |
|---|---|---|---|---|---|---|---|---|---|---|---|---|---|---|---|---|---|
| | | | $Pf_{eH}$ | $Pa_T$ | *Pp* | $Pce_S$_a | $Pce_S$_f | $Pf_{eH}$ | $Pa_T$ | *Pp* | $Pce_S$_a | $Pce_S$_f | $Pf_{eH}$ | $Pa_T$ | *Pp* | $Pce_S$_a | $Pce_S$_f |
| I (retro-transposons) | **LTR** | - | 2142 | 1723 | 2607 | 2483 | 2800 | 472290 | 264621 | 446395 | 502909 | 589486 | 0.13 | 0.08 | 0.20 | 0.09 | 0.10 |
| | | Cassandra | 1852 | 1179 | 753 | 962 | 1378 | 910040 | 323669 | 329299 | 338144 | 605740 | 0.25 | 0.09 | 0.15 | 0.06 | 0.11 |
| | | Caulimovirus | 793 | 1586 | 697 | 1260 | 798 | 627333 | 861208 | 933515 | 1376203 | 692369 | 0.17 | 0.25 | 0.41 | 0.24 | 0.12 |
| | | **Copia** | **41192** | **32612** | **23578** | **25346** | **31727** | **27822528** | **12487829** | **14606294** | **15713285** | **21571049** | **7.59** | **3.64** | **6.47** | **2.76** | **3.79** |
| | | **Gypsy** | **68445** | **45868** | **25860** | **41178** | **55259** | **76652400** | **20554372** | **19004947** | **41439100** | **57972951** | **20.91** | **5.99** | **8.42** | **7.29** | **10.20** |
| | | **ERV1** | - | **94** | - | **317** | **306** | - | **17404** | - | **29672** | **32867** | - | **0.01** | - | **0.01** | **0.01** |
| | | **ERVK** | - | - | **195** | **35** | **29** | - | - | **41513** | **20720** | **17722** | - | - | **0.02** | **0.004** | **0.003** |
| | | **Pao** | **344** | - | **200** | **265** | **235** | **96802** | - | **156072** | **133246** | **117728** | **0.03** | - | **0.07** | **0.02** | **0.02** |
| | LINE | I-Jockey | 413 | 464 | 107 | - | - | 140619 | 110916 | 38834 | - | - | 0.04 | 0.03 | 0.02 | - | - |
| | | L1 | 8844 | 6349 | 4286 | 6703 | 7722 | 4167515 | 2449897 | 1392308 | 2637481 | 3118916 | 1.14 | 0.71 | 0.62 | 0.46 | 0.55 |
| | | L2 | 434 | - | 110 | - | - | 64430 | - | 21600 | - | - | 0.02 | - | 0.01 | - | - |
| | | Penelope | 176 | - | 218 | - | - | 25448 | - | 34866 | - | - | 0.01 | - | 0.02 | - | - |
| | | RTE-BovB | 516 | 214 | - | 201 | 281 | 87801 | 91608 | - | 62501 | 79770 | 0.02 | 0.03 | - | 0.01 | 0.01 |
| | | **L1-Tx1** | - | - | **211** | - | - | - | - | **46145** | - | - | - | - | **0.02** | - | - |
| | | **R1-LOA** | - | - | - | **88** | **72** | - | - | - | **27316** | **17623** | - | - | - | **0.005** | **0.003** |
| | | **RTE-X** | - | - | - | **38** | **82** | - | - | - | **52387** | **55967** | - | - | - | **0.01** | **0.01** |
| | | **R2-NeSL** | - | - | **184** | - | - | - | - | **36760** | - | - | - | - | **0.02** | - | - |
| | | **Rex-Babar** | - | - | **89** | - | - | - | - | **30378** | - | - | - | - | **0.01** | - | - |
| | | **CR1** | - | - | **712** | - | - | - | - | **609177** | - | - | - | - | **0.27** | - | - |
| | | **TAD1** | - | **37** | - | - | - | - | **6991** | - | - | - | - | **0.002** | - | - | - |
| | SINE | - | 457 | 620 | 1886 | 2039 | 1854 | 62956 | 49823 | 192330 | 213023 | 193058 | 0.02 | 0.01 | 0.09 | 0.04 | 0.03 |
| | | **ID** | - | - | **1450** | | - | - | - | **121213** | - | - | - | - | **0.05** | - | - |
| | | **tRNA-DEU-L2** | - | - | - | **587** | **578** | - | - | - | **45058** | **46403** | - | - | - | **0.01** | **0.01** |
| | | **tRNA-Core-L2** | - | - | **1109** | | - | - | - | **122977** | - | | - | - | **0.05** | - | - |
| | | B2 | 1517 | 123 | - | 670 | 619 | 122973 | 9505 | - | 49331 | 46555 | 0.03 | 0.003 | - | 0.01 | 0.01 |
| | | tRNA | 4593 | 5378 | 2229 | 3110 | 3118 | 509966 | 517003 | 203010 | 283183 | 283895 | 0.14 | 0.15 | 0.09 | 0.05 | 0.05 |

continuation of Table 1

| Class | Order | Family | No. of elements | | | | | Length (bp) | | | | | Percentage of the genome (%) | | | | |
|---|---|---|---|---|---|---|---|---|---|---|---|---|---|---|---|---|---|
| | | | $Pf_{eH}$ | $Pa_T$ | $Pp$ | $Pce_S$_a | $Pce_S$_f | $Pf_{eH}$ | $Pa_T$ | $Pp$ | $Pce_S$_a | $Pce_S$_f | $Pf_{eH}$ | $Pa_T$ | $Pp$ | $Pce_S$_a | $Pce_S$_f |
| II (DNA transposons) | | TcMar-Fot1 | 276 | 123 | - | - | - | 210022 | 48369 | - | - | - | 0.06 | 0.01 | - | - | - |
| | | **hAT-Charlie** | - | - | **1984** | - | - | - | - | **468323** | - | - | - | - | **0.21** | - | - |
| | | **IS3EU** | - | **103** | - | - | - | - | **19992** | - | - | - | - | **0.01** | - | - | - |
| | | **P** | - | **141** | - | - | - | - | **23742** | - | - | - | - | **0.01** | - | - | - |
| | | **Sola-3** | - | **111** | - | - | - | - | **42070** | - | - | - | - | **0.01** | - | - | - |
| | | **TcMAr** | - | - | **51** | - | - | - | - | **2513** | - | - | - | - | **0.00** | - | - |
| | TIR | **TcMAr-Tigger** | - | - | **152** | - | - | - | - | **121325** | - | - | - | - | **0.05** | - | - |
| Subclass I | | **Zisupton** | - | - | **141** | - | - | - | - | **18330** | - | - | - | - | **0.01** | - | - |
| | | **TcMar-ISRm11** | **81** | **55** | - | - | - | **24496** | **21760** | - | - | - | **0.01** | **0.01** | - | - | - |
| | | **TcMar-Mariner** | - | - | - | **240** | **190** | - | - | - | **57867** | **51568** | - | - | - | **0.01** | **0.01** |
| | | hAT-Ac | 11533 | 14886 | 6511 | 7832 | 10293 | 3430880 | 2101683 | 2127280 | 2654943 | 3223143 | 0.94 | 0.61 | 0.01 | 0.47 | 0.57 |
| | | hAT-Tag1 | 5353 | 5258 | 3219 | 3462 | 3909 | 1263452 | 971199 | 896657 | 799082 | 899518 | 0.34 | 0.28 | 0.40 | 0.14 | 0.16 |
| | | hAT-Tip100 | 9680 | 8622 | 6079 | 6784 | 7869 | 2348735 | 1677234 | 1336903 | 1302197 | 1611332 | 0.64 | 0.49 | 0.59 | 0.23 | 0.28 |
| | | PIF-Harbinger | 14230 | 14435 | 9390 | 10305 | 10886 | 4268364 | 3498800 | 4123535 | 3331917 | 3337330 | 1.16 | 1.02 | 1.83 | 0.59 | 0.59 |
| | | **PIF-Spy** | - | - | - | **31** | **22** | - | - | - | **1588** | **1094** | - | - | - | **0.0003** | **0.0002** |
| | Crypton | **Crypton-H/A** | **237** | - | - | **90** | **146** | **195974** | - | - | **19058** | **31113** | **0.05** | - | - | **0.003** | **0.005** |
| | Maverick | Maverick | 576 | 357 | - | 172 | 333 | 155067 | 76204 | - | 70984 | 138588 | 0.04 | 0.02 | - | 0.01 | 0.02 |
| Subclass II | | Helitron | 5378 | 4241 | 3584 | 2534 | 3367 | 2220498 | 1496918 | 1255959 | 1016700 | 1295589 | 0.61 | 0.44 | 0.56 | 0.18 | 0.23 |
| | Helitron | unknown/Helitron | 228 | 78 | 92 | 106 | 92 | 155920 | 17774 | 44752 | 84000 | 71994 | 0.04 | 0.01 | 0.02 | 0.01 | 0.01 |
| | | - | 13120 | 9946 | 9302 | 8616 | 9926 | 2310744 | 1774664 | 1865367 | 1592948 | 1972787 | 0.63 | 0.52 | 0.83 | 0.28 | 0.35 |
| | | **Academ /-2** | **42** | - | - | **479** | **590** | **20252** | - | - | **129228** | **147664** | **0.01** | - | - | **0.02** | **0.03** |
| Other | | **CMC-EnSpm** | **16958** | **14886** | **10222** | **12443** | **14104** | **8879643** | **4747818** | **12856725** | **7174757** | **7768726** | **2.42** | **1.38** | **5.70** | **1.26** | **1.37** |
| | | **Dada** | - | - | - | **110** | **113** | - | - | - | **51998** | **50989** | - | - | - | **0.01** | **0.01** |
| | | Ginger | 325 | - | 99 | - | - | 77794 | - | 12430 | - | - | 0.02 | - | 0.01 | - | - |
| | | MULE-MuDR | 17464 | 16744 | 9606 | 14069 | 14558 | 4459943 | 3751469 | 3049949 | 3668069 | 3844162 | 1.22 | 1.09 | 1.35 | 0.65 | 0.68 |
| rRNA | | | 326 | 30 | 223 | 84 | 346 | 231622 | 10508 | 334205 | 66518 | 473346 | 0.06 | 0.003 | 0.15 | 0.01 | 0.08 |
| snRNA | | | - | 55 | 127 | - | - | - | 4318 | 21457 | - | - | - | 0.001 | 0.01 | - | - |
| Satellite | | | 870 | 397 | 342 | 229 | 244 | 220737 | 88777 | 137022 | 53968 | 55261 | 0.06 | 0.03 | 0.06 | 0.01 | 0.01 |
| Simple repeat | | | 106232 | 81995 | 77870 | 82364 | 84513 | 4353840 | 3831673 | 6797949 | 3270826 | 3450293 | 1.19 | 1.12 | 3.01 | 0.58 | 0.61 |
| Low complexity | | | 19611 | 15501 | 13876 | 15436 | 16617 | 984829 | 756687 | 653107 | 769005 | 839253 | 0.27 | 0.22 | 0.29 | 0.14 | 0.15 |
| **Unknown** | | | **168094** | **171338** | **99691** | **208161** | **179132** | **42110587** | **41991523** | **21700006** | 40951426 | 37691937 | **11.49** | **12.25** | **9.61** | **7.20** | **6.63** |
| **SUM** | | | **522228** | **450112** | **319042** | **458829** | **464108** | **189663955** | **104698028** | **96191427** | **129990638** | **152397786** | **51.75** | **30.53** | **42.62** | **48.32** | **50.89** |

**Table 2** Comparison between the number of transcripts and %-IAA obtained from *P. fruticosa* ecotype Hármashatárhegy ($Pf_{eH}$) and *P. avium* cv 'Tieton' ($Pa_T$) representing the two ancestral species of *P. cerasus*

| Chromosome | No. of transcripts | No. obtained from donor species | | | %higher iAA to | | |
|---|---|---|---|---|---|---|---|
| | | $Pf_{eH}$ | $Pa_T$ | $Pf_{eH}$ & $Pa_T$ | $Pf_{eH}$ | $Pf_{eH}$ & $Pa_T$ | $Pa_T$ |
| $Pce_S$_a _Chro1 | 10675 | 6162 | 3953 | 2934 | 0.12 | 0.07 | 0.81 |
| $Pce_S$_a _Chro2 | 6511 | 3757 | 2301 | 1713 | 0.18 | 0.09 | 0.73 |
| $Pce_S$_a _Chro3 | 5597 | 3248 | 2019 | 1491 | 0.17 | 0.07 | 0.76 |
| $Pce_S$_a _Chro4 | 4969 | 2830 | 1743 | 1281 | 0.19 | 0.07 | 0.74 |
| $Pce_S$_a _Chro5 | 4446 | 2575 | 1627 | 1242 | 0.15 | 0.05 | 0.80 |
| $Pce_S$_a _Chro6 | 6963 | 4212 | 2555 | 1986 | 0.19 | 0.07 | 0.74 |
| $Pce_S$_a _Chro7 | 5280 | 3025 | 1969 | 1418 | 0.21 | 0.07 | 0.72 |
| $Pce_S$_a _Chro8 | 5257 | 3100 | 1860 | 1360 | 0.21 | 0.11 | 0.68 |
| $Pce_S$_f _Chro1 | 10133 | 6345 | 3495 | 2835 | 0.56 | 0.09 | 0.35 |
| $Pce_S$_f _Chro2 | 6254 | 3865 | 2056 | 1602 | 0.53 | 0.13 | 0.34 |
| $Pce_S$_f _Chro3 | 5869 | 3686 | 1953 | 1564 | 0.61 | 0.09 | 0.30 |
| $Pce_S$_f _Chro4 | 5362 | 3296 | 1820 | 1411 | 0.61 | 0.09 | 0.29 |
| $Pce_S$_f _Chro5 | 4032 | 2511 | 1425 | 1172 | 0.64 | 0.07 | 0.29 |
| $Pce_S$_f _Chro6 | 6968 | 4512 | 2441 | 1981 | 0.60 | 0.09 | 0.31 |
| $Pce_S$_f _Chro7 | 5033 | 3201 | 1687 | 1312 | 0.60 | 0.09 | 0.31 |
| $Pce_S$_f _Chro8 | 4925 | 3158 | 1566 | 1230 | 0.61 | 0.12 | 0.28 |
| Sum | | | | | | | |
| $Pce_S$_a * | 49698 | 28909 | 18027 | 13425 | 0.17 | 0.07 | **0.75** |
| $Pce_S$_f * | 48576 | 30574 | 16443 | 13107 | **0.59** | 0.10 | 0.32 |
| $Pce_S$* | 98274 | 59483 | 34470 | 26532 | 0.38 | 0.08 | 0.54 |

### The structure of the tetraploid sour cherry 'Schattenmorelle' (*Prunus cerasus* L.) genome reveals insights into its segmental allopolyploid nature

Thomas W. Wöhner[1], Ofere F. Emeriewen[1], Alexander H.J. Wittenberg[2], Koen Nijbroek[2], Rui Peng Wang[2], Evert-Jan Blom[2], Jens Keilwagen[4], Thomas Berner[4], Katharina J. Hoff[3], Lars Gabriel[3], Hannah Thierfeldt[3], Omar Almolla[5], Lorenzo Barchi[5], Mirko Schuster[1], Janne Lempe[1], Andreas Peil[1], Henryk Flachowsky[1]

[1]Julius Kühn Institute (JKI) – Federal Research Centre for Cultivated Plants, Institute for Breeding Research on Fruit Crops, Pillnitzer Platz 3a, D-01326, Dresden, Germany

[2]Keygene N.V., P.O. Box 216, 6700 AE Wageningen, Netherlands

[3]Institute of Mathematics and Computer Science, University of Greifswald, Walther-Rathenau-Str. 47, 17489 Greifswald, Germany

[4]Julius Kühn Institute (JKI) – Federal Research Centre for Cultivated Plants, Institute for Biosafety in Plant Biotechnology, Erwin-Baur-Str. 27, D-06484 Quedlinburg, Germany

[5]DISAFA – Plant genetics, University of Turin, Grugliasco (TO), 10095 Italy

**Keywords:**

genome assembly, *P. cerasus*, sour cherry, tetraploid

**This file contains information about supplemental material and methods**

#### 1. Supplemental material and methods

*1.1 Plant Material and RNA extraction, sequencing and iso-seq analysis*

*Prunus cerasus* L. 'Schattenmorelle' (accession KIZC99-2) young leaf material (tetraploid, small tree, size ca. 1,5 m, BioProject accession PRJNA742509) was collected from single grafted trees grown in the experimental field of the Julius Kühn Institute (JKI) – Federal Research Centre for Cultivated Plants, Institute for Breeding Research on Fruit Crops, Dresden, Germany (Figure 1, coordinates 50.996389 N 13.885465 E).

Two pools were generated for RNA extraction (pool 1: mature leaves, premature leaves, flower buds, vegetative buds and pool 2: open flower after pollination, green fruits, fruits in color change from green to red).

*1.2 De novo assembly and scaffolding*

To provide compatibility with Phase Genomics Hi-C scaffolding pipeline, separation of the ancestral genomes was performed by mapping publicly available *Prunus avium* reads (DRA004760, DRA004761, DRA004762, DRA004763, DRA004764, DRA004765, DRA004768, DRA004769, DRA004770, DRA004771, DRA004772) to the preliminary assembly with BWA-Mem (Li and Durbin 2009, Li and Durbin 2010). A combination of external python packages (scipy linregress, scipy find_peaks) were used to select contigs that fit the hypothesis of 1 or more clear coverage peaks on all *Prunus avium* derived contigs. The remaining contigs were then assigned to *Prunus fruticosa*. To test whether the separation was successful, the two subgenomes $\mathit{Pce}_{S}$_a and $\mathit{Pce}_{S}$_f were purged using the purge_dups (V1.0.1).

*1.3 Correctness, completeness and contiguity of the Prunus cerasus genome sequence*

K-mer analysis was performed due to the following steps: adapter sequences of both short-read sequence data sets (125 bp and 150bp paired end) were trimmed with Trimmomatic software (Bolger et al. 2014) and the trimmed data were used to estimate the best k-mer size using the genomic k-mer counting software Meryl (Galaxy Version 1.3+galaxy2, Rhie 2020): count the occurrence of canonical k-mers | estimate the best k-mer size | 300 Mb | 0.001. A database file was processed for each data set. Using the union-sum function, the resulting database files were merged.

*1.4 Structural and functional annotation*

*1.4.1 Data preparation*

Repeat masking for structural genome annotation was performed with RepeatModeler2 using the following dependency software versions: rmblast 2.11.0+, TRF 4.09 (Benson, 1999), RECON (Bao and Eddy, 2002), RepeatScout 1.0.6, RepeatMasker 4.1.2, LTR Structural Analysis: Enabled (GenomeTools 1.6.2 (Gremme et al., 2013), LTR_Retriever (Ou and Jiang, 2018) v2.9.0, Ninja (Wheeler, 2009) 0.95-cluster_only, MAFFT (Katoh and Standley, 2013) 7.487, CD-HIT (Fu et al, 2012) 4.8.1).

*1.4.2 Long read integration*

A primary gene set (transcriptome.gff) was generated with Cupcake.

Coverage information of PacBio transcripts is by default stored in the header of the transcript FASTA file. Genome annotation pipelines have not yet been adapted to use the information this way, but they require coverage information for adequate data incorporation. Therefore, we created coverage-equivalent redundant copies of transcripts with a coverage > 1 using a custom script explode_pacbio_ccs.pl (availabe at https://github.com/Gaius-Augustus/BRAKER/blob/long_reads/scripts/explode_pacbio_ccs.pl) as follows:

```
cat longreads.fastq | ./explode_pacbio_ccs.pl > exploded.fq
```

This file was spliced-aligned to the genome using Minimap2 version 2.17-r941 (Li, 2018). The resuling SAM file was converted to BAM format using SAMtools (Li et al., 2009). The resulting BAM file was provided to BRAKER1 (Hoff et al., 2016; Hoff et al., 2019) as input.

In addition, the Cupcake transcripts were processed as follows: The script stringtie2fa.py (available at https://github.com/Gaius-Augustus/Augustus/blob/master/scripts/stringtie2fa.py) was used to convert the Cupcake GTF-file into transcripts with the following command line:

```
stringtie2fa.py -g genome.chr.fa.masked -f transcriptome.chr.gff ¥     -o
cupcake.fa
```

GeneMarkS-T version 5.1 March 2014 was executed to find CDS in transcript fasta file as follows:

```
gmst.pl --strand direct cupcake.fa.mrna --output gmst.out ¥
      --format GFF
```

The local CDS coordinates were projected to the genome with another custom script (available at https://github.com/Gaius-Augustus/BRAKER/blob/long_reads/scripts/gmst2globalCoords.py ):

```
gmst2globalCoords.py -t transcriptome.chr.gff -p gmst.out ¥
      -o gmst.global.gtf -g genome.chr.fa.masked
```

The global coordinate GTF-File was converted to Hints for Augustus with another custom script contained in folder scripts (availble at https://github.com/Gaius-Augustus/BRAKER/blob/long_reads/scripts/gmst_global2hints.pl), the source key is M, i.e. the prediction of these hints will be enforced in AUGUSTUS:

gmst_global2hints.pl gmst.global.gtf > pacbio.hints

*1.4.3. Combination of BRAKER gene sets with TSEBRA*

BRAKER1, BRAKER2, and long read gene sets were combined with a modified version of the TSEBRA combiner tool (modified version available at https://github.com/Gaius-Augustus/TSEBRA/tree/long_reads) using a custom configuration file:

```
tsebra.py -g braker1/augustus.hints.gtf,braker2/augustus.hints.gtf ¥   -e
braker1/hintsfile.gff,braker2/hintsfile.gff ¥
      -l gmst.global.gtf -c long_reads.cfg -o tsebra.gtf
```

Content of the custom TSEBRA configuration file (long_reads.cfg):

```
# Weight for each hint source
# Values have to be >= 0
P 31
E 0.150
C 15
M 100.5
L 0.5
# Required fraction of supported introns or supported start/stop-codons for a transcript
# Values have to be in [0,1]
intron_support 10.8
stopstasto_support 1
start_support 2
# Allowed difference for each feature
# Values have to be in [0,1]
e_1 0.1
e_2 0.51
e_3 1
# Values have to be >0
e_34 25300
e_54 1050
e_6 20
```

*1.4.4. Gene structure prediction using GeMoMa*

Homology-based gene annotation was performed with GeMoMa version 1.9 (Keilwagen et al. 2019) using the mapped RNA-seq data from *P. cerasus* cv 'Schattenmorelle' and the genome and gene annotation from the following reference organisms that are available at NCBI: *Arabidopsis thaliana* (TAIR10.1, RefSeq GCF_000001735.4), *Vitis vinifera* (12x, RefSeq GCF_000003745.3), *Populus trichocarpa* (Pop_tri_v3, GCF_000002775.4).

Other references were downloaded from the GDR database (www. https://www.rosaceae.org): *M. domestica* (*Malus* x *domestica* HFTH1 Whole Genome v1.0), *F. vesca* (*Fragaria vesca* Whole Genome v4.0.a1), *P. avium* (*Prunus avium* Tieton Genome v2.0), *P. persica* (*Prunus persica* Whole Genome Assembly v2.0, v2.0.a1), *P. dulcis* (*Prunus dulcis* Lauranne Genome v1.0) and *P. armeniaca* (*Prunus armeniaca* Marouch n14 Whole Genome v1.0), *P. yedonensis* (*Prunus yedoensis* var. *nudiflora* Genome v1.0), *P. domestica* (*Prunus domestica* Draft Genome Assembly v1.0), *P. communis* (*Pyrus communis* Bartlett DH Genome v2.0), *R. occidentalis* (*Rubus occidentalis* Whole Genome v3.0). *P. fruticosa* was downloaded from OpenAgrar (Wöhner et al. 2021 b).

*1.4.5 Final gene structure generation*

BUSCO version 5.2.2 with set embryophyta_odb10 (number of genomes: 50, number of BUSCOs: 1614) was used for the assessment of protein completeness. For handling alternative transcripts correctly and not as duplicates, a custom script was ran on the BUSCO full table, assigning gene ID instead of transcript ID.

The chloroplast and mitochondria sequences were annotated with GeSeq (Tillich et al. 2017) using the all available references for chloroplast (*P. armenica, P. avium, P. campanulata, P. cerasoides, P. davidiana, P. dictyoneura, P. domestica, P. dulcis, P. humilis, P. kansuensis, P. matuurae, P. maximowiczii, P. mira, P. mongolica, P. mume, P. penduculata, P. persica, P. pseudocerasus, P. rufa, P. salicina, P. serotine, P. speciosa, P. takesimensis, P. tenella, P. tomentosa, P. triloba, P. yedonensis, P. zippeliana*) from NCBI and mitochondria from *P. avium* (GenBank accession MK816392) published by Yan et al. (2019). GeSeq pipeline analysis was performed using the annotation packages ARAGORN, blatN, blatX, Chloe and HMMER.

*1.4.6 Protein clustering, multiple sequence alignment and divergence of time estimation*

The Proteinortho (Galaxy Version 6.0.32+galaxy0) was used to find orthologous proteins within the datasets with the following parameters: LAST, e-value treshold = 0.001, minimal algebraic connectivity = 0.1, in add. options: Minimal coverage of best alignment in % = 50, min. seq. similarity in % = 95, minimal percent identity of best blast hits in % = 25.

MAFFT (Galaxy Version 7.505+galaxy0, Katoh and Standley 2013) was used to align each obtained orthogroup with the following parameters: gap extent 0.123, gap open 1.53, no matrix, output fasta.

A timetree was inferred by applying the RelTime method (Tamura et al. 2012, Tamura et al. 2018) to the user-supplied phylogenetic tree whose branch lengths were calculated using the Ordinary Least Squares method. The timetree was computed using 1 calibration constraint. The Tao et al. (2020) method was used to set minimum and maximum time boundaries on nodes for which calibration densities were provided. Confidence intervals were computed using the Tao et al. (2020) method. The evolutionary distances were computed using the JTT matrix-based method (Jones et al. 1992) and are in the units of the number of amino acid substitutions per site. The rate variation among sites was modelled with a gamma distribution (shape parameter = 1). This analysis involved nine amino acid sequences. There were 419,586 positions in the final dataset.

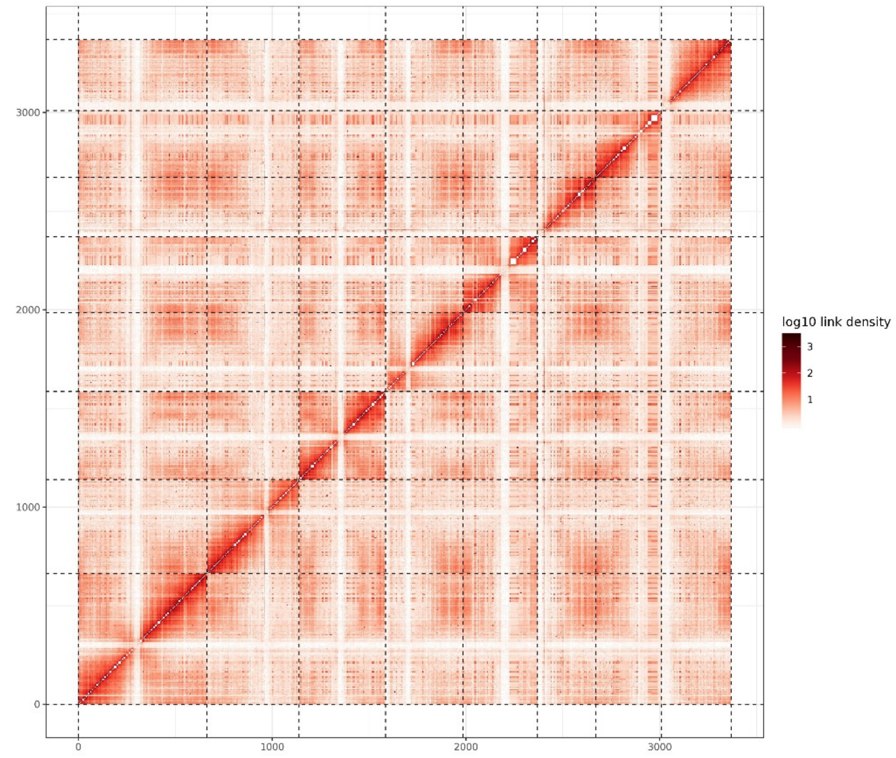


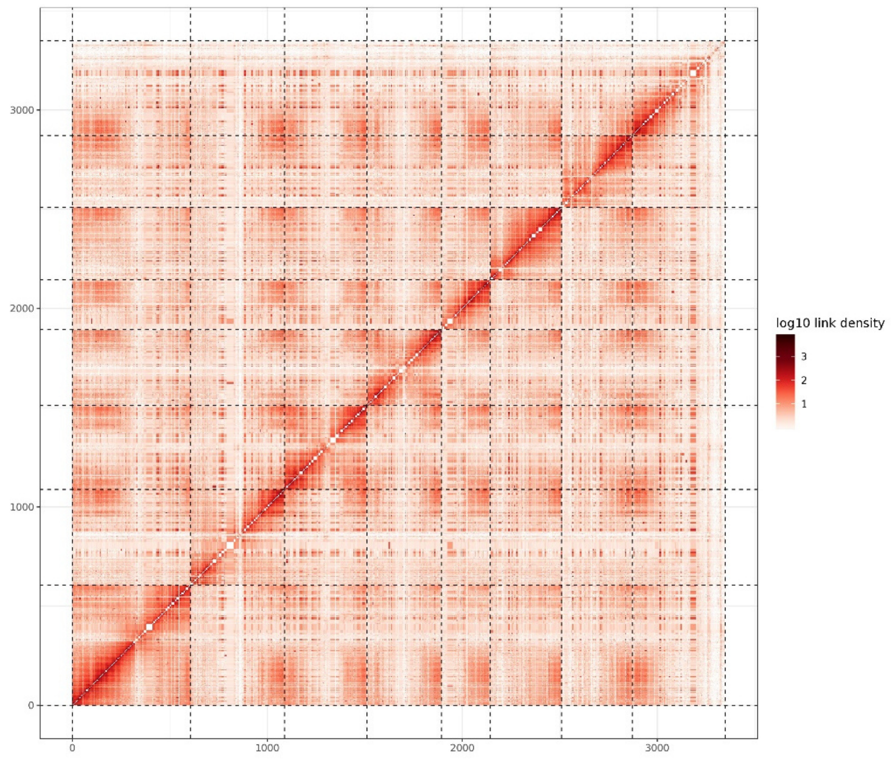


**Figure S1** Hi-C heatmap post-scaffolding for the subgenomes *Pce*$_S$_a (left) and *Pce*$_S$_f (right) of *P. cerasus* cv 'Schattenmorelle'. The heatmap indicates the density of paired Hi-C reads which interact to each other in close proximity. High intense colour indicates high interaction.

(A)

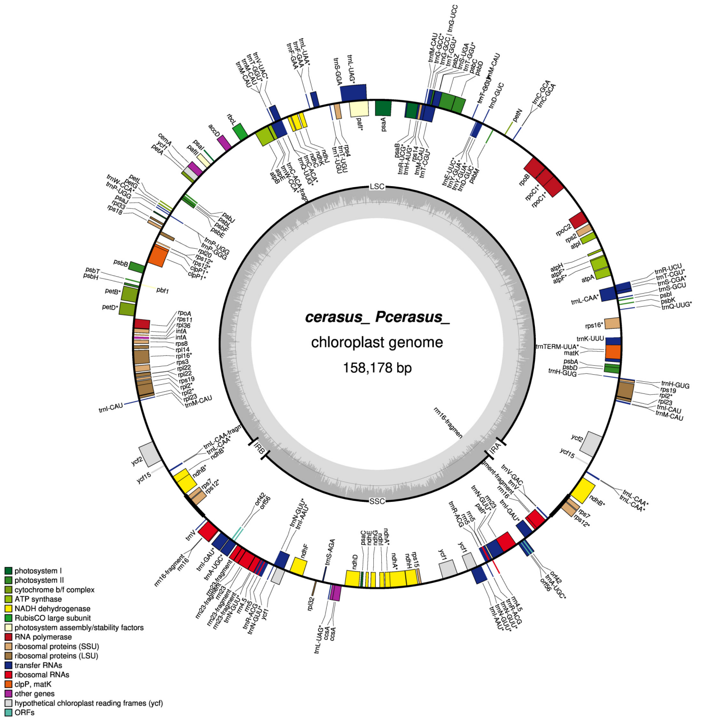


(B)

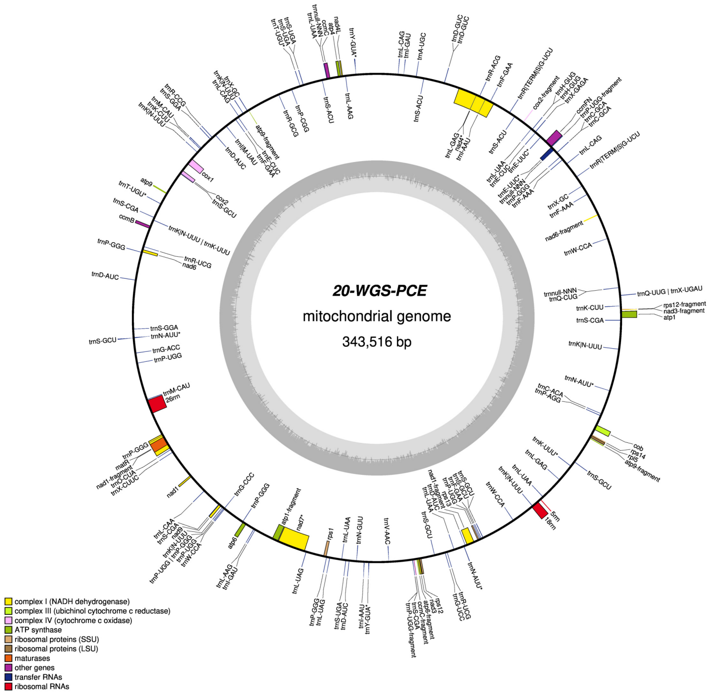


**Figure S2** The chloroplast (a) and mitochondrial (b) sequences of *P. cerasus* L. cv 'Schattenmorelle'.

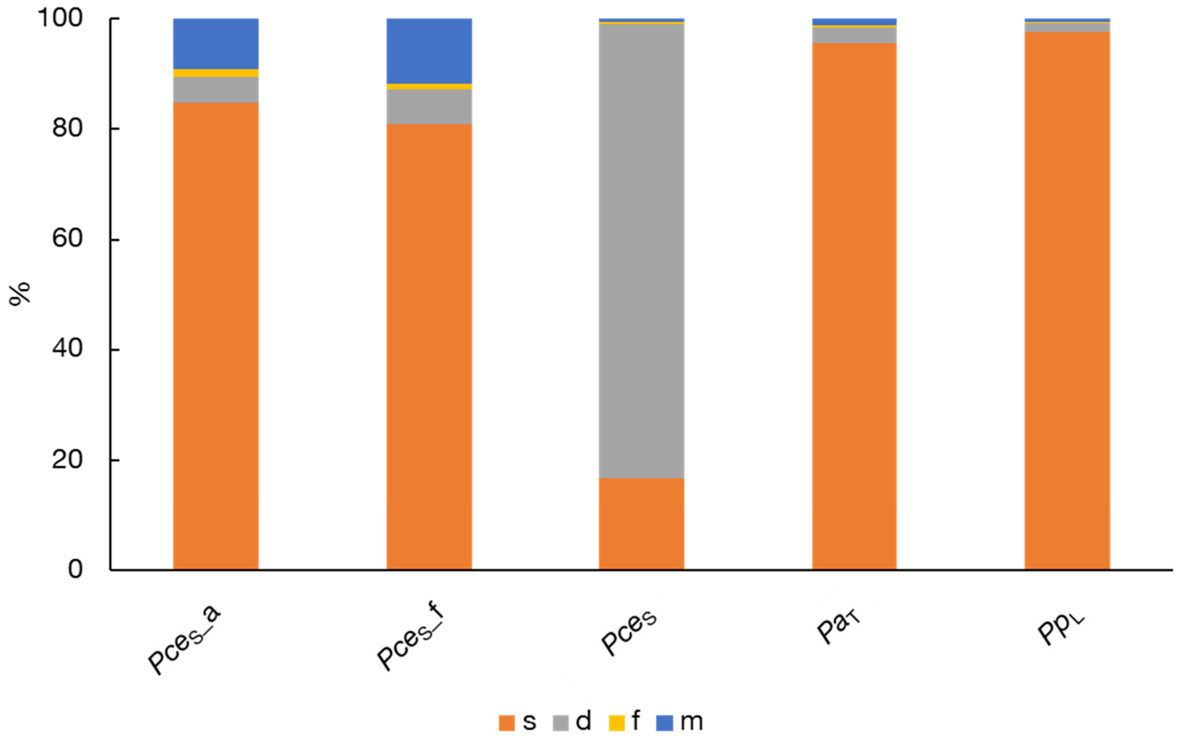


**Figure S3** Analysis of completeness of the *P. cerasus* cv. Schattenmorelle subgenomes *P. cerasus* cv 'Schattenmorelle' subgenome *avium* (*Pce*$_S$_a) and *P. cerasus* cv 'Schattenmorelle' subgenome *fruticosa* (*Pce*$_S$_f) and combined datasets compared to *P. avium* cv. 'Tieton' (*Pa*$_T$) and *P. persica* cv. Lovell (*Pp*$_L$) by mapping of a set of universal single-copy orthologs using BUSCO*.* The bar charts indicate complete **s**ingle copy (orange), complete **d**uplicated (gray), **f**ragmented (yellow) and **m**issing (blue) genes. For evaluation the embryophyta_odb10 BUSCO dataset (n=1614) was used. *P. cerasus* cv. Schattenmorelle show a 99 % completeness (S: 16.7 %, D: 82.3 %, F: 0.4 %, M: 0.6 %, n: 1614) which reaches the completeness of *P. avium* cv. 'Tieton' (C: 98.3 %, S: 95.6 %, D: 2.7 %, F: 0.5 %, M:1.5 %, n:1614) and *P. persica* 'Lovell' (C: 99.3 %, S: 97.5 %, D: 1.8%, F: 0.1 %, M: 0.6 %, n:1614).

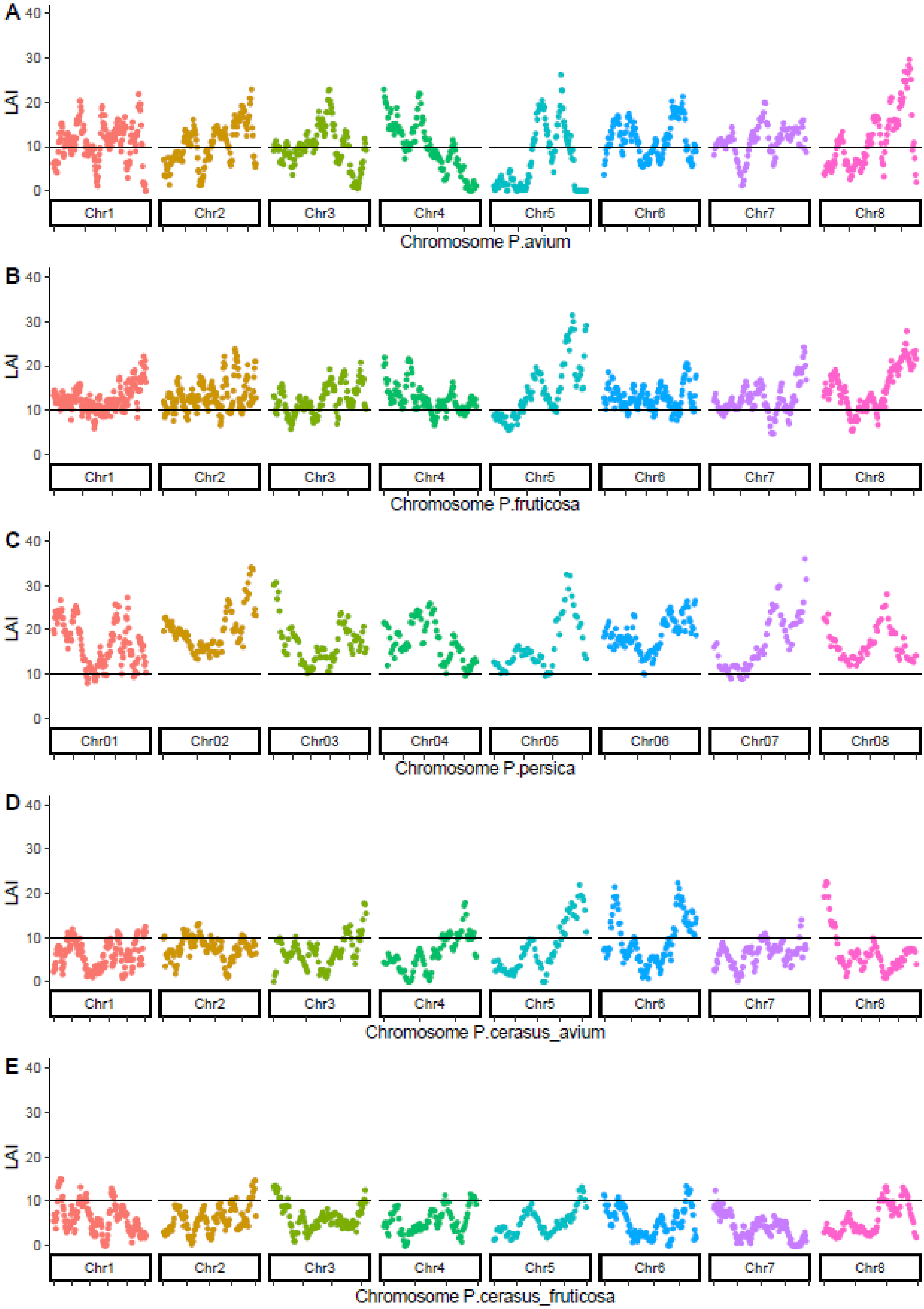
A
B
C
D
E
LAI
40
30
20
10
0
Chr1
Chr2
Chr3
Chr4
Chr5
Chr6
Chr7
Chr8
Chr01
Chr02
Chr03
Chr04
Chr05
Chr06
Chr07
Chr08
Chromosome P.avium
Chromosome P.fruticosa
Chromosome P.persica
Chromosome P.cerasus_avium
Chromosome P.cerasus_fruticosa

**Figure S4** Assessing the quality of repetitive sequences between the chromosome sequences of *P. avium* ‘Tieton’ (A), *P. fruticosa* ecotype Hármashatárhegy (B), *P. perisca* 'Lovell', and (C) *P. cerasus* subgenome *avium* and *P. cerasus* subgenome *fruticosa* using the LAI index. The genomes *P. cerasus* [this study] and *P. avium* were sequenced with ONT 9.4.1 and Illumina (Wang wet al. 2020), *P. fruticosa* with ONT 10.3 (Wöhner et al. 2021) and *P. persica* with Illumina and Sanger sequencing of fosmid and BAC clones (Verde et al. 2017).

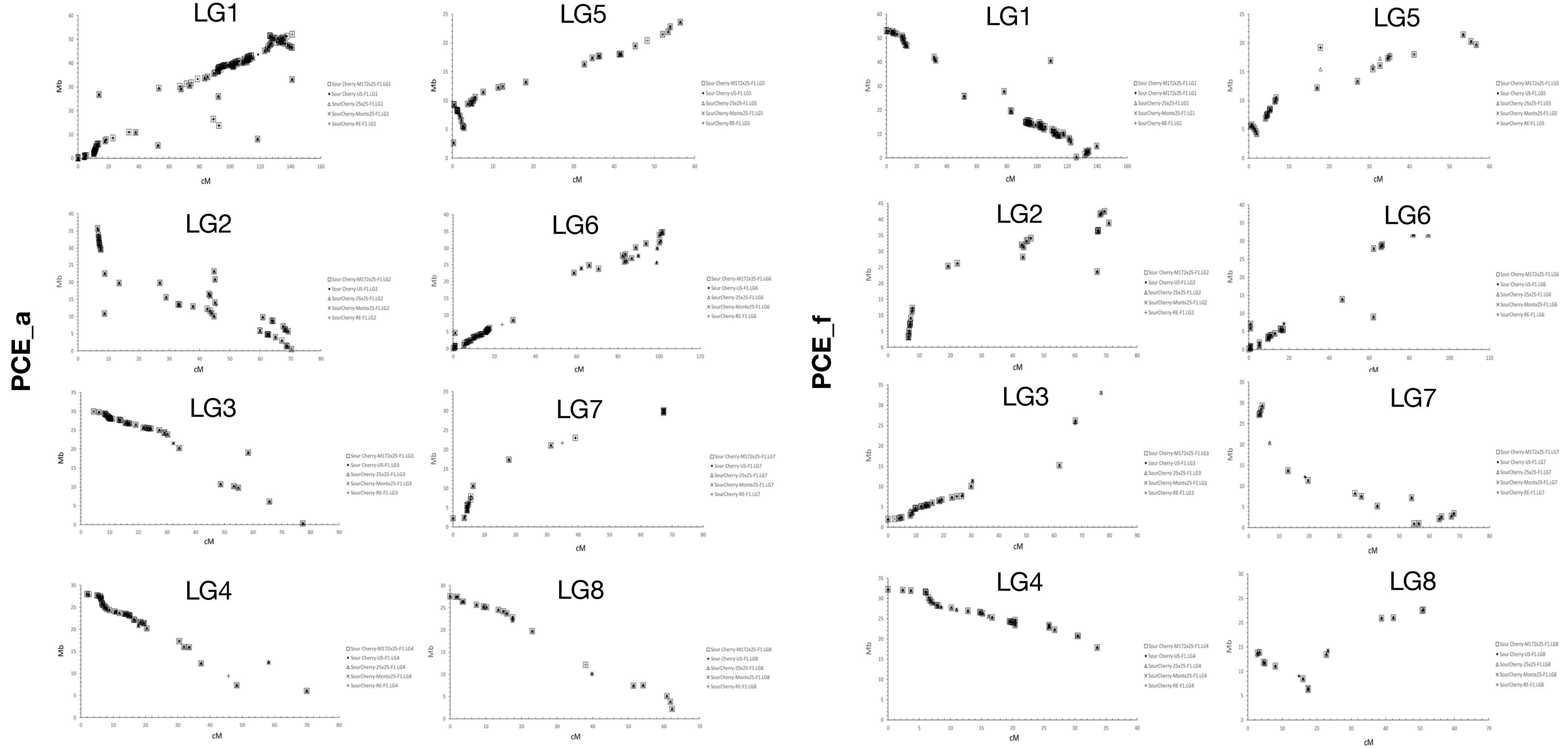


**Figure S5** Collinearity plots between the five published genetic maps of sour cherry (M172x25-F1, US-F1, 25x25-F1, Montx25-F1, RE-F1) and the *P. cerasus* cv 'Schattenmorelle' subgenome *avium* (*Pce*$_S$_a) and *fruticosa* (*Pce*$_S$_f).X-axis represents the genetic position of a marker in the genetic linkage map given in centi Morgan (cM). Y-axis represents the physical position of a marker sequence within the genome sequence of the respective subgenome given in Mega base pairs (Mbp).

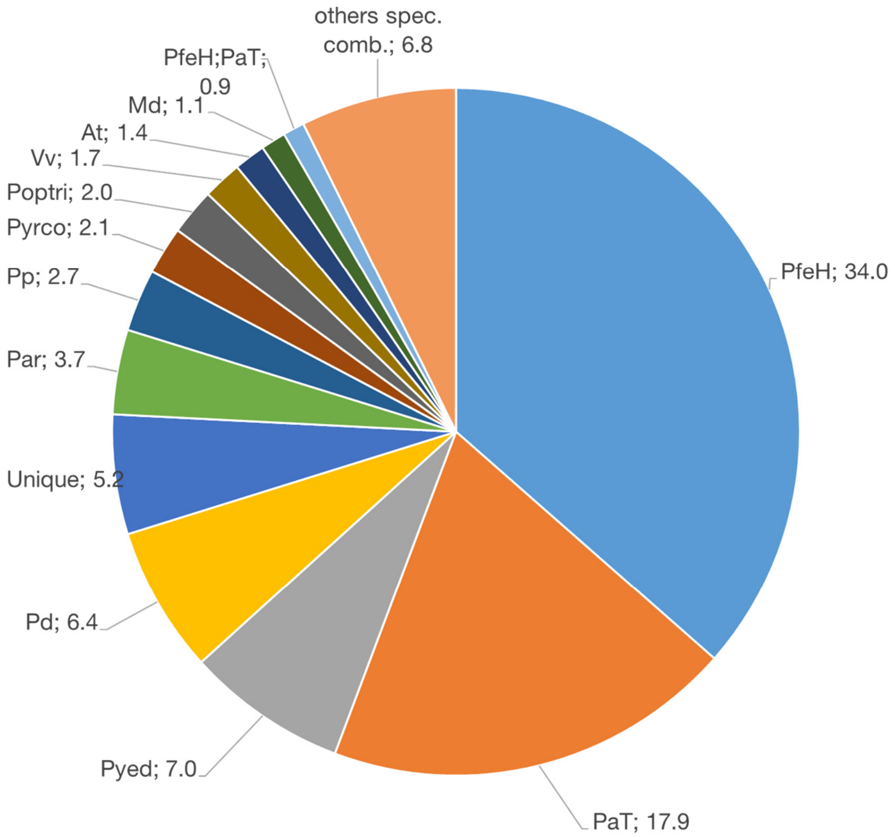


**Figure S6** Percentage of *P. cerasus* (*Pce*) proteins by IAA compared with 15 reference species. *P. fruticosa* ecotype Hármashatárhegy ($P_{feH}$), *P. avium* 'Tieton' ($Pa_{T}$), *P. yedonensis* (*Pyed*), *P. domestica* (*Pd*), *P. armeniaca* (*Par*), *P. persica* (*Pp*), *Pyrus communis* (*Pyrco*), *Populus trichocarpa* (*Poptri*), *Vitis vinifera* (*Vv*), *Arabidopsis thaliana* (*At*), *Malus domestica* (*Md*)

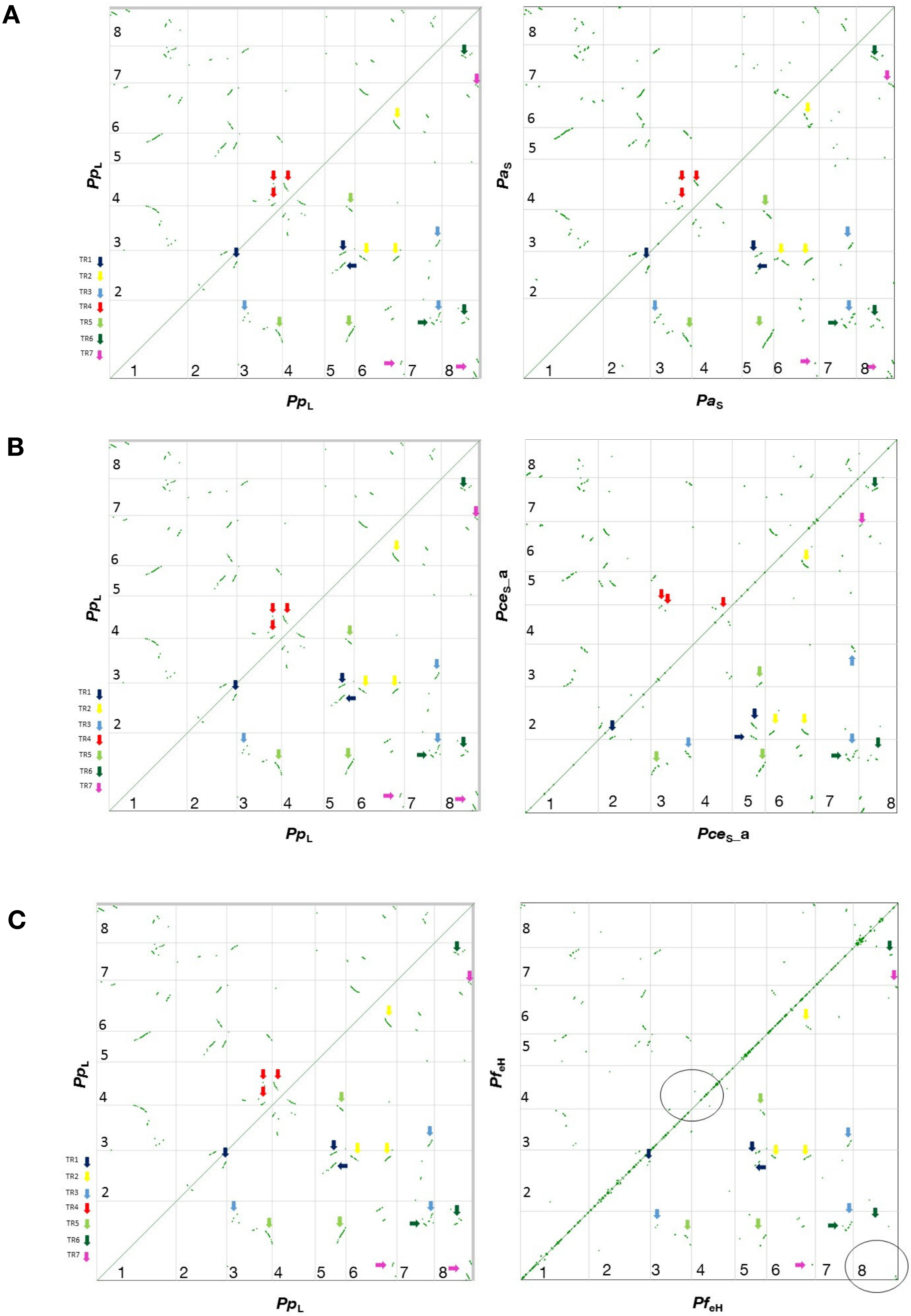
A
B
C
TR1
TR2
TR3
TR4
TR5
TR6
TR7
PpL
PaS
PceS_a
PfeH
1
2
3
4
5
6
7
8

**D**

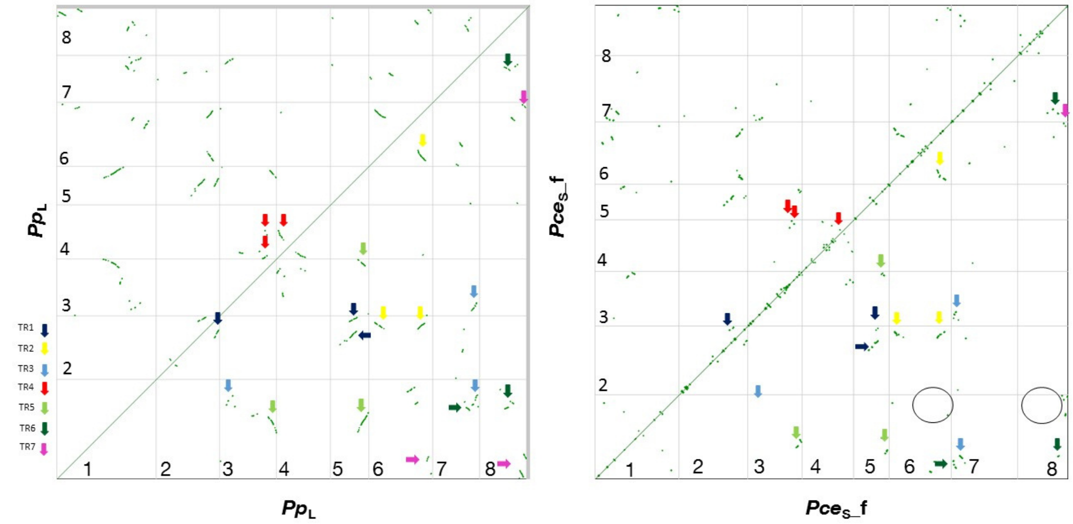


**Figure S7** Synmap2 plots of self-self comparisons between (A) *Prunus persica* 'Lovell' (*Pp*$_L$) and *P. avium* 'Sato Nishiki' (*Pa*$_S$), (B) *P. fruticosa* 'Hármashatárhegy' (*Pf*$_{eH}$), (C) *P. cerasus_avium* 'Schattenmorelle' (*Pce*$_S$_a), (D) *P. cerasus_fruticosa* 'Schattenmorelle' (*Pce*$_S$_f) for the identification of triplicated regions (TR) 1-7.

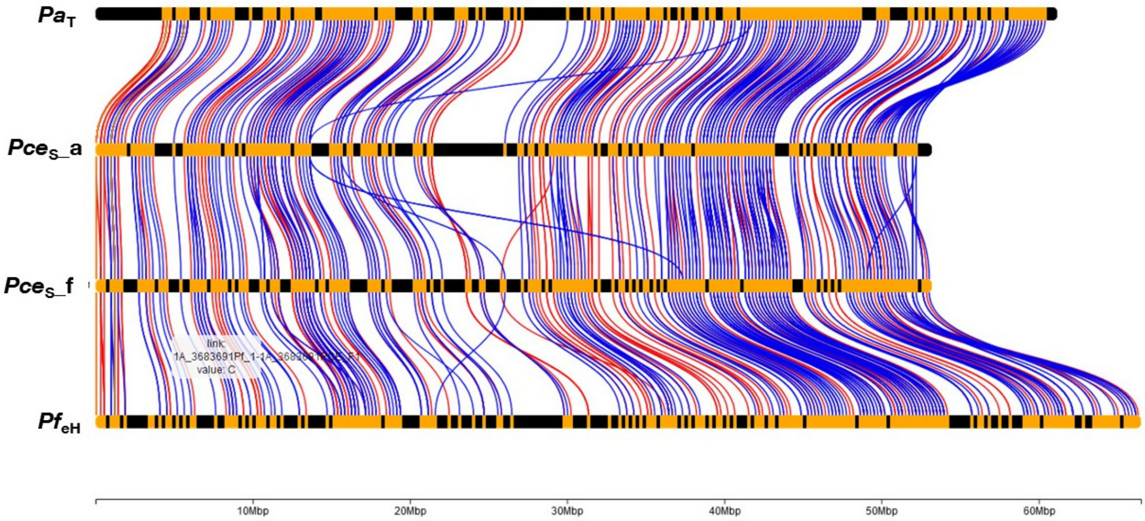


**Chr1**

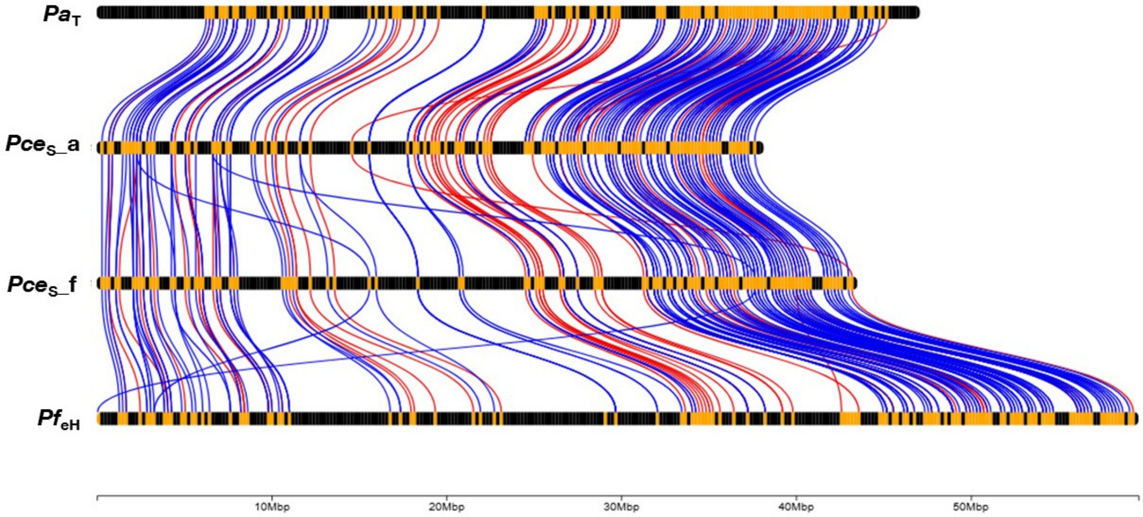


**Chr2**

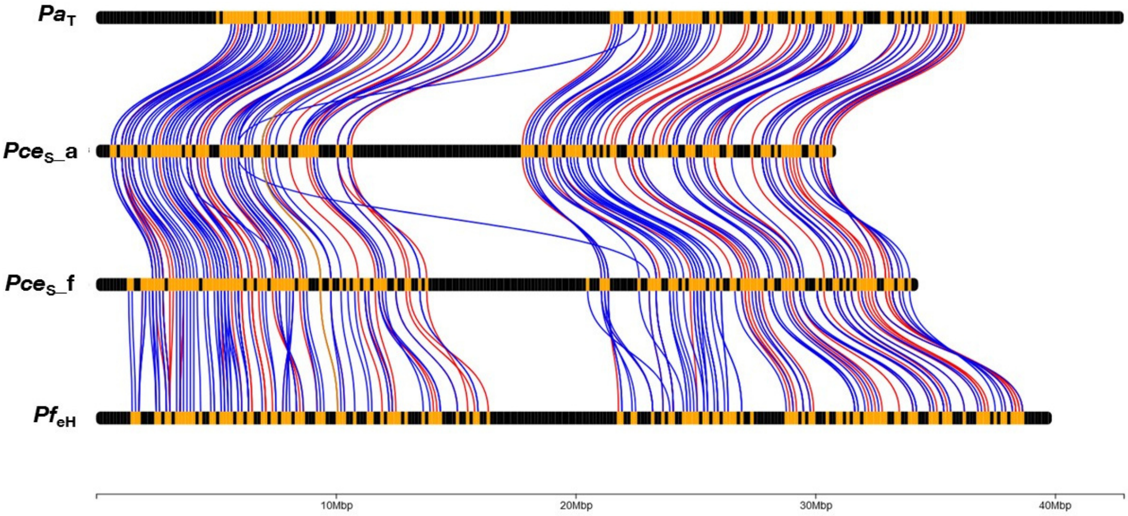


**Chr3**

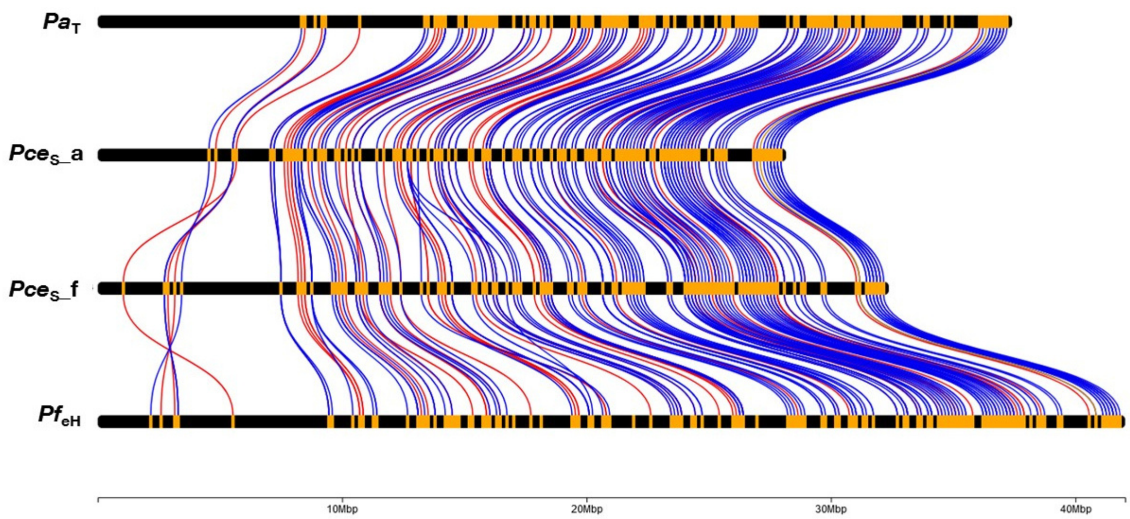


**Chr4**

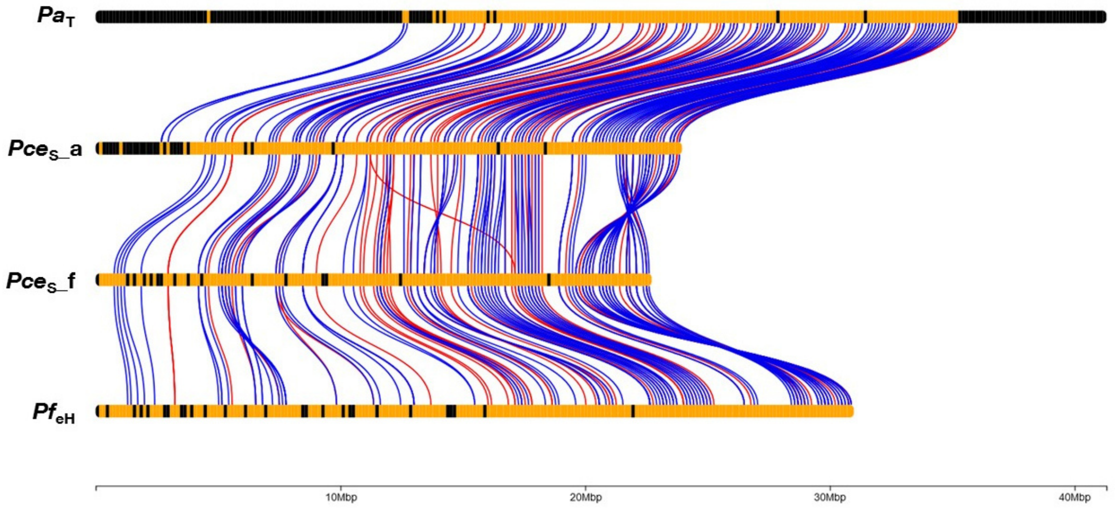


**Chr5**

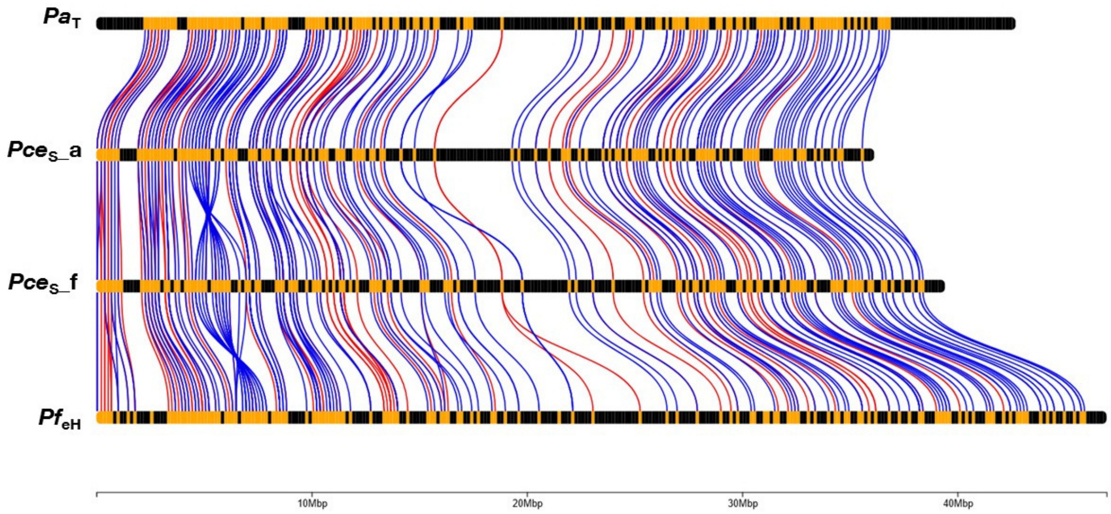


**Chr6**

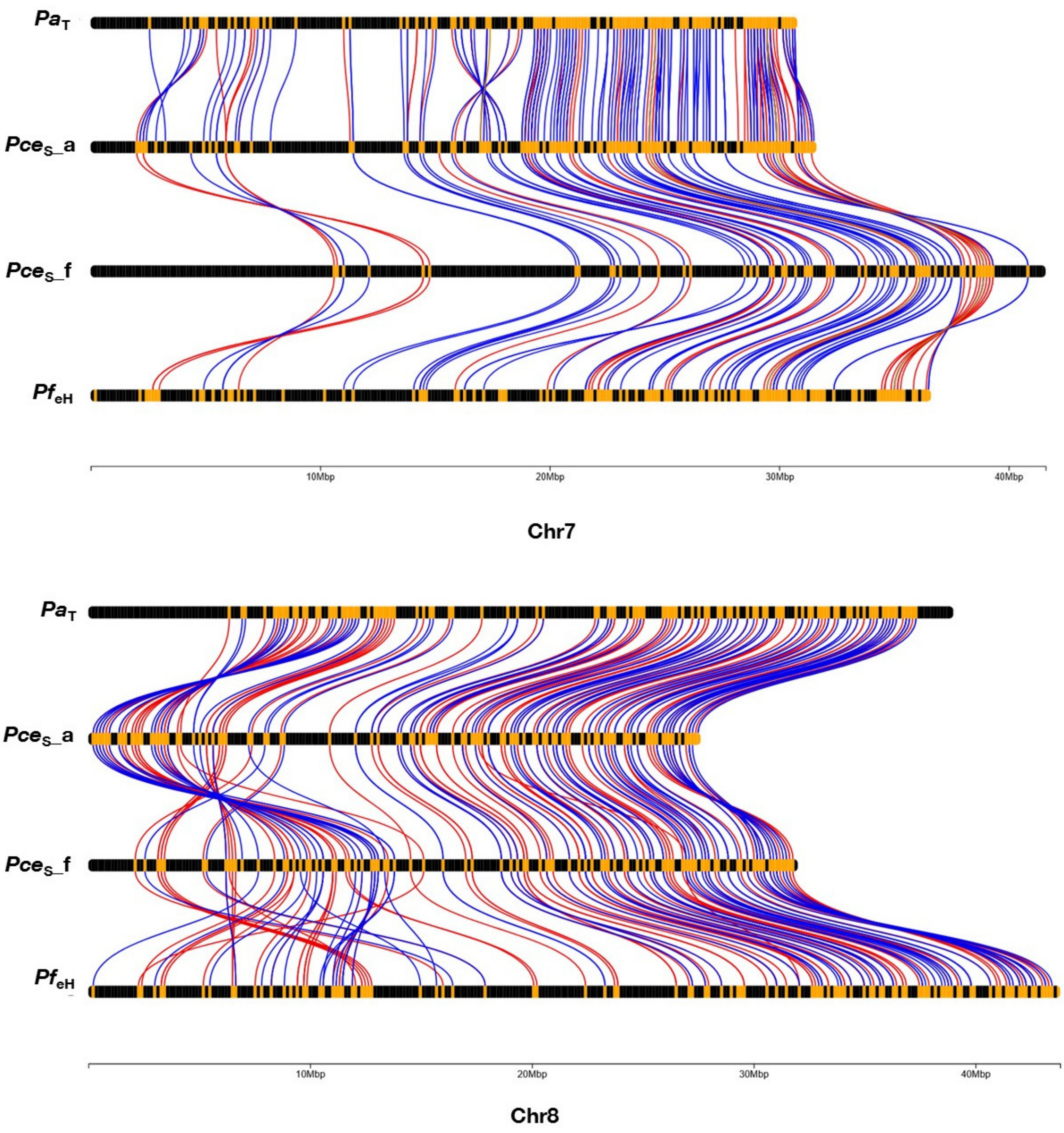


**Figure S8** Positional co-linearity comparison between the two subgenomes *P. cerasus_avium* 'Schattenmorelle' (*Pce*$_{S}$_a), *P. cerasus_fruticosa* ‘Schattenmorelle’ (*Pce*$_{S}$_f) and *P. avium* 'Tieton' (*Pa*$_{T}$), *P. fruticosa* 'Hármashatárhegy' (*Pf*$_{eH}$), using the molecular markers from the 9+6k SNP array. The plots were generated using the R-software package chromoMap v0.4.1.

A

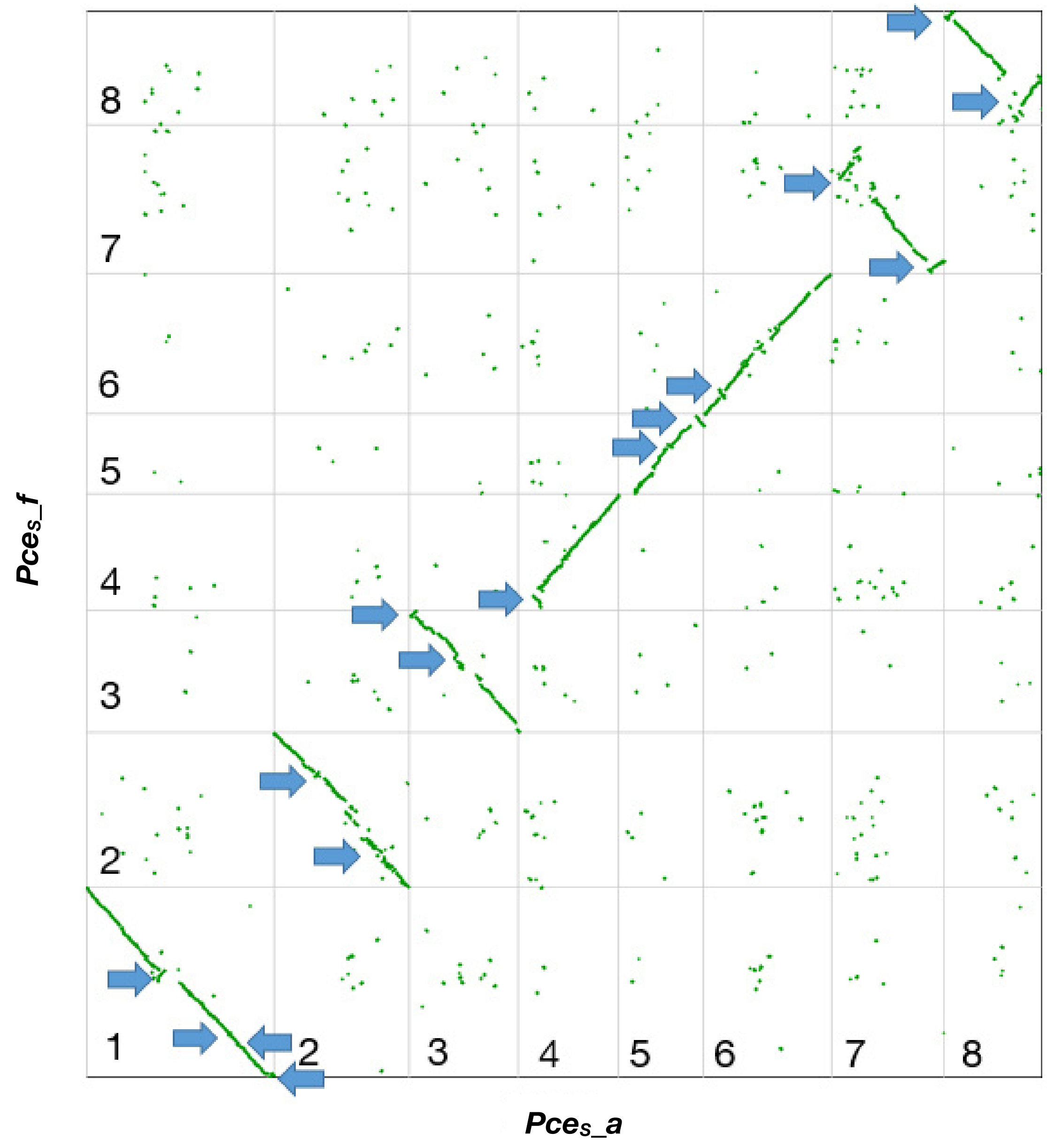

**B**

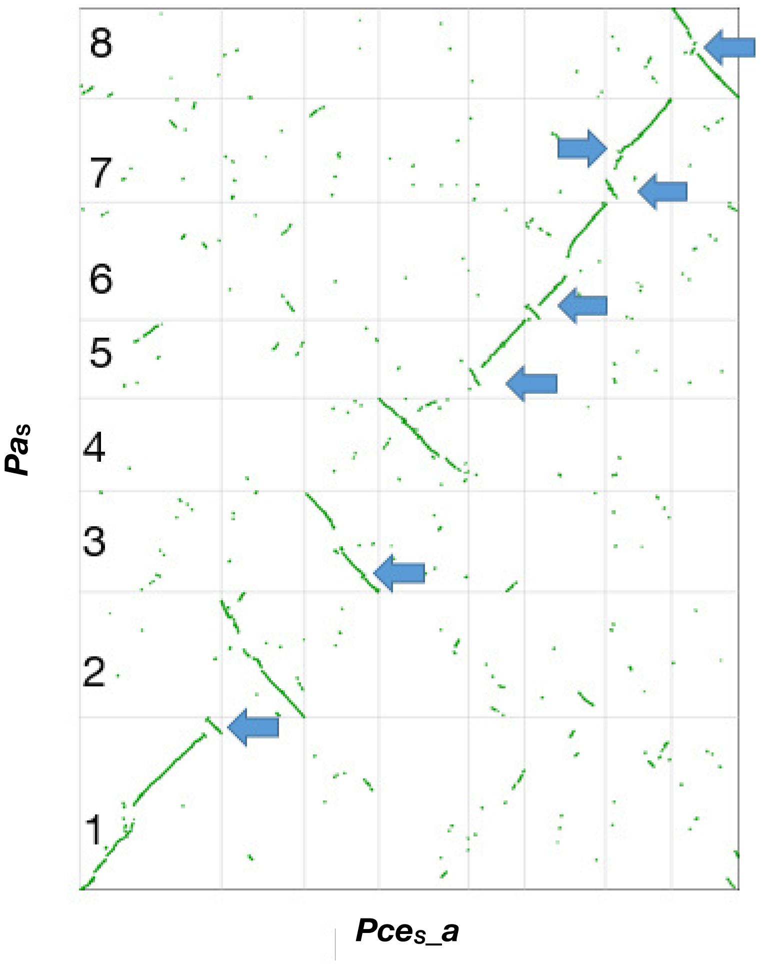


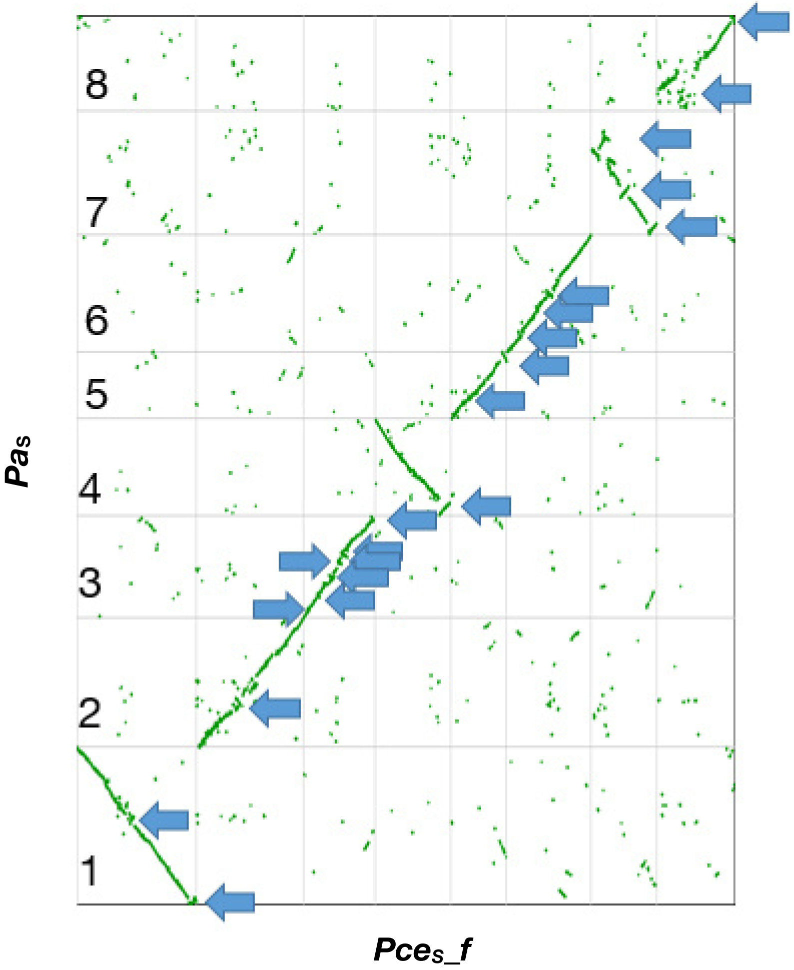


**Figure S9** Synteny between (a) *P. cerasus* subgenome *_avium* (PceS_a) and *P. cerasus* subgenome *_fruticosa* (PceS_f), and (b) the subgenomes and the genotypes *Prunus avium* 'Tieton' ($Pa_T$) and *P. fruticosa* ($Pf_{eH}$) of the ancestral species *P. avium* and *P. fruticosa.* Blue arrows indicate positions where inversions occurred.

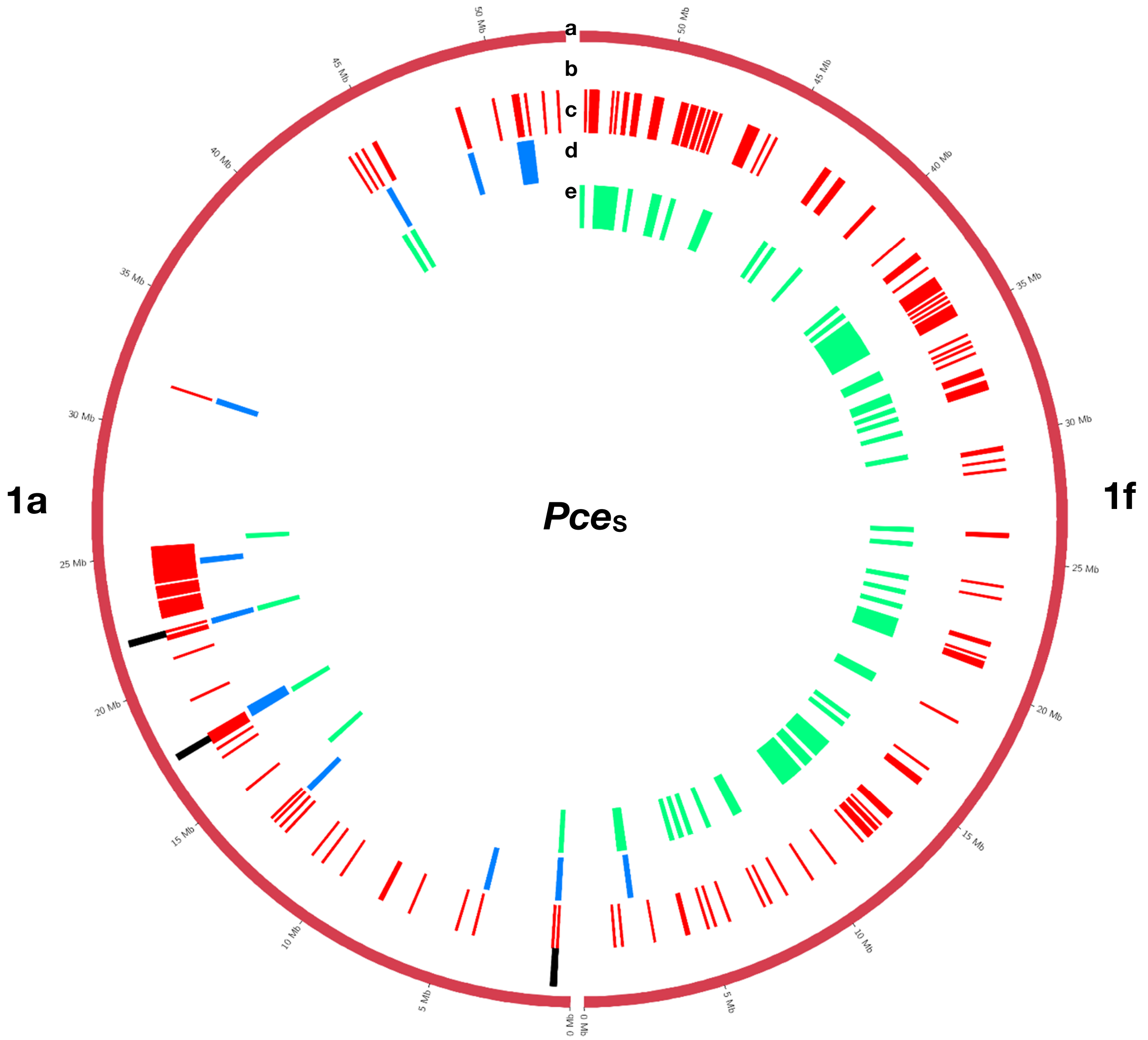


**Figure S10** Detected regions of homoeologous exchanges in the genome of *P. cerasus* 'Schattenmorelle'. Circos plot of 16 single pseudomolecules 1 to 8 (see continuating plots) of the subgenomes of *Pce*$_{S}$_a and *Pce*$_{S}$_f. (a) chromosome length (Mb); (b) region in *Pce*$_{S}$_a and in *Pce*$_{S}$_f detected that match all three following analysis methods: (c) regions (100k window) were intraspecific %-covered bases from mapped reads (*Pce*$_{S}$_a to *Pa*$_{T}$, *Pce*$_{S}$_f to *Pf*$_{eH}$) was < than interspecific %-covered bases from mapped reads (*Pce*$_{S}$_a to *Pf*$_{eH}$, *Pce*$_{S}$_f to *Pa*$_{T}$); (d) regions were intraspecific difference of %-covered bases from obtained RNAseq reads (*Pa* and *Pce*$_{S}$_a, *Pf* and *Pce*$_{S}$_f) > than interspecific difference of %-covered bases from obtained RNAseq reads (*Pf* and *Pce*$_{S}$_a, *Pa* and *Pce*$_{S}$_f); (e) regions were the proportion of transcripts with intraspecific amino acid identity (*Pa*$_{T}$ and *Pce*$_{S}$_a, *Pf*$_{eH}$ and *Pce*$_{S}$_f) < than the proportion of transcripts with interspecific amino acid identity (*Pf*$_{eH}$ and *Pce*$_{S}$_a , *Pa*$_{T}$ and *Pce*$_{S}$_f).

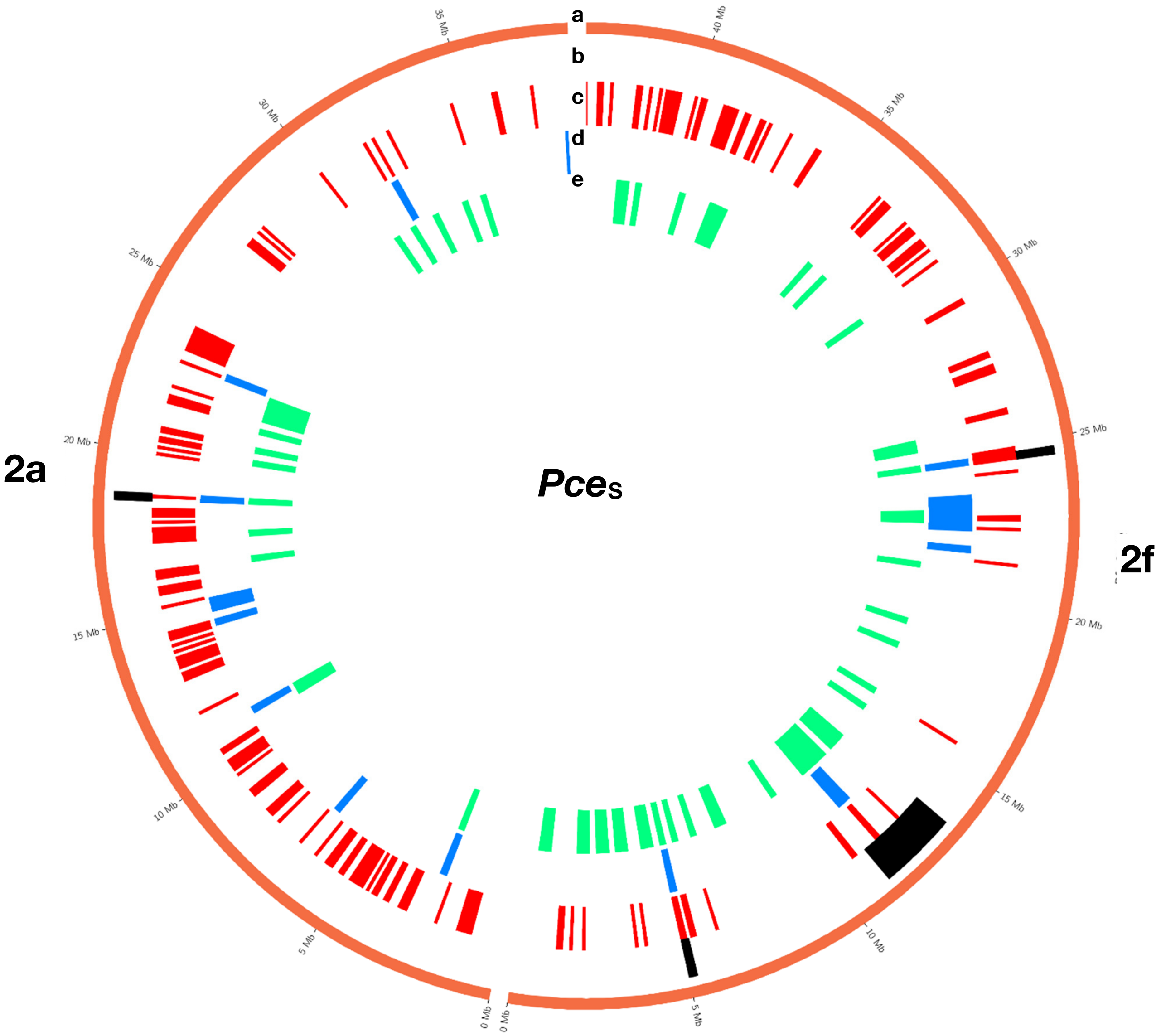


**Continuation of figure S10** Detected regions of homoeologous exchanges in the genome of *P. cerasus* 'Schattenmorelle', pseudomolecule 2.

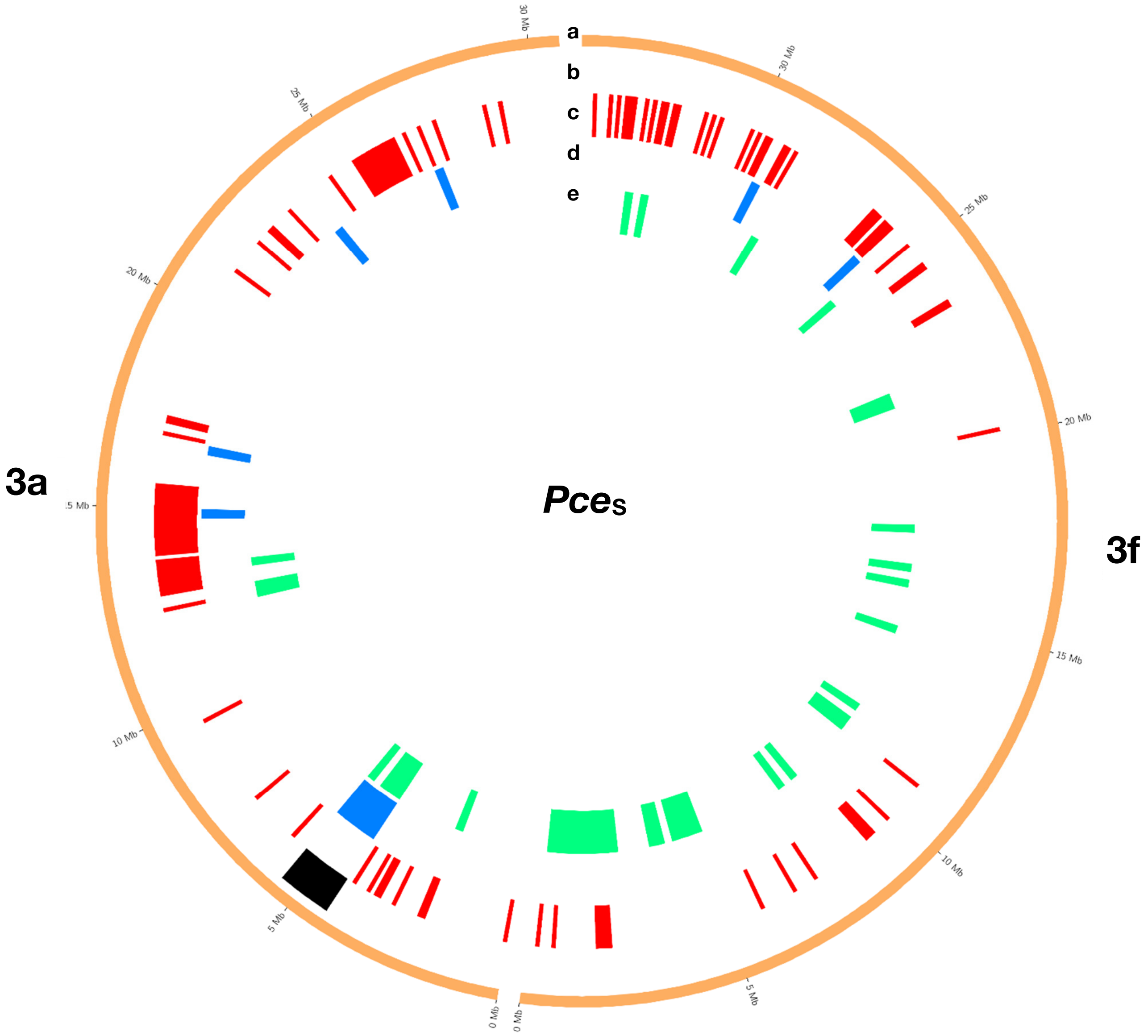


**Continuation of figure S10** Detected regions of homoeologous exchanges in the genome of *P. cerasus* 'Schattenmorelle', pseudomolecule 3.

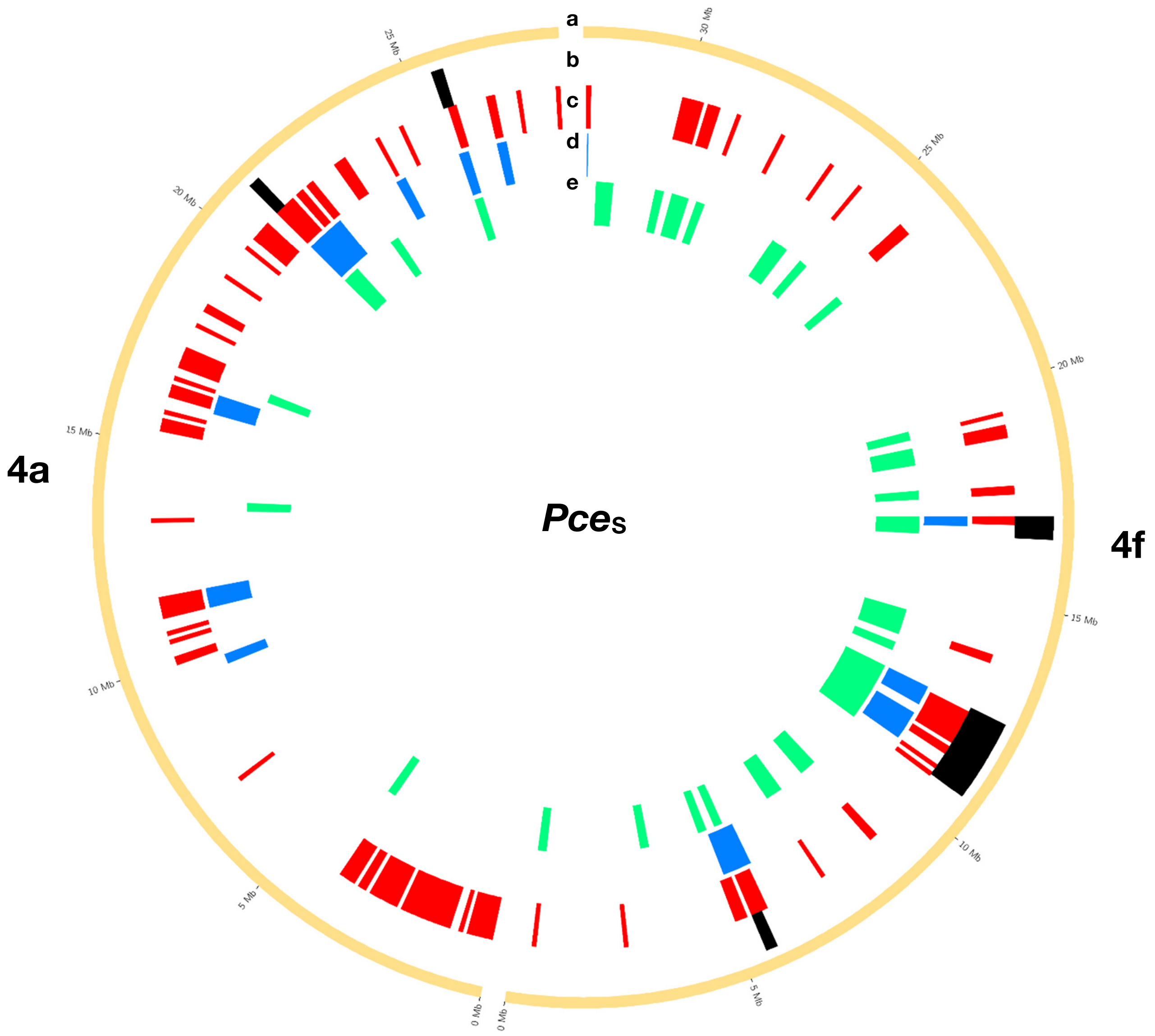


**Continuation of figure S10** Detected regions of homoeologous exchanges in the genome of *P. cerasus* 'Schattenmorelle', pseudomolecule 4.

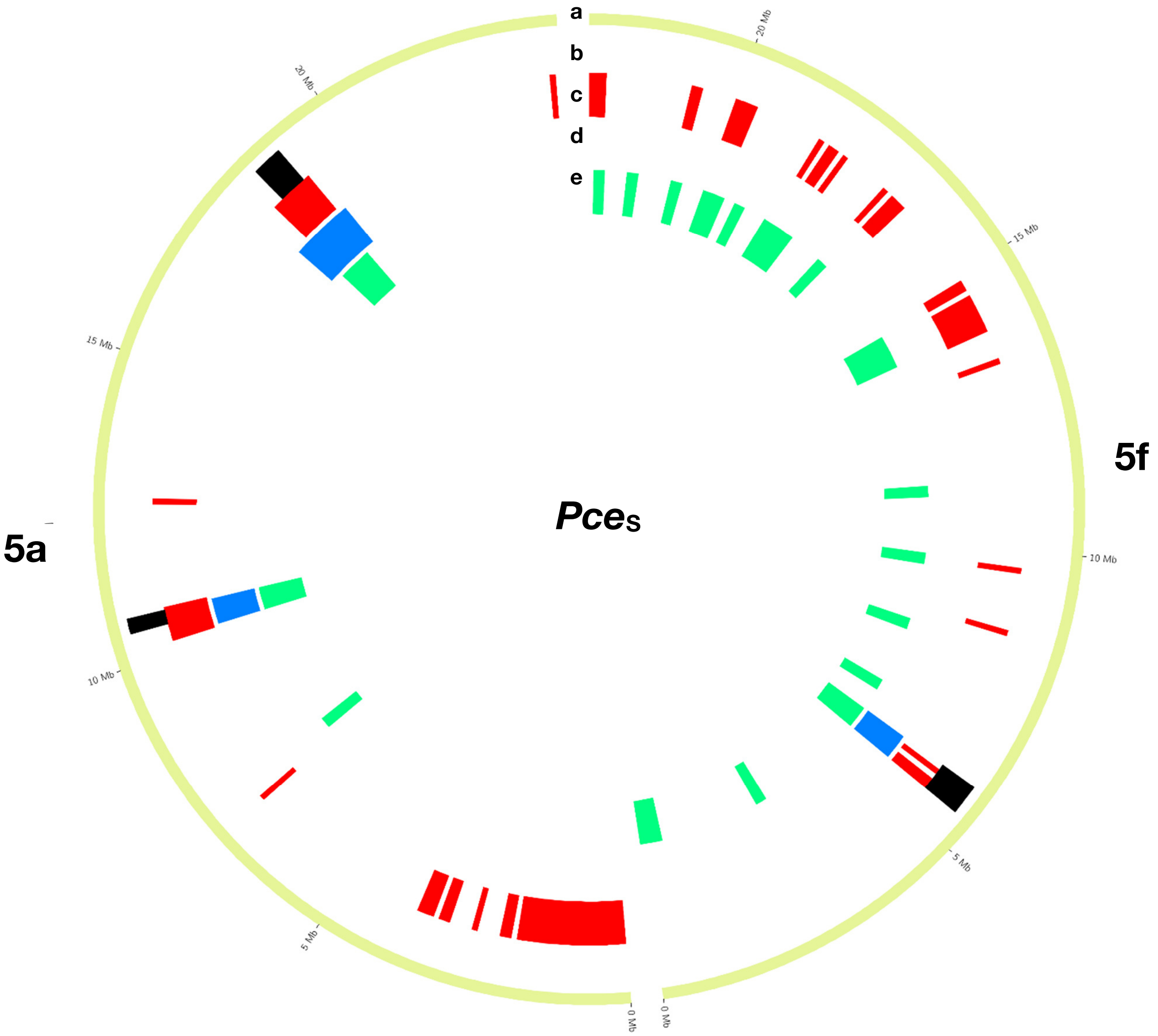


**Continuation of figure S10** Detected regions of homoeologous exchanges in the genome of *P. cerasus* 'Schattenmorelle', pseudomolecule 5.

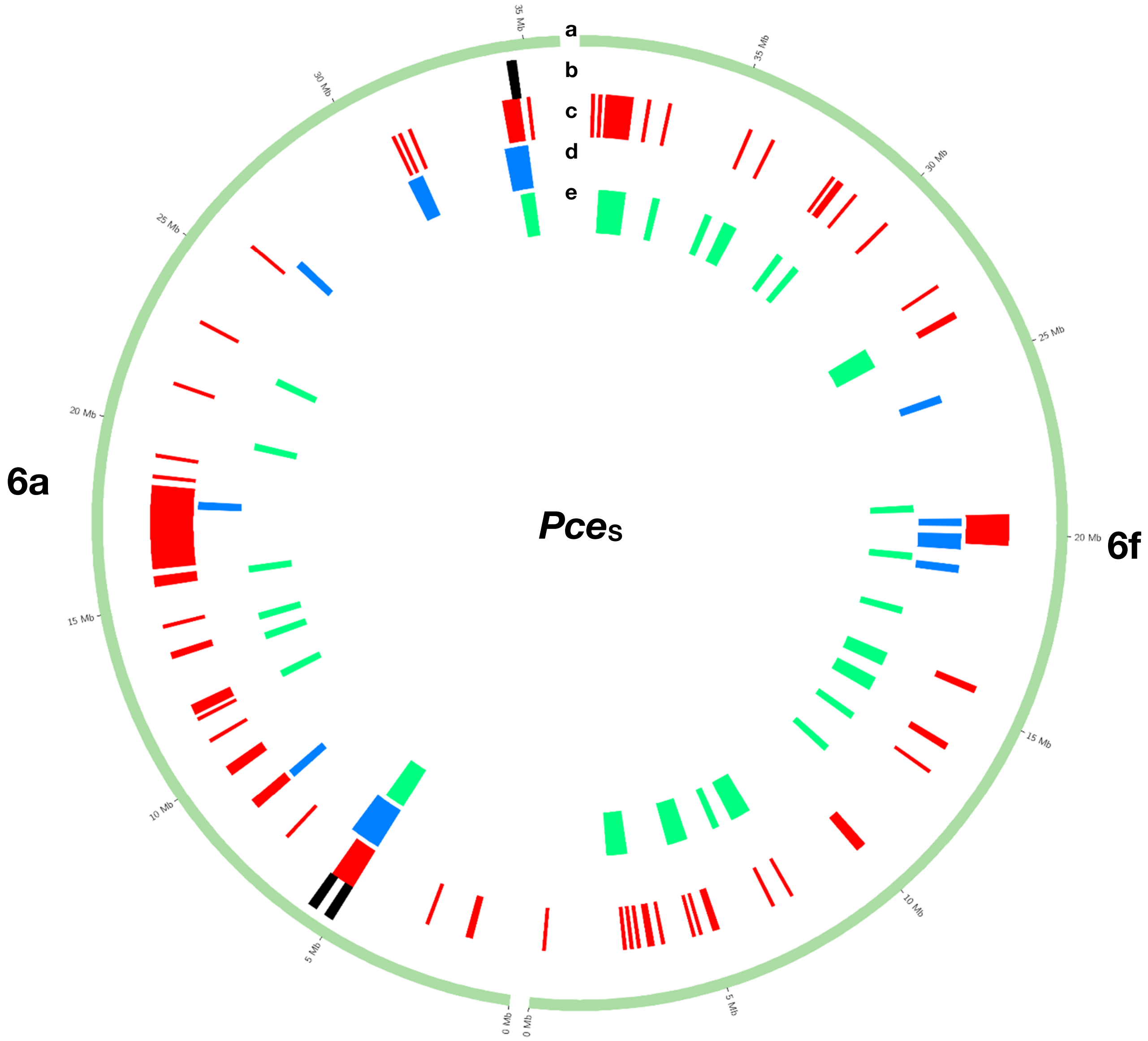


**Continuation of figure S10** Detected regions of homoeologous exchanges in the genome of *P. cerasus* 'Schattenmorelle', pseudomolecule 6.

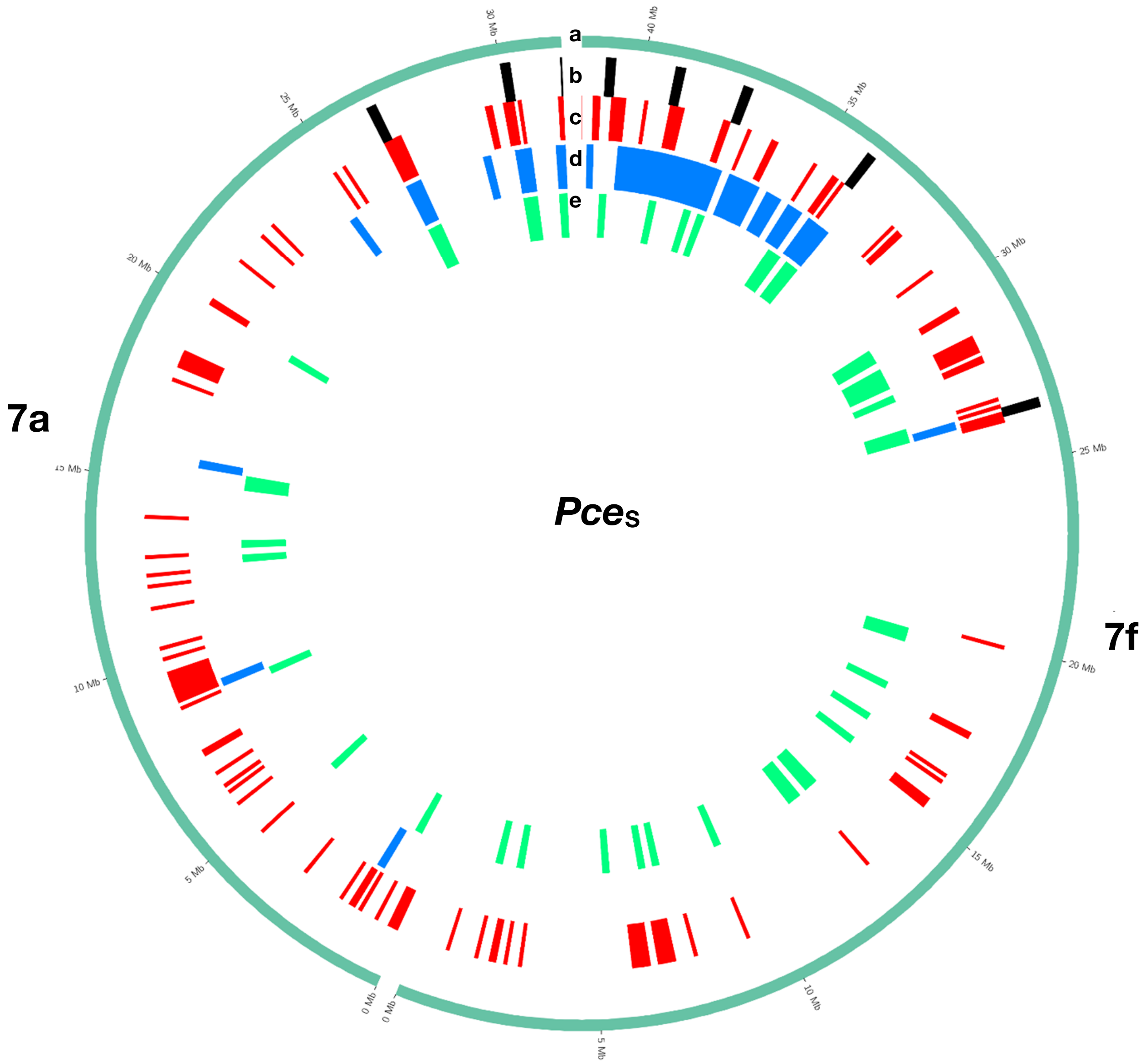


**Continuation of figure S10** Detected regions of homoeologous exchanges in the genome of *P. cerasus* 'Schattenmorelle', pseudomolecule 7.

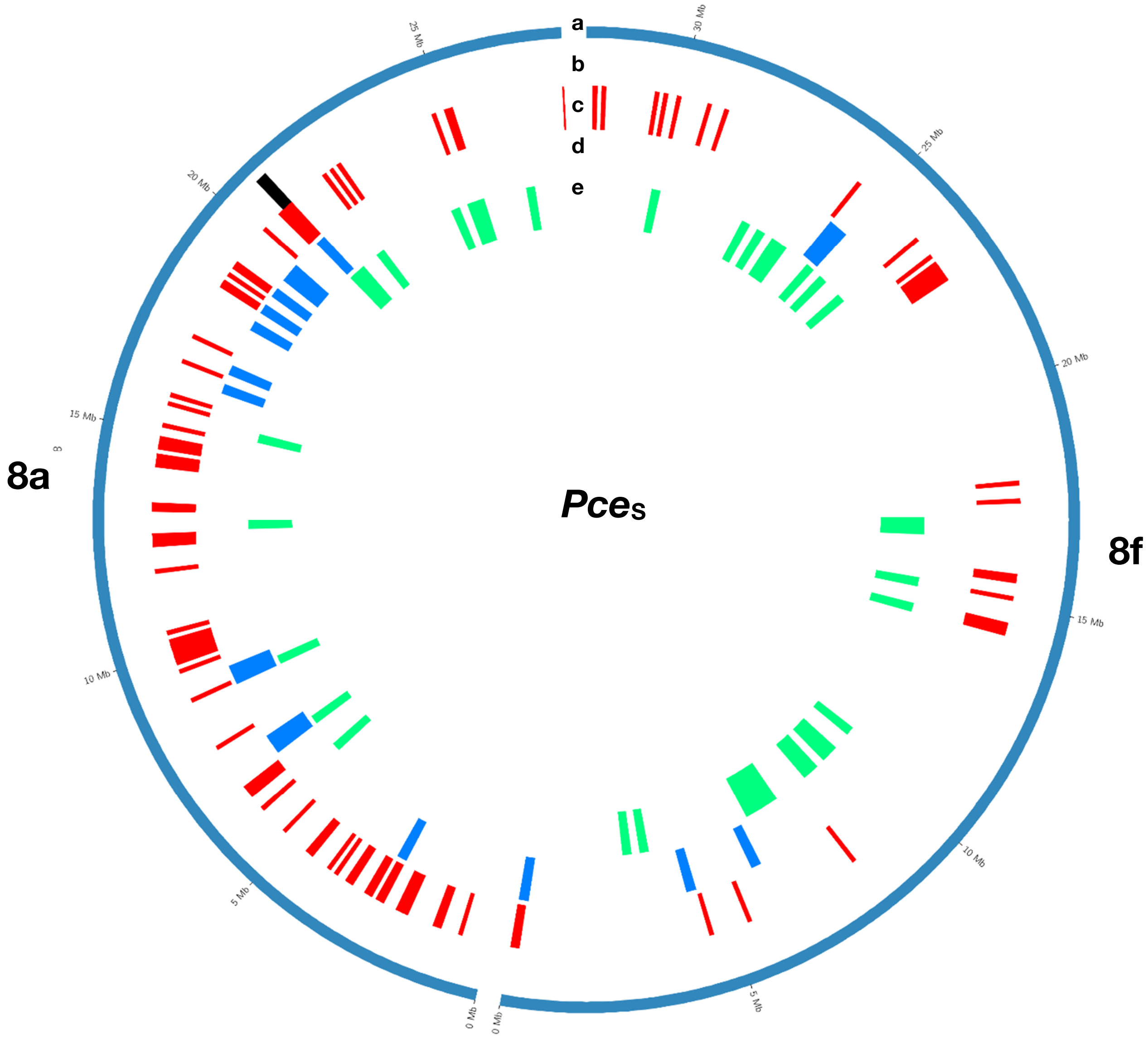


**Continuation of figure S10** Detected regions of homoeologous exchanges in the genome of *P. cerasus* 'Schattenmorelle', pseudomolecule 8.

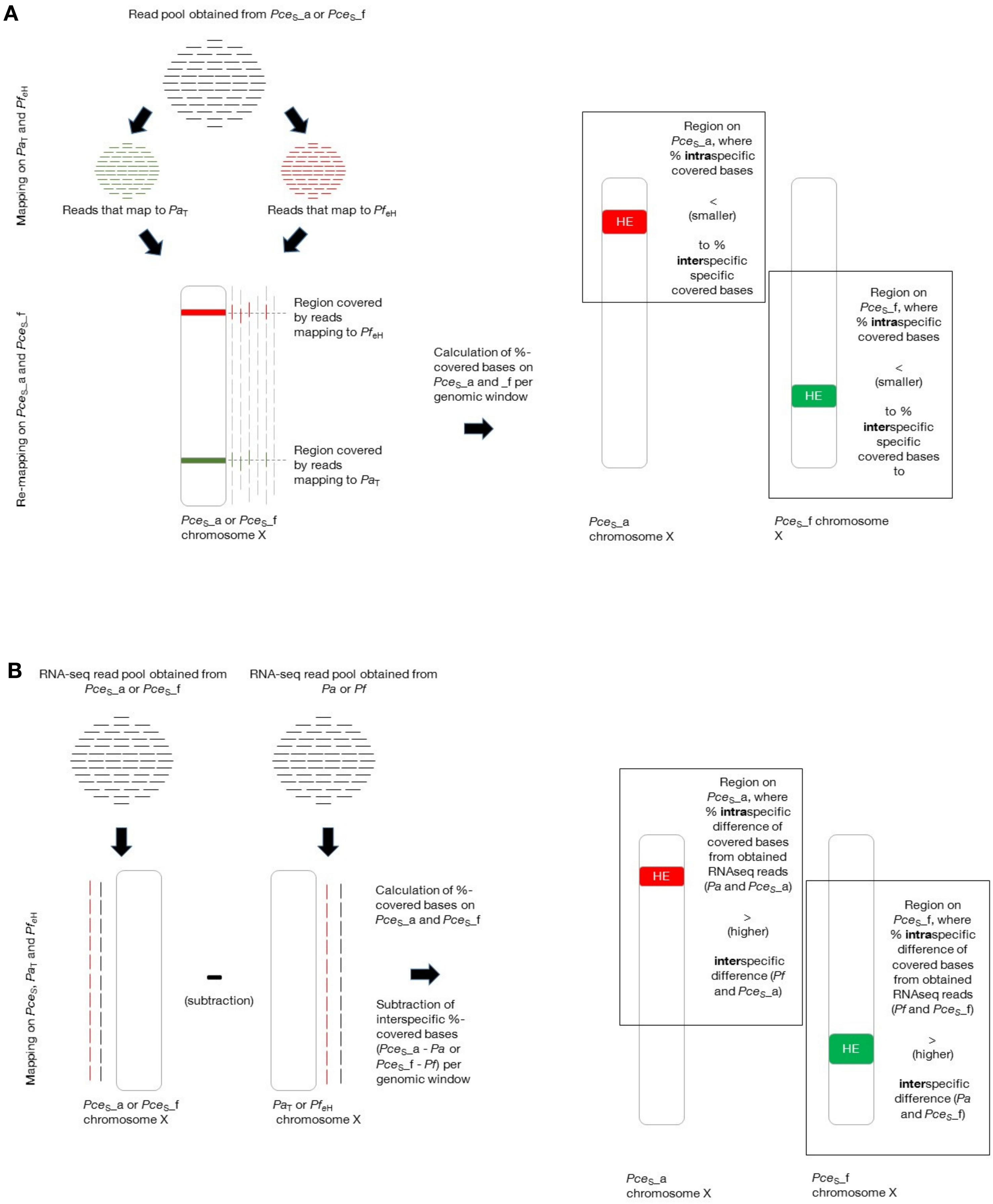
A
Read pool obtained from Pce_S_a or Pce_S_f
Mapping on Pa_T and Pf_eH
Reads that map to Pa_T
Reads that map to Pf_eH
Re-mapping on Pce_S_a and Pce_S_f
Region covered by reads mapping to Pf_eH
Region covered by reads mapping to Pa_T
Pce_S_a or Pce_S_f chromosome X
Calculation of %-covered bases on Pce_S_a and _f per genomic window
Region on Pce_S_a, where % intraspecific covered bases
<
(smaller)
to % interspecific specific covered bases
HE
Region on Pce_S_f, where % intraspecific covered bases
<
(smaller)
to % interspecific specific covered bases to
HE
Pce_S_a chromosome X
Pce_S_f chromosome X
B
RNA-seq read pool obtained from Pce_S_a or Pce_S_f
RNA-seq read pool obtained from Pa or Pf
Mapping on Pce_S, Pa_T and Pf_eH
(subtraction)
Pce_S_a or Pce_S_f chromosome X
Pa_T or Pf_eH chromosome X
Calculation of %-covered bases on Pce_S_a and Pce_S_f
Subtraction of interspecific %-covered bases (Pce_S_a - Pa or Pce_S_f - Pf) per genomic window
Region on Pce_S_a, where % intraspecific difference of covered bases from obtained RNAseq reads (Pa and Pce_S_a)
>
(higher)
interspecific difference (Pf and Pce_S_a)
HE
Region on Pce_S_f, where % intraspecific difference of covered bases from obtained RNAseq reads (Pf and Pce_S_f)
>
(higher)
interspecific difference (Pa and Pce_S_f)
HE
Pce_S_a chromosome X
Pce_S_f chromosome X

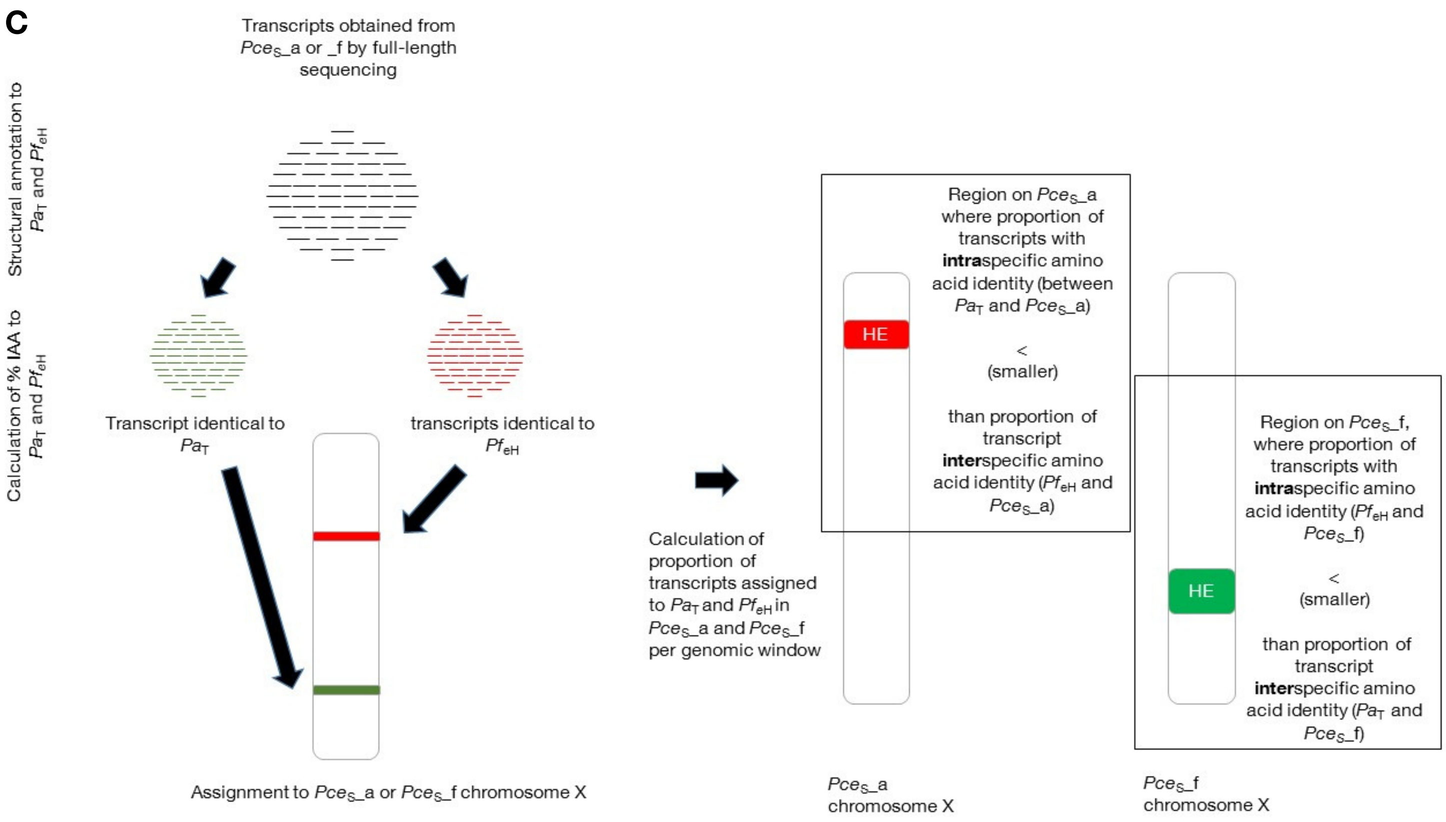


**Figure S11** Schematic summary of the 3 approaches to evaluate homoelogous exchanges (HE) in the *P. cerasus* L. genome sequence of 'Schattenmorelle' by (**A**) genomic reads: the intraspecific % of covered bases from mapped reads ($Pce_S$_a to $Pa_T$, $Pce_S$_f to $Pf_{eH}$) is smaller (<) than the interspecific % of covered bases from mapped reads ($Pce_S$_a to $Pf_{eH}$, $Pce_S$_f to $Pa_T$); (**B**) transcriptomic reads: Intraspecific difference of covered bases from obtained RNAseq reads (% covered bases from *Pa* reads subtracted (-) from % covered bases from $Pce_S$_a, and vice versa for *Pf* and $Pce_S$_f) is higher (>) than the interspecific difference of covered bases from obtained RNAseq reads (% covered bases from *Pf* subtracted from $Pce_S$_a, and vice versa for *Pa* and $Pce_S$_f); (**C**) Identity of amino acids (IAA): were proportion of transcripts with intraspecific amino acid identity ($Pa_T$ and $Pce_S$_a, $Pf_{eH}$ and $Pce_S$_f) smaller (<) than the proportion of transcripts with interspecific amino acid identity ($Pf_{eH}$ and $Pce_S$_a , $Pa_T$ and $Pce_S$_f).

**Table S1** Statistics of different assemblies for *P. cerasus* cv 'Schattenmorelle' (*Pce*$_S$) and the subgenomes *Pce*$_S$_a and *Pce*$_S$_f

| Assembly | Ploidy | Number of contigs* | Contig N50 (kb) | Total contig length (Mb) | Total in scaffolds (Mb) | Total in scaffolds (%) |
|---|---|---|---|---|---|---|
| 20-WGS-PCE.1.0 | 4n | 3750 | 384.68 | 1071.9 | | |
| *Pce*$_S$_a | 2n | 1714 | 328.2 | 473.3 | | |
| *Pce*$_S$_a _purged | 1n | 1058 | 406.2 | 339.3 | | |
| *Pce*$_S$_f | 2n | 2036 | 460.3 | 598.6 | | |
| *Pce*$_S$_f purged | 1n | 1065 | 592.4 | 411.3 | | |
| *Pce*$_S$_a _scaffolded | 1n | 86 | 31,537.40 | 291.7 | 269.00 | 92.2 |
| *Pce*$_S$_f _scaffolded | 1n | 134 | 39,437.03 | 336.8 | 299.47 | 88.9 |

**Table S2** Pseudomolecule statistics for *Pce*S

| Pseudomolecule | Total size (bp) | % |
| --- | --- | --- |
| *Pce*S _a_1.0_chr1 | 52824182 | 9.3 |
| *Pce*S _a _1.0_chr2 | 38212775 | 6.7 |
| *Pce*S _a _1.0_chr3 | 30675746 | 5.4 |
| *Pce*S _a _1.0_chr4 | 28226313 | 5.0 |
| *Pce*S _a _1.0_chr5 | 23885644 | 4.2 |
| *Pce*S _a _1.0_chr6 | 35906979 | 6.3 |
| *Pce*S _a _1.0_chr7 | 31537404 | 5.5 |
| *Pce*S _a _1.0_chr8 | 27735405 | 4.9 |
| *Pce*S _f_1.0_chr1 | 53495658 | 9.4 |
| *Pce*S _f _1.0_chr2 | 43438225 | 7.6 |
| *Pce*S _f _1.0_chr3 | 34378064 | 6.0 |
| *Pce*S _f _1.0_chr4 | 32374258 | 5.7 |
| *Pce*S _f _1.0_chr5 | 22600031 | 4.0 |
| *Pce*S _f _1.0_chr6 | 39437034 | 6.9 |
| *Pce*S _f _1.0_chr7 | 41603557 | 7.3 |
| *Pce*S _f _1.0_chr8 | 32145936 | 5.7 |
| | 568477211 | 100 |

**Table S3** Iso-Seq results

| | No |
|---|---|
| SMRT cells | 2 |
| Circular Consensus Sequencing reads (Raw) | 4,815,872 |
| Circular Consensus Sequencing reads (Filtered) | 4,499,940 |
| Circular Consensus Sequencing reads (Full-Length) | 4,476,054 |
| HQ isoforms | 248,218 |
| LQ isoforms | 480 |

**Table S4** Functional annotation results generated by interproscan using BRAKER & GeMoMa combination of ab-initio and homology-based structural gene annotation and statistics

| Interproscan annotations | No. | *Pce*$_S$_a | *Pce*$_S$_f | Contigs_A | Contigs_F |
|---|---|---|---|---|---|
| Coils | 26427 | 12258 | 11902 | 983 | 1284 |
| Gene3D | 135081 | 63213 | 60649 | 4627 | 6592 |
| Hamap | 2773 | 1323 | 1217 | 108 | 125 |
| PANTHER | 217622 | 100591 | 98613 | 7622 | 10796 |
| Pfam | 155028 | 72373 | 69489 | 5449 | 7717 |
| Phobius | 314975 | 147171 | 139810 | 11215 | 16779 |
| PIRSF | 9105 | 4208 | 4187 | 328 | 382 |
| PRINTS | 85141 | 40737 | 37507 | 2991 | 3906 |
| ProDom | 1125 | 484 | 512 | 59 | 70 |
| ProSitePatterns | 32422 | 15152 | 14560 | 1097 | 1613 |
| ProSiteProfiles | 89962 | 42225 | 40678 | 3050 | 4009 |
| SignalP_EUK | 11628 | 5290 | 5360 | 373 | 605 |
| SMART | 77106 | 36030 | 35078 | 2420 | 3578 |
| SUPERFAMILY | 104068 | 48770 | 46637 | 3666 | 4995 |
| TIGRFAM | 19751 | 9185 | 8780 | 756 | 1030 |
| TMHMM | 99627 | 47068 | 43744 | 3488 | 5327 |
| Sum | **1381841** | 646078 | 530906 | 48232 | 156625 |

| | | | | | |
|---|---|---|---|---|---|
| Genes | | | | | |
| total | 60123 | 26947 | 27876 | 2122 | 3178 |
| annotated | 56047 | 25152 | 25960 | 1988 | 2947 |
| Transcripts | | | | | |
| total | 107508 | 49698 | 48576 | 3799 | 5435 |
| annotated | 103243 | 47807 | 46585 | 3657 | 5194 |
| annotated GO | 71870 | 33554 | 32284 | 2561 | 3471 |
| annotated pathways | 9114 | 4317 | 4115 | 299 | 383 |
| Mean length (bp) | 3580 | 3864 | 3808 | 4031 | 3565 |
| Mean length of predicted proteins | 469 | 469 | 460 | 466 | 450 |

**Table S5** Mapping of marker sequences to *Prunus cerasus* 'Schattenmorelle' (*Pce*$_S$) genome

| Genetic map | No. of marker input | No. of mapped markers |
|---|---|---|
| M172x25-F1 | 1795 | 1514 |
| US-F1 | 1787 | 1506 |
| 25x25-F1 | 1807 | 1510 |
| Montx25-F1 | 1795 | 1517 |
| RE-F1 | 1856 | 1509 |

**Table S6** Position of inversions within the subgenomes of *Pce*$_S$ indicated by collinearity of marker positions.

| Chr. *Pce*$_S$ vs. *Pce*$_S$ | position of inversion in Mbp | Chr. *Pce*$_S$_a vs. *Pa*$_T$ | position of inversion in Mbp | Chr. *Pce*$_S$_f vs. *Pf*$_{eH}$ | position of inversion in Mbp |
|---|---|---|---|---|---|
| Chr1a | 9-11, 42-43, 51-52 | Chr1a | 49-52 | Chr1f | 50-53 |
| Chr1f | 11-12, 43-44, 52-53 | Chr1a | 56-62 | Chr1f | 61-65 |
| Chr2a | - | Chr2a | - | Chr2f | - |
| Chr2f | - | Chr2a | - | Chr2f | - |
| Chr3a | 30-32 | Chr3a | - | Chr3f | 7-9, 33-34 |
| Chr3f | 33-34 | Chr3a | - | Chr3f | 8-9, 37-39 |
| Chr4a | 1-7 | Chr4a | - | Chr4f | 1-7 |
| Chr4f | 1-7 | Chr4a | - | Chr4f | 1-7 |
| Chr5a | 8-9, 15-17, 22-25 | Chr5a | - | Chr5f | 6-7, 15-17, 19-22 |
| Chr5f | 6-7, 15-17, 19-22 | Chr5a | - | Chr5f | 7-8, 20-21, 28-31 |
| Chr6a | 5-6 | Chr6a | - | Chr6f | 5-7 |
| Chr6f | 5-7 | Chr6a | - | Chr6f | 6-8 |
| Chr7a | 1-7, 29-32 | Chr7a | 1-7, 17-18, 29-32 | Chr7f | 10-15, 38-42 |
| Chr7f | 10-15, 38-42 | Chr7a | 1-5, 17-19, 31-32 | Chr7f | 1-7, 32-37 |
| Chr8a | 1-9, 26-28 | Chr8a | 1-9 | Chr8f | 1-15 |
| Chr8f | 1-15, 30-32 | Chr8a | 1-12 | Chr8f | 1-13 |

**Note S1** Access to the assembly hub for genome and annotation visualization at UCSC Genome Browser.

An assembly hub for genome and annotation visualization is permantly hosted at http://bioinf.uni-greifswald.de/private-hubs/pcer/hub.txt . To connect this assembly hub to the UCSC Genome Browser, go to https://genome.ucsc.edu, click on "My Data" -> "Track Hubs" -> select one of the mirrors-> "My Hubs" -> insert the link.

**Note S2** Calculation of the 70% quantile from IAA.

For each reference species, the best iAA per transcript was determined. Midpoints of all transcripts were calculated. Windows of size 0.5Mb centered at these midpoints were used to determine the 70% quantile of the iAA values. Therefor all transcripts with midpoint in this windows were collected. The 70% quantile was computed using the corresponding iAA values and ignoring missing values. The 70% quantile was plotted in a graph against the two subgenomes $Pce_S$_a and $Pce_S$_f of *P. cerasus* cv 'Schattenmorelle'. A higher IAA value of proteins from *P. avium* cv 'Tieton' ($Pa_T$) in comparison to proteins from *P. fruticosa* ecotype Hármashatárhegy ($Pf_{eH}$) is expected for $Pce_S$_a and vice versa. An example is given below (Example plot 1). The blue line represents the 70% quantile value obtained from the comparison between $Pce_S$_a and $Pa_T$. Due to the relation of both species, the blue line is nearly 1, except for some regions. The red line represents the comparison between $Pce_S$_f and $Pf_{eH.}$ The species are not related which is indicated by a lower 70% quantile value. In case of a translocation between $Pce_S$_a and $Pce_S$_f, the values of the red line will increase whereas the value of the blue line will decrease. An example is given in Example plot 2. Note S2 contains 8 plots were the 70% quantile values of IAA obtained from comparisons between *P. avium* 'Tieton' ($Pa_T$) and *P. fruticosa* ecotype Hármashatárhegy ($Pf_{eH}$) with *P. cerasus* 'Schattenmorelle' ($Pce_S$) were plotted against the chromosomes of subgenome $Pce_S$_a. Additionally, eight plots were the 70% quantile values of IAA obtained from comparisons between $Pa_T$ and $Pf_{eH}$ with $Pce_S$ were plotted against the chromosomes of subgenome $Pce_S$_f. The same was calculated for the 14 reference species used in the final annotation procedure.

Comparison of 70%-quantile of IAA between transcripts from *P. cerasus* cv 'Schattenmorelle' assigned to *P. fruticosa* ecotype Hármashatárhegy (Prufru, red) or *P. avium* cv 'Tieton' (Pruavi, blue) for each chromosome.

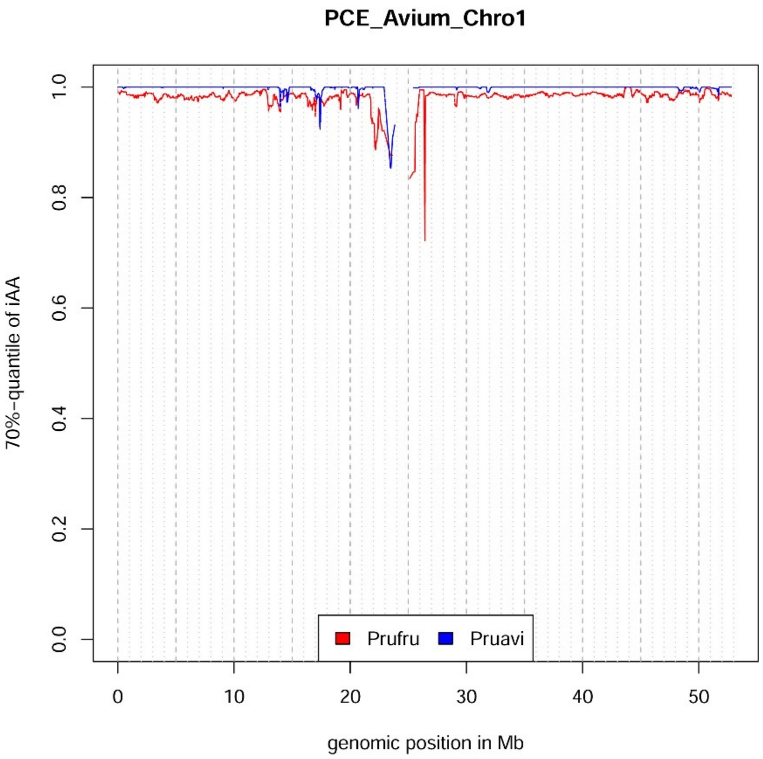


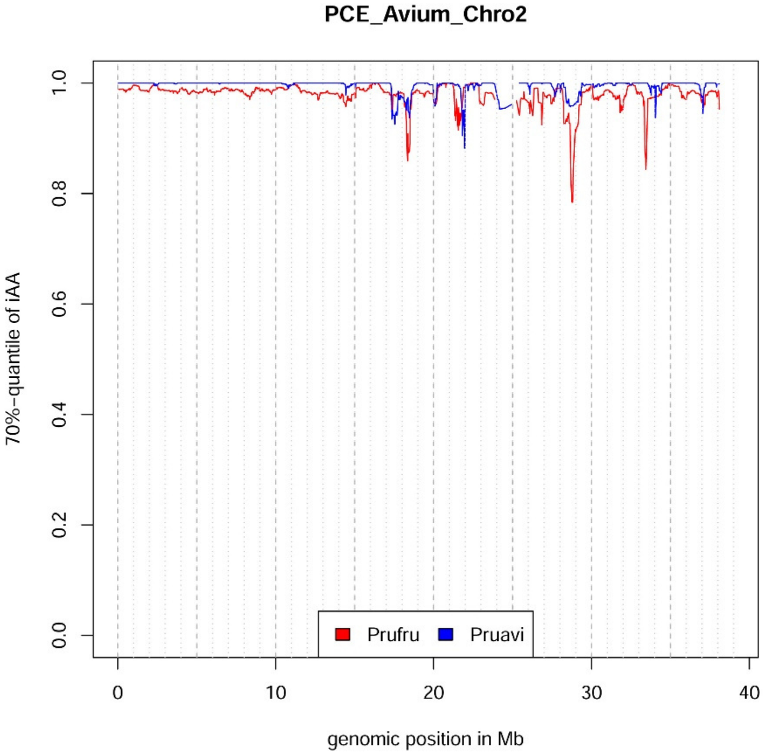

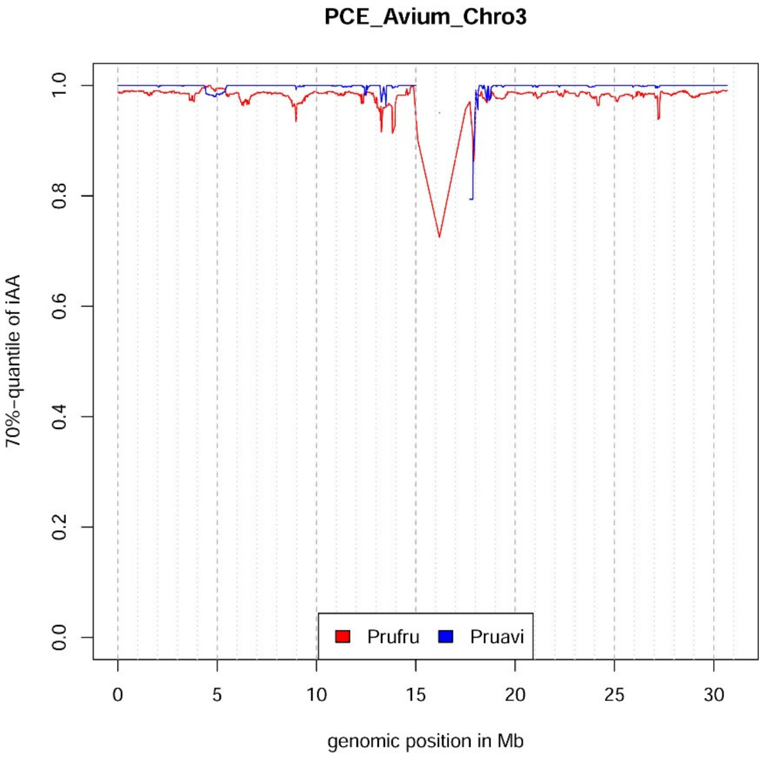
PCE_Avium_Chro3
70%-quantile of iAA
1.0
0.8
0.6
0.4
0.2
0.0
Prufru
Pruavi
0
5
10
15
20
25
30
genomic position in Mb

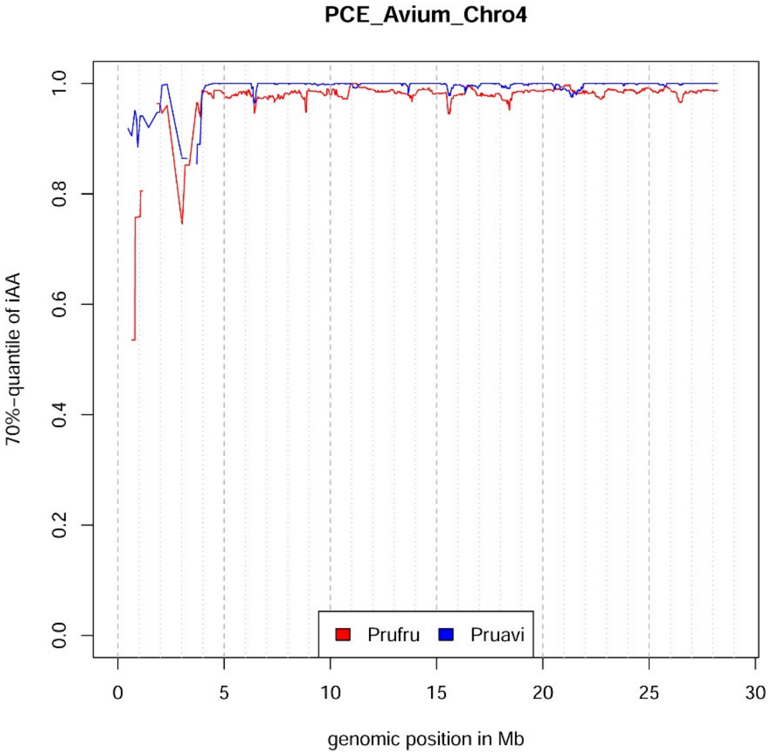
PCE_Avium_Chro4
70%-quantile of iAA
1.0
0.8
0.6
0.4
0.2
0.0
Prufru
Pruavi
0
5
10
15
20
25
30
genomic position in Mb

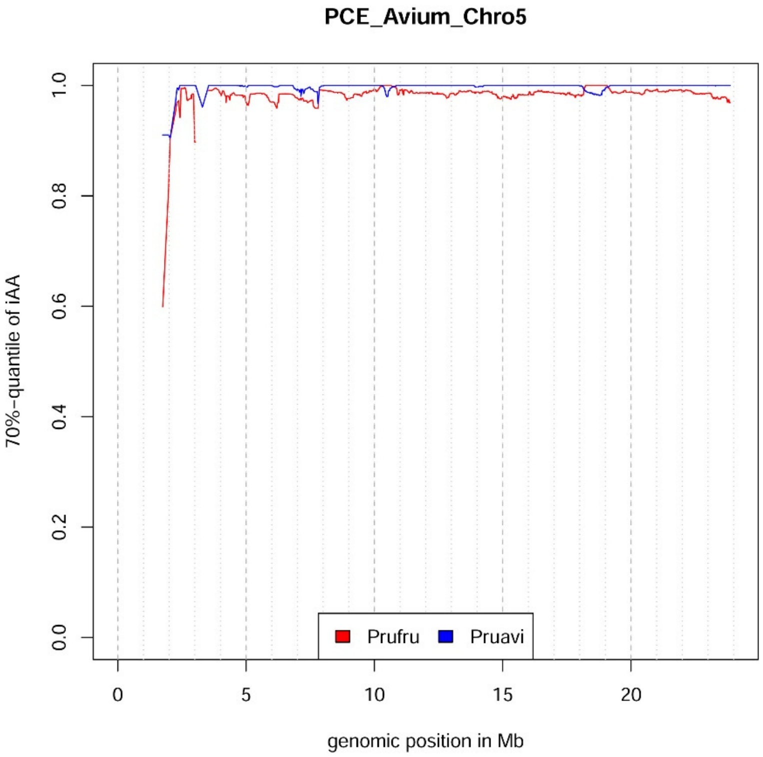
PCE_Avium_Chro5
70%-quantile of iAA
0.0
0.2
0.4
0.6
0.8
1.0
0
5
10
15
20
genomic position in Mb
Prufru
Pruavi

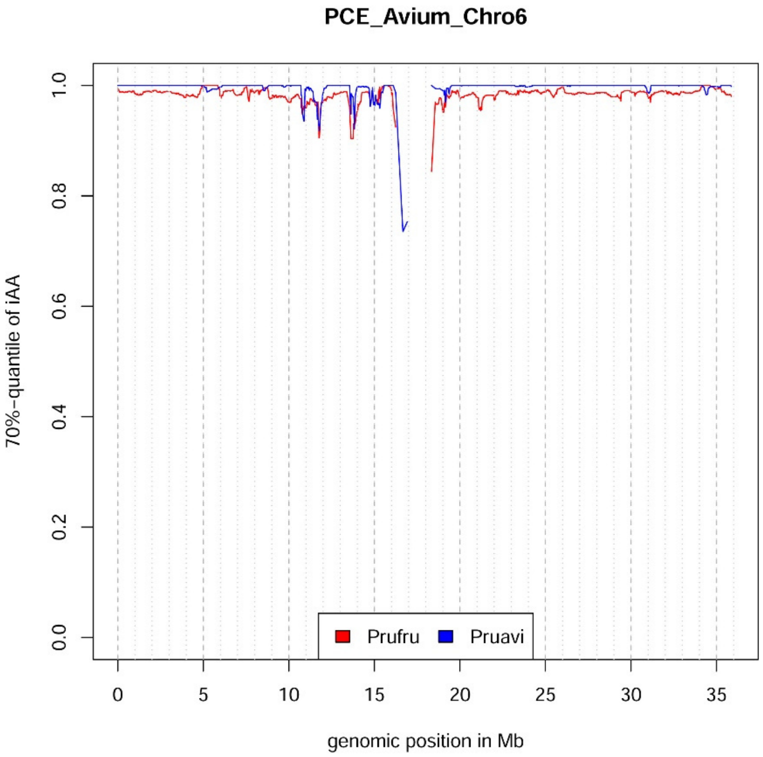
PCE_Avium_Chro6
70%-quantile of iAA
0.0
0.2
0.4
0.6
0.8
1.0
0
5
10
15
20
25
30
35
genomic position in Mb
Prufru
Pruavi

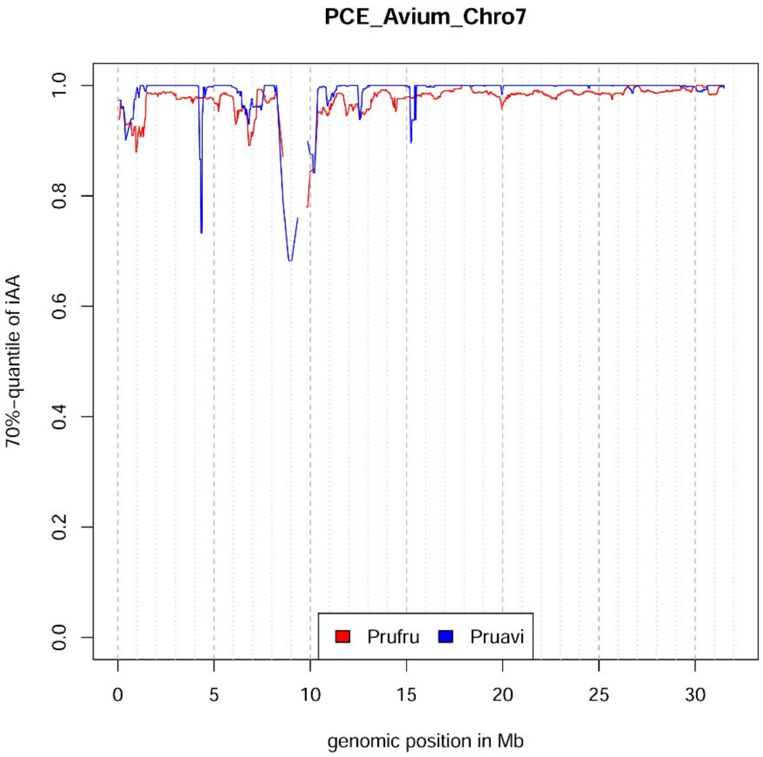

PCE_Avium_Chro7
70%-quantile of iAA
0.0
0.2
0.4
0.6
0.8
1.0
0
5
10
15
20
25
30
genomic position in Mb
Prufru
Pruavi


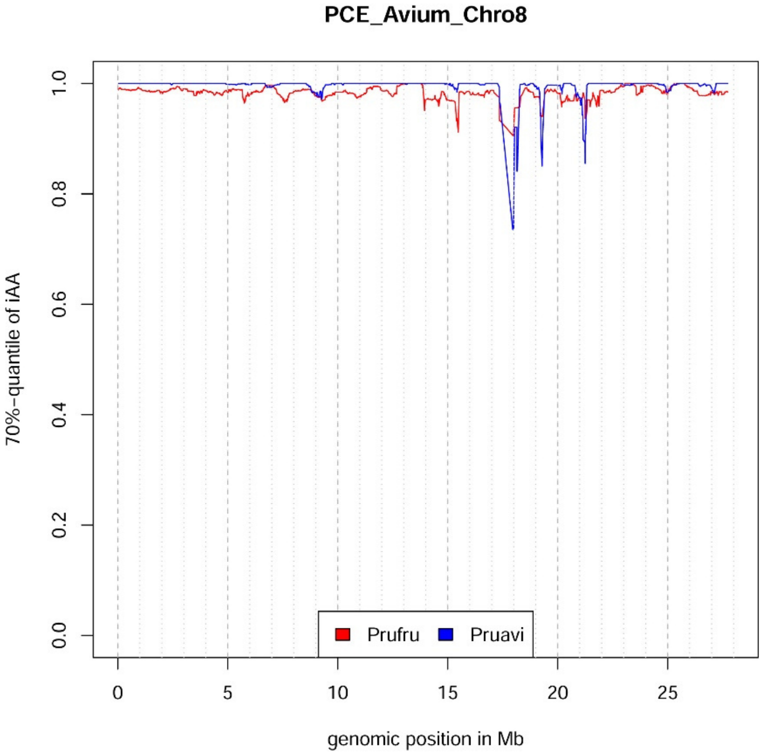

PCE_Avium_Chro8
70%-quantile of iAA
0.0
0.2
0.4
0.6
0.8
1.0
0
5
10
15
20
25
genomic position in Mb
Prufru
Pruavi

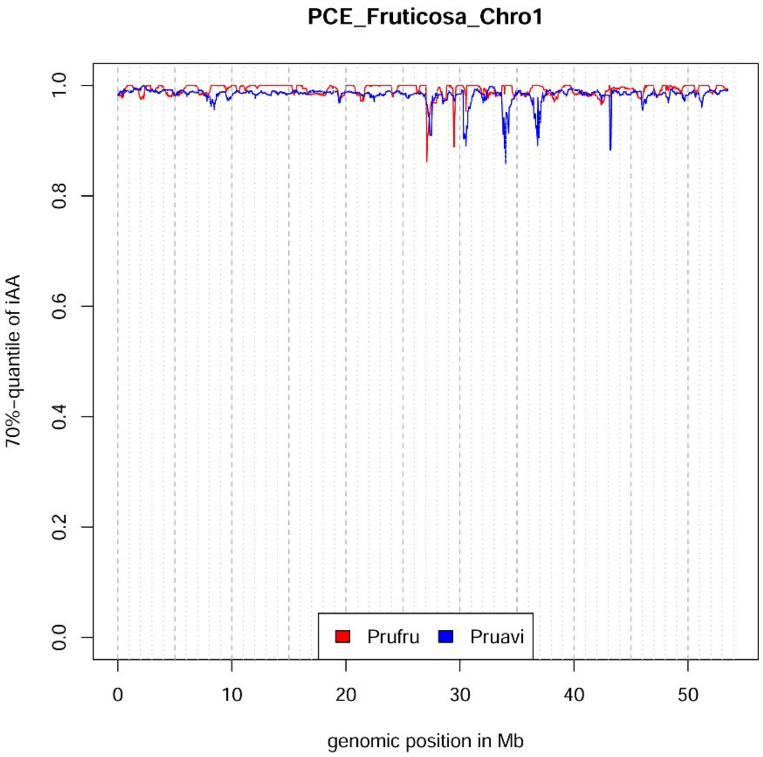
PCE_Fruticosa_Chro1
70%-quantile of iAA
1.0
0.8
0.6
0.4
0.2
0.0
Prufru
Pruavi
0
10
20
30
40
50
genomic position in Mb

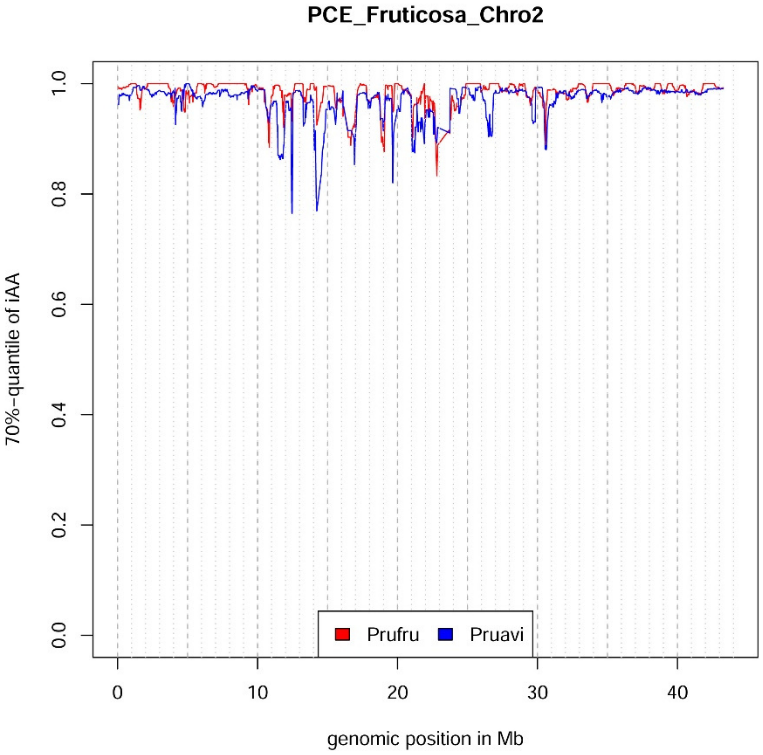
PCE_Fruticosa_Chro2
70%-quantile of iAA
1.0
0.8
0.6
0.4
0.2
0.0
Prufru
Pruavi
0
10
20
30
40
genomic position in Mb

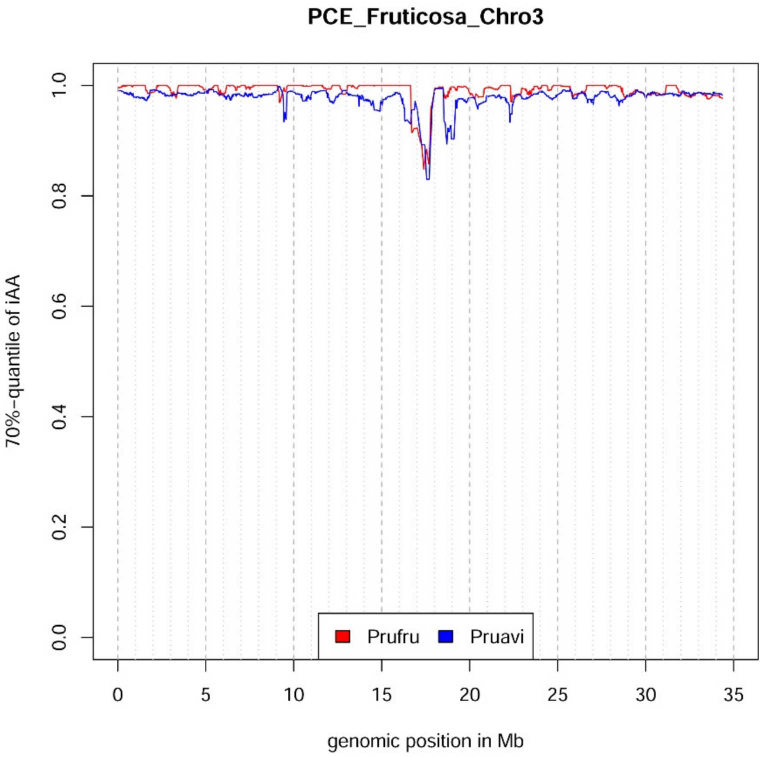
PCE_Fruticosa_Chro3
70%-quantile of iAA
1.0
0.8
0.6
0.4
0.2
0.0
Prufru
Pruavi
0
5
10
15
20
25
30
35
genomic position in Mb

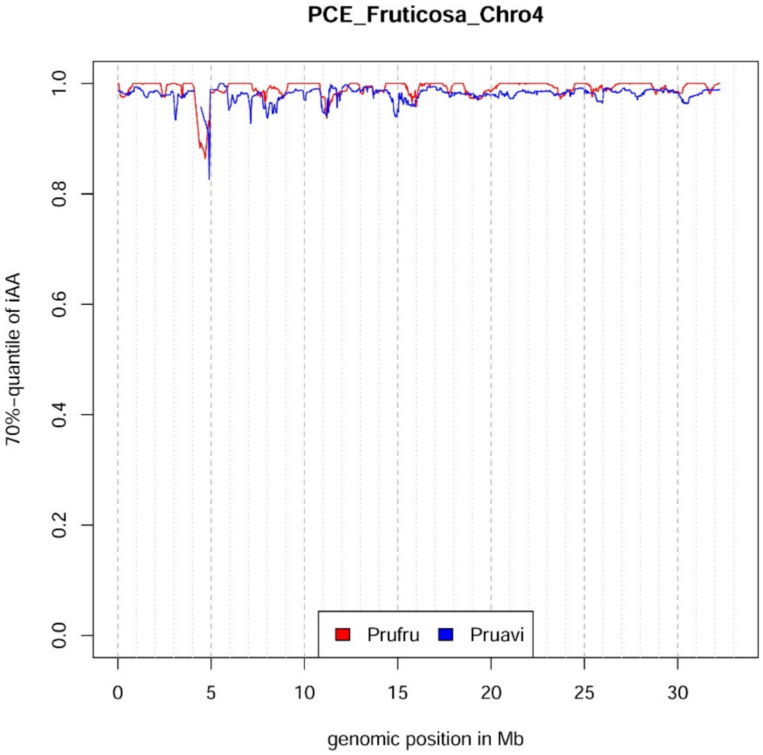
PCE_Fruticosa_Chro4
70%-quantile of iAA
1.0
0.8
0.6
0.4
0.2
0.0
Prufru
Pruavi
0
5
10
15
20
25
30
genomic position in Mb

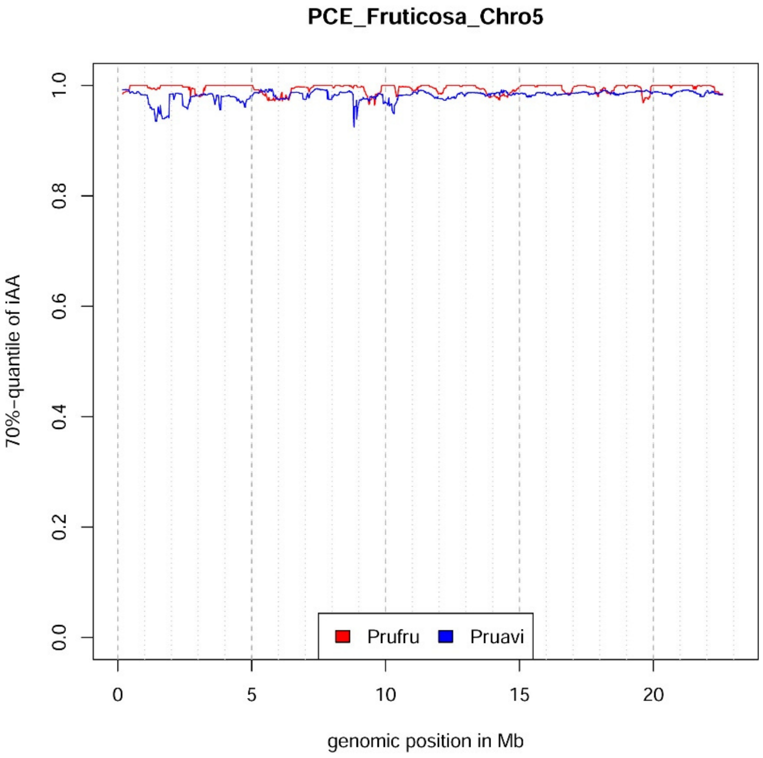
PCE_Fruticosa_Chro5
70%-quantile of iAA
0.0
0.2
0.4
0.6
0.8
1.0
0
5
10
15
20
genomic position in Mb
Prufru
Pruavi

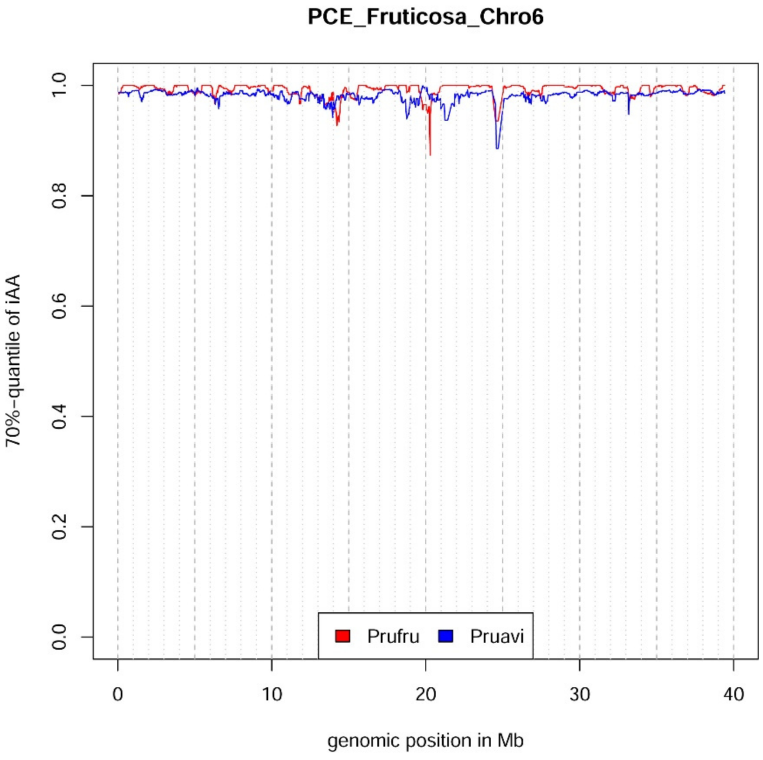
PCE_Fruticosa_Chro6
70%-quantile of iAA
0.0
0.2
0.4
0.6
0.8
1.0
0
10
20
30
40
genomic position in Mb
Prufru
Pruavi

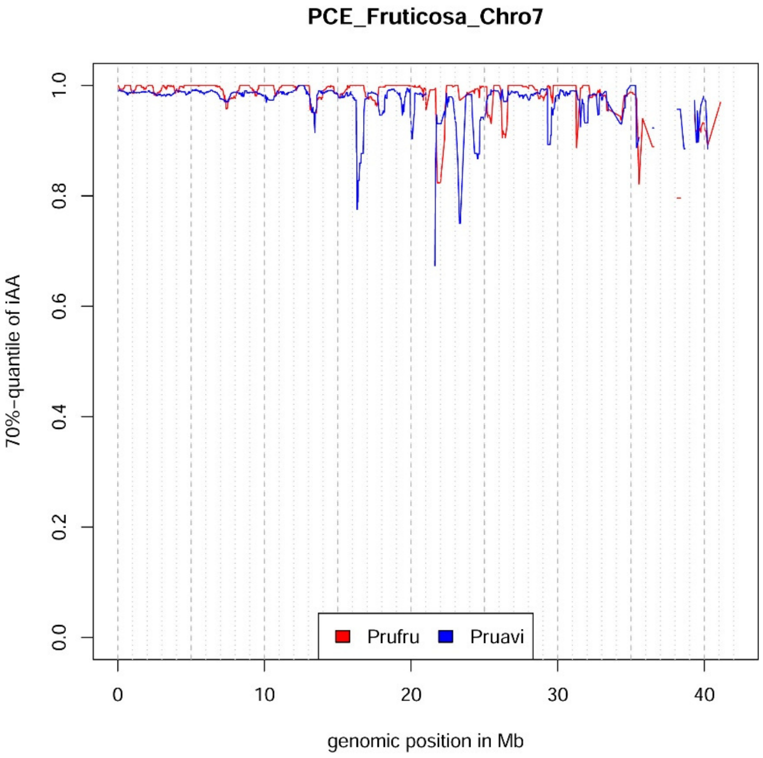

PCE_Fruticosa_Chro7
70%-quantile of iAA
genomic position in Mb
Prufru
Pruavi


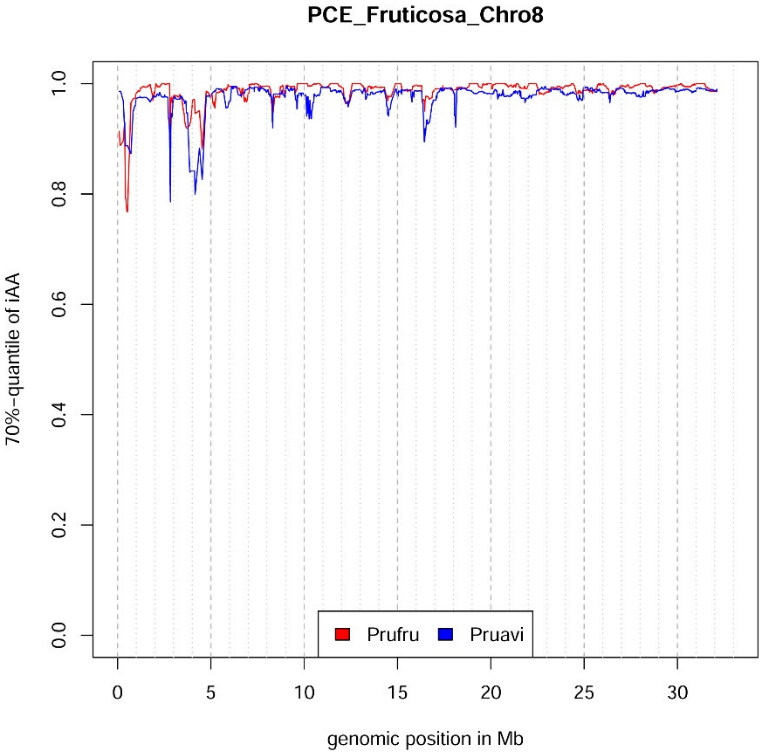

PCE_Fruticosa_Chro8
70%-quantile of iAA
genomic position in Mb
Prufru
Pruavi

Comparison of 70%-quantile of IAA between transcripts from *P. cerasus* cv 'Schattenmorelle' assigned to *P. fruticosa* ecotype Hármashatárhegy (Prufru, red), *P. avium* cv 'Tieton' (Pruavi, blue), *P. yedonensis* (Pyed, green), *P. domestica* (Pd, brown), *P. armeniaca* (Par, orange), *P. persica* (Pp, black), *Pyrus communis* (Pyrco, rose pink), *Populus trichocarpa* (Poptri, *yellow*), *Vitis vinifera* (Vv, green-blue), *Arabidopsis thaliana* (At, pink), *Malus domestica* (Md, lime pink), *Fragaria vesca* (Fraves, hot pink), *Rubus occidentalis* (Rubocc, mint-green), for each chromosome.

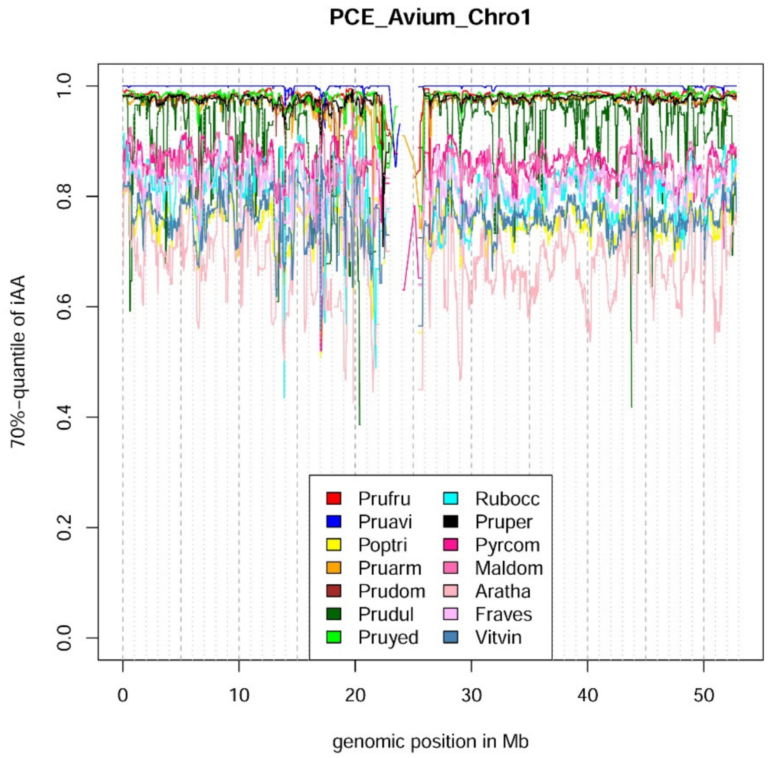


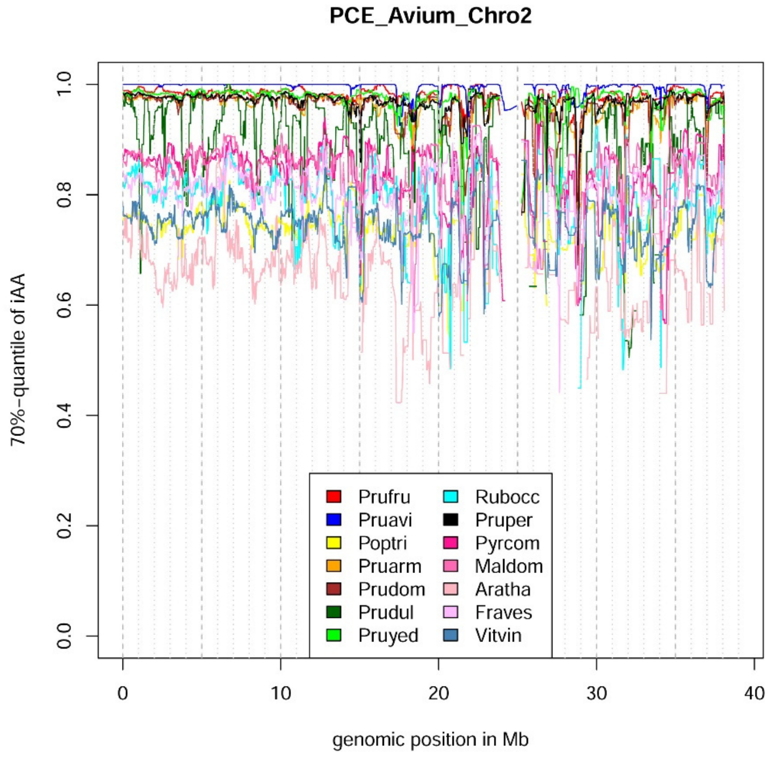

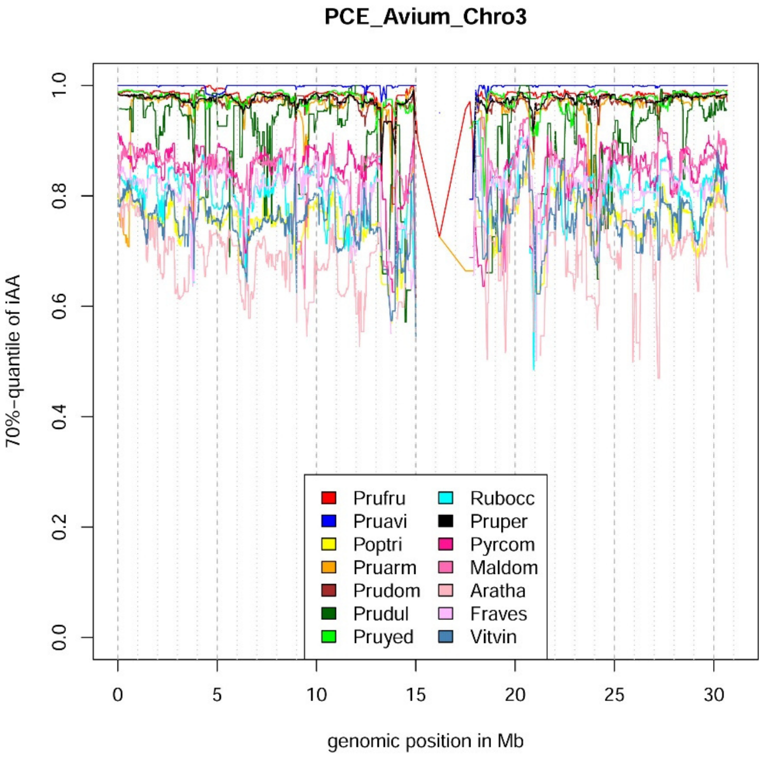
PCE_Avium_Chro3
70%-quantile of iAA
0.0
0.2
0.4
0.6
0.8
1.0
0
5
10
15
20
25
30
genomic position in Mb
Prufru
Pruavi
Poptri
Pruarm
Prudom
Prudul
Pruyed
Rubocc
Pruper
Pyrcom
Maldom
Aratha
Fraves
Vitvin

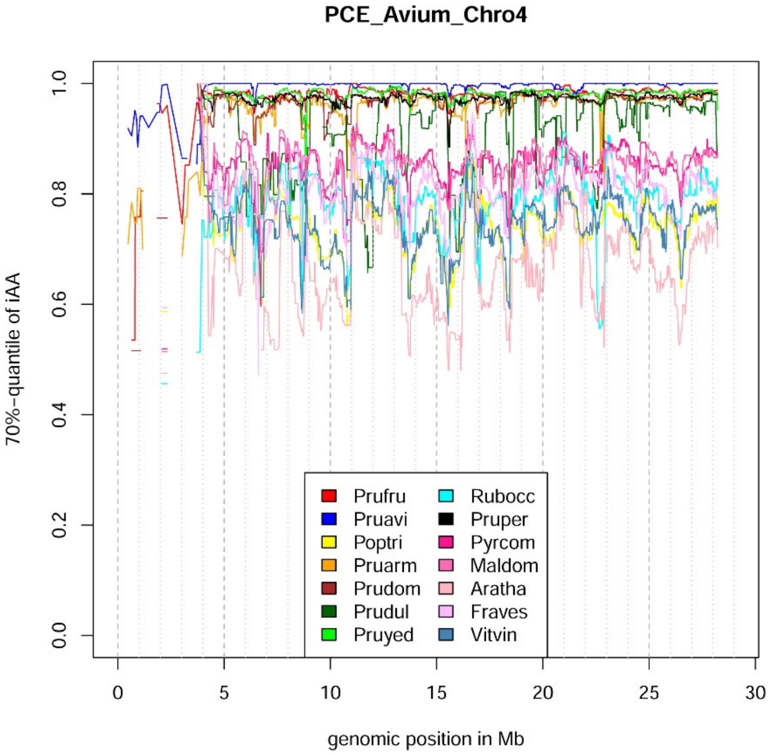
PCE_Avium_Chro4
70%-quantile of iAA
0.0
0.2
0.4
0.6
0.8
1.0
0
5
10
15
20
25
30
genomic position in Mb
Prufru
Pruavi
Poptri
Pruarm
Prudom
Prudul
Pruyed
Rubocc
Pruper
Pyrcom
Maldom
Aratha
Fraves
Vitvin

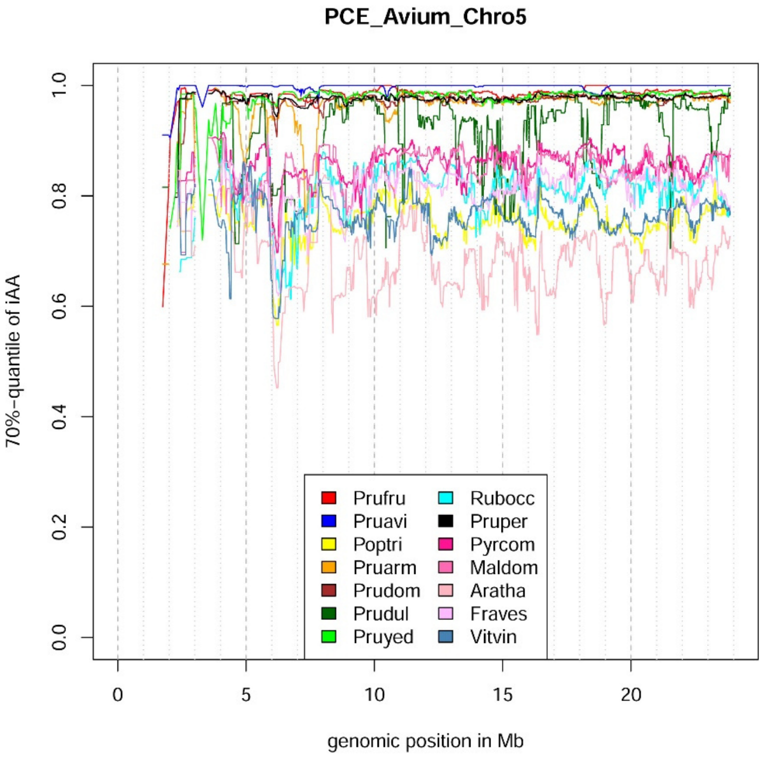
PCE_Avium_Chro5
70%-quantile of iAA
0.0
0.2
0.4
0.6
0.8
1.0
0
5
10
15
20
genomic position in Mb
Prufru
Pruavi
Poptri
Pruarm
Prudom
Prudul
Pruyed
Rubocc
Pruper
Pyrcom
Maldom
Aratha
Fraves
Vitvin

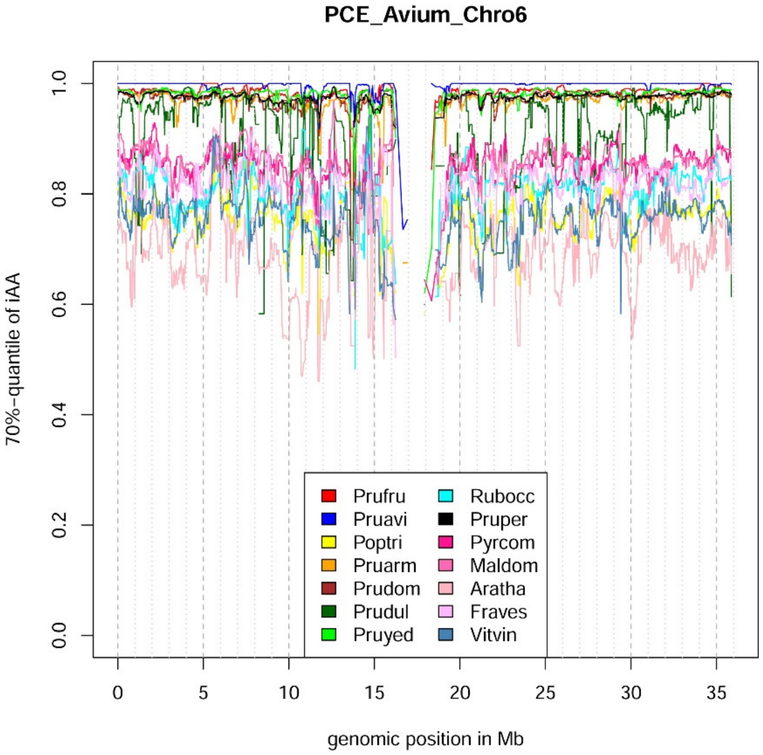
PCE_Avium_Chro6
70%-quantile of iAA
0.0
0.2
0.4
0.6
0.8
1.0
0
5
10
15
20
25
30
35
genomic position in Mb
Prufru
Pruavi
Poptri
Pruarm
Prudom
Prudul
Pruyed
Rubocc
Pruper
Pyrcom
Maldom
Aratha
Fraves
Vitvin

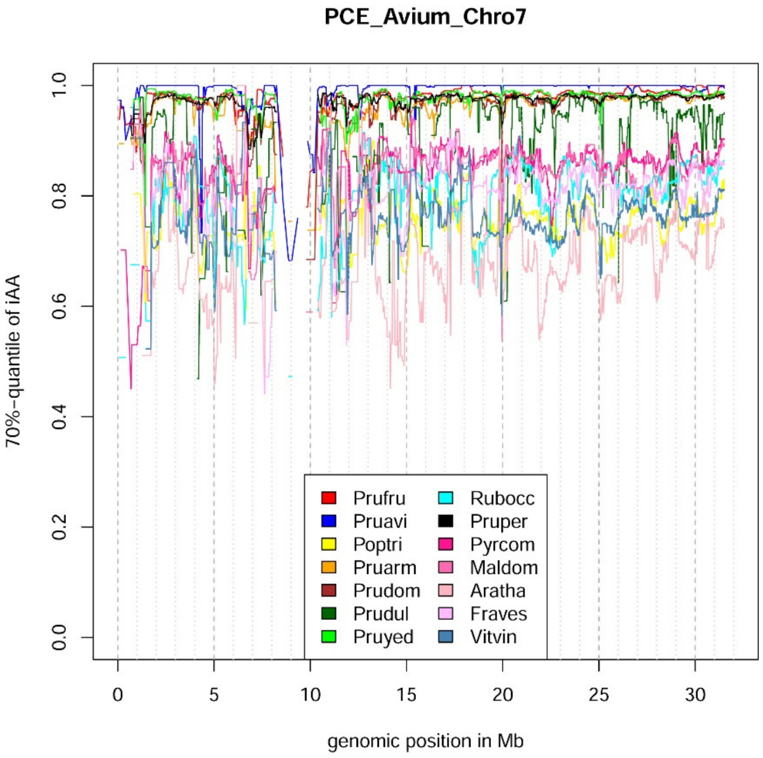
PCE_Avium_Chro7
70%-quantile of iAA
0.0
0.2
0.4
0.6
0.8
1.0
0
5
10
15
20
25
30
genomic position in Mb
Prufru
Pruavi
Poptri
Pruarm
Prudom
Prudul
Pruyed
Rubocc
Pruper
Pyrcom
Maldom
Aratha
Fraves
Vitvin

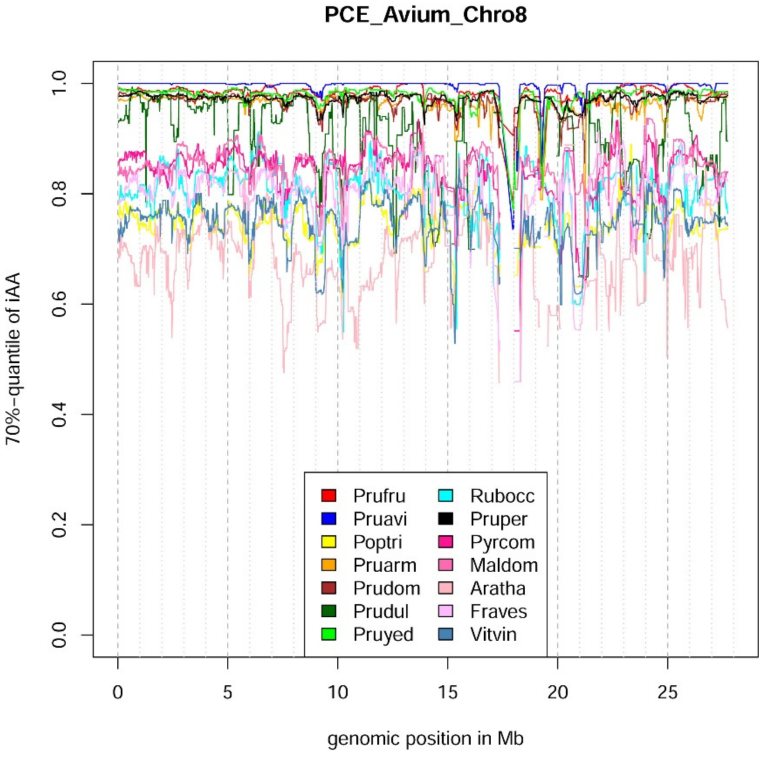
PCE_Avium_Chro8
70%-quantile of iAA
0.0
0.2
0.4
0.6
0.8
1.0
0
5
10
15
20
25
genomic position in Mb
Prufru
Pruavi
Poptri
Pruarm
Prudom
Prudul
Pruyed
Rubocc
Pruper
Pyrcom
Maldom
Aratha
Fraves
Vitvin

#### PCE_Fruticosa_Chro1

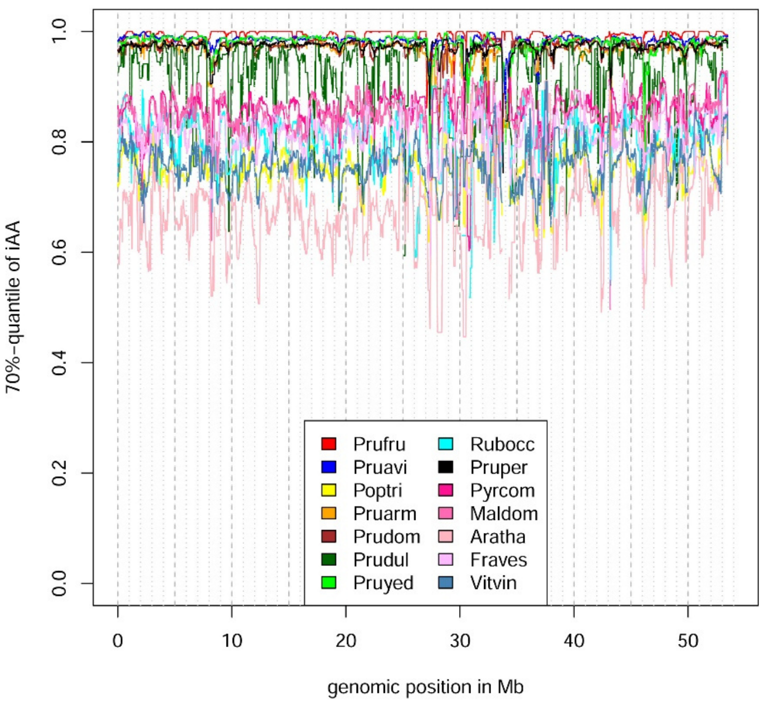


#### PCE_Fruticosa_Chro2

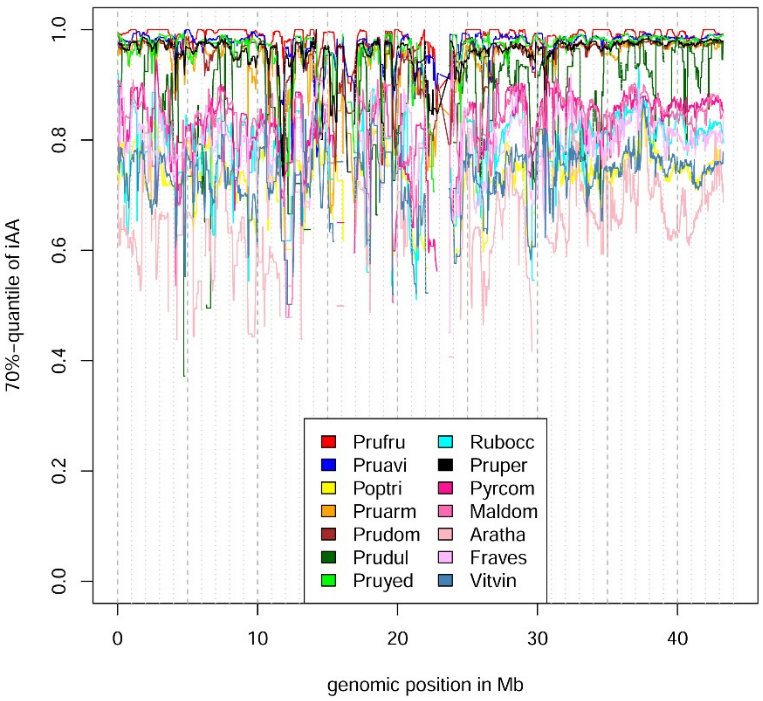

**PCE_Fruticosa_Chro3**

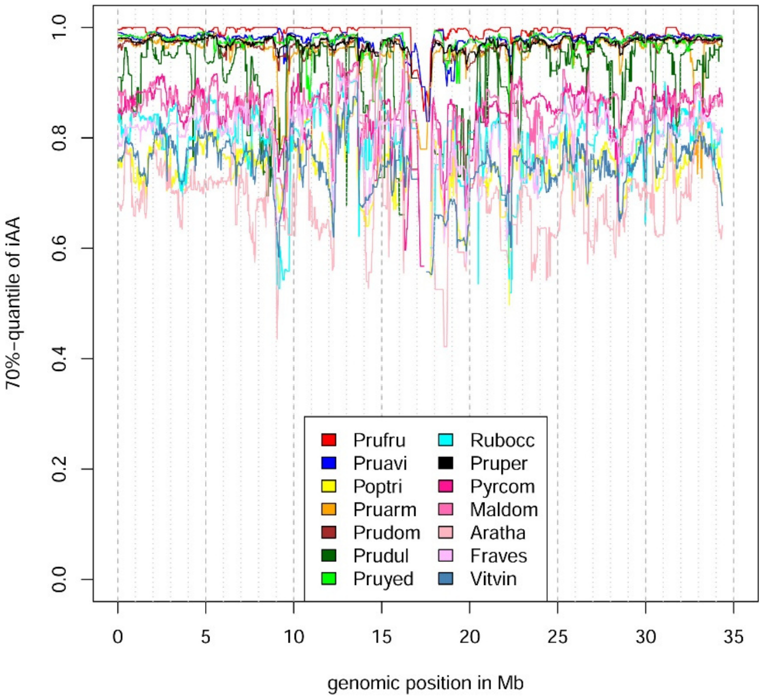


**PCE_Fruticosa_Chro4**

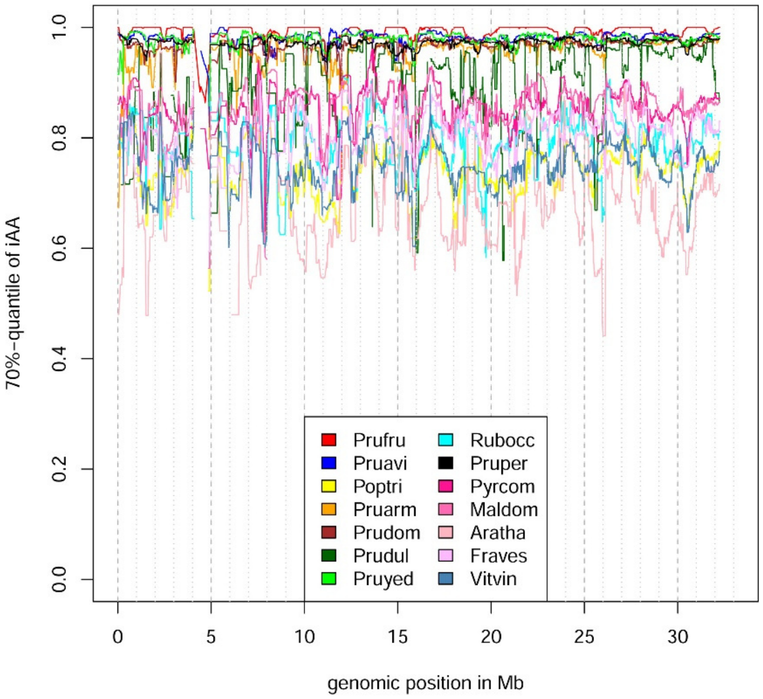

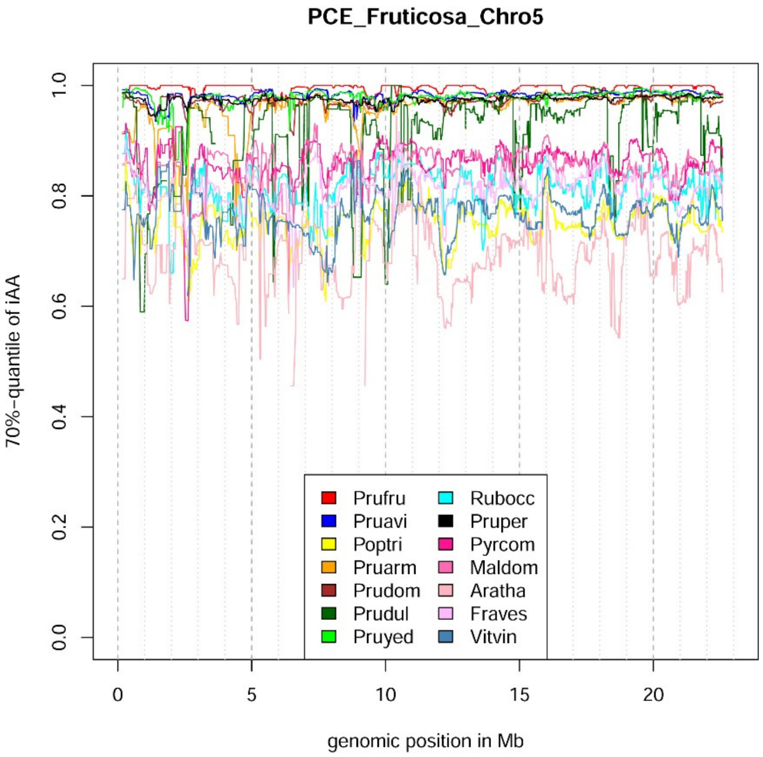
PCE_Fruticosa_Chro5
70%-quantile of iAA
0.0
0.2
0.4
0.6
0.8
1.0
0
5
10
15
20
genomic position in Mb
Prufru
Pruavi
Poptri
Pruarm
Prudom
Prudul
Pruyed
Rubocc
Pruper
Pyrcom
Maldom
Aratha
Fraves
Vitvin

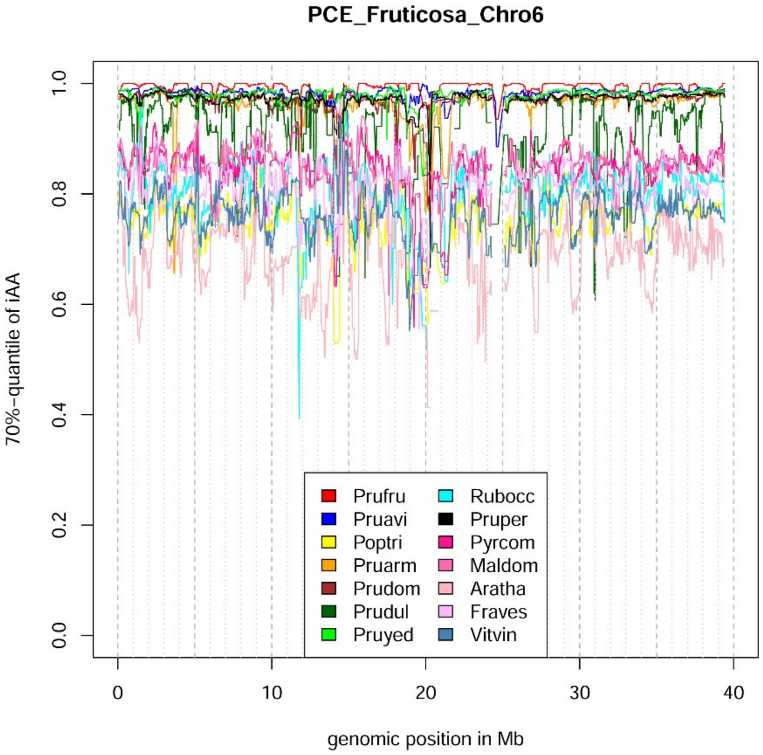
PCE_Fruticosa_Chro6
70%-quantile of iAA
0.0
0.2
0.4
0.6
0.8
1.0
0
10
20
30
40
genomic position in Mb
Prufru
Pruavi
Poptri
Pruarm
Prudom
Prudul
Pruyed
Rubocc
Pruper
Pyrcom
Maldom
Aratha
Fraves
Vitvin

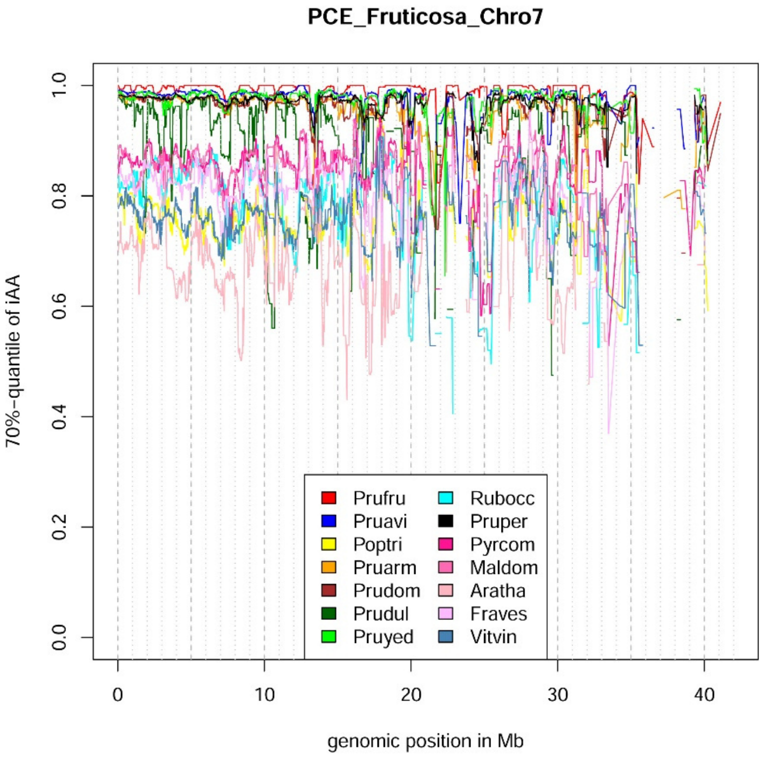

PCE_Fruticosa_Chro7
70%-quantile of iAA
0.0
0.2
0.4
0.6
0.8
1.0
0
10
20
30
40
genomic position in Mb
Prufru
Pruavi
Poptri
Pruarm
Prudom
Prudul
Pruyed
Rubocc
Pruper
Pyrcom
Maldom
Aratha
Fraves
Vitvin


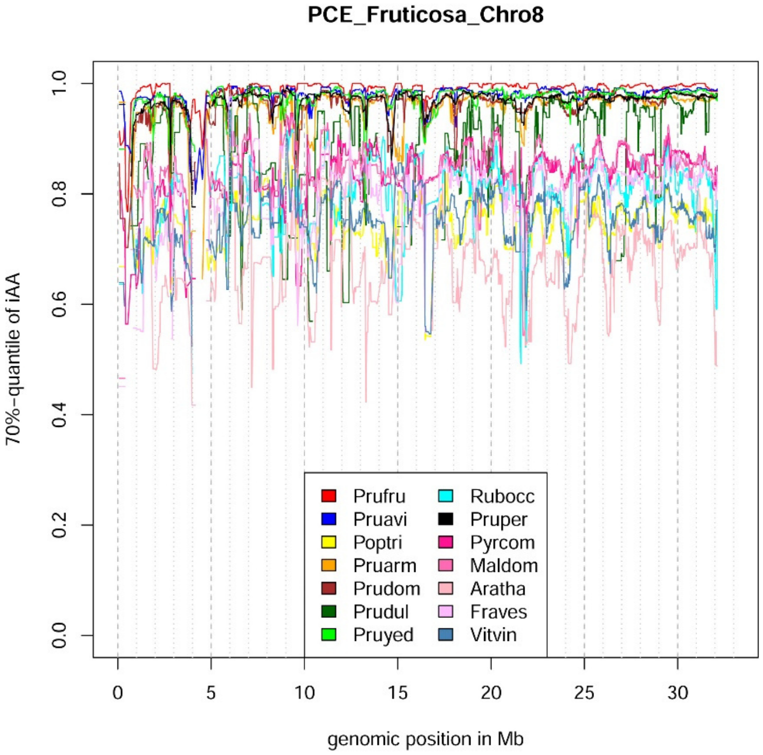

PCE_Fruticosa_Chro8
70%-quantile of iAA
0.0
0.2
0.4
0.6
0.8
1.0
0
5
10
15
20
25
30
genomic position in Mb
Prufru
Pruavi
Poptri
Pruarm
Prudom
Prudul
Pruyed
Rubocc
Pruper
Pyrcom
Maldom
Aratha
Fraves
Vitvin

**Note S3** Calculation of %-covered bases obtained from RNAseq data.

The *Pce*$_S$ genome was divided into 250k windows. The percentage of covered bases using RNAseq data of *P. canescens* (SRX14816137), *P. serrulata* (SRX14816136), *P. mahaleb* (SRX14816140), *P. pensylvanica* (SRX14816144), *P. maackii* (SRX14816139), *P. subhirtella* (SRX14816145) was estimated for each window at a depth of 1. The results were plotted with a standard script using R.

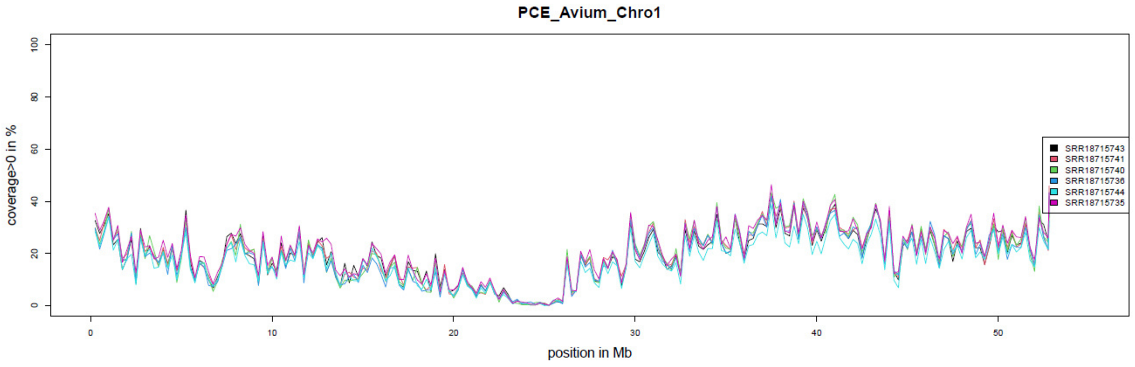


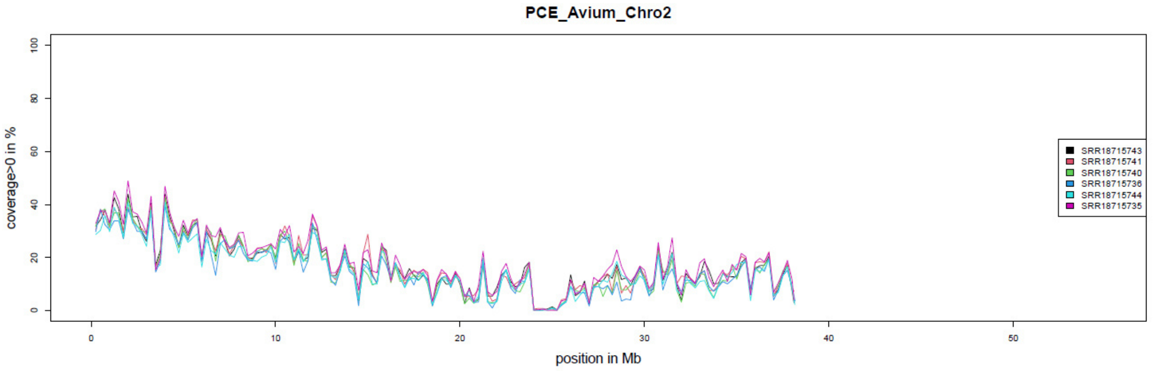


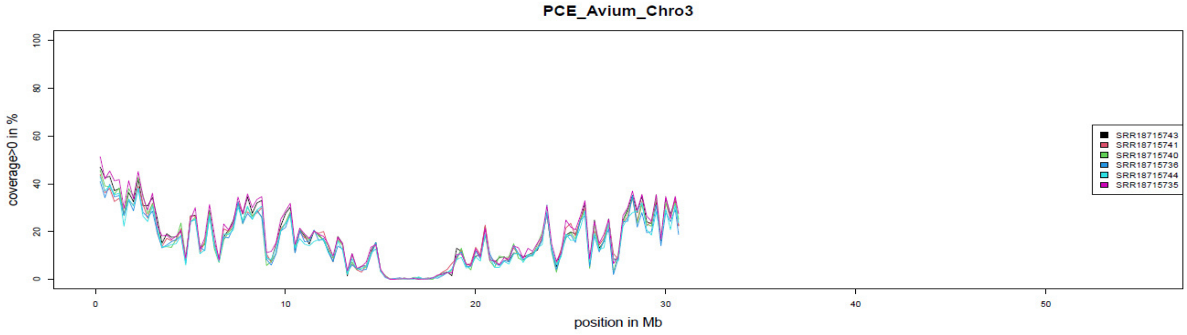


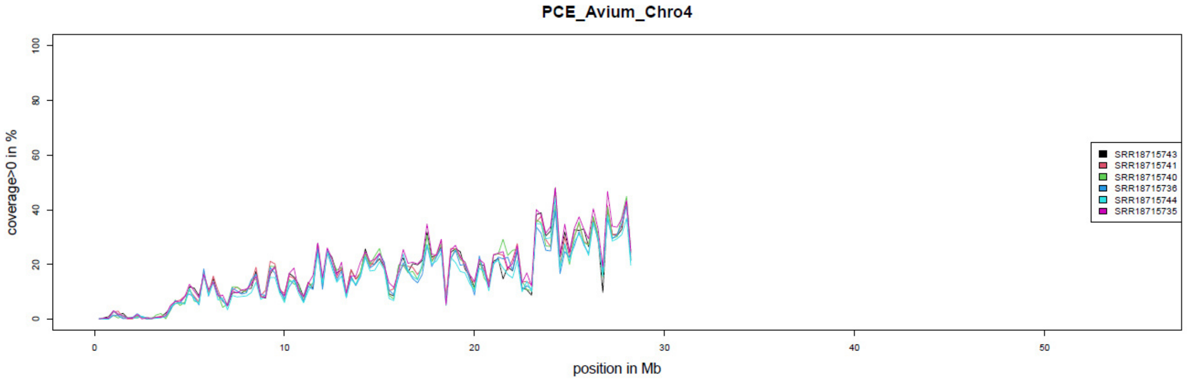

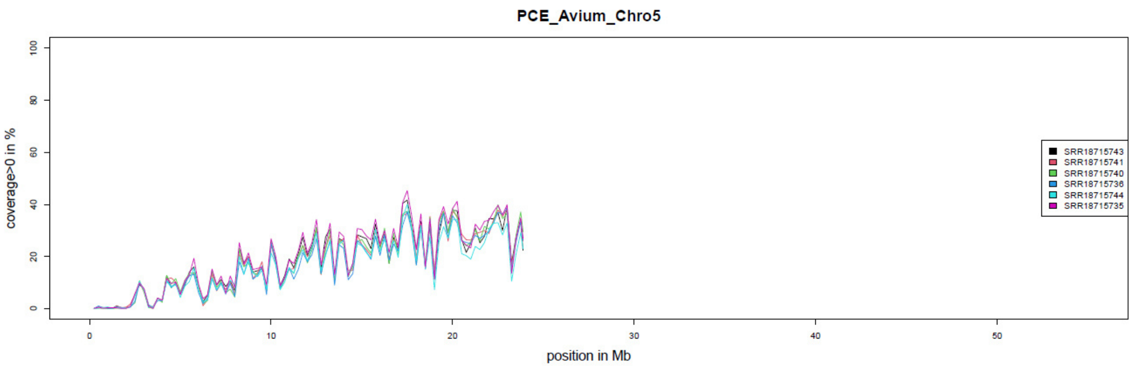
PCE_Avium_Chro5
coverage>0 in %
position in Mb
SRR18715743
SRR18715741
SRR18715740
SRR18715736
SRR18715744
SRR18715735

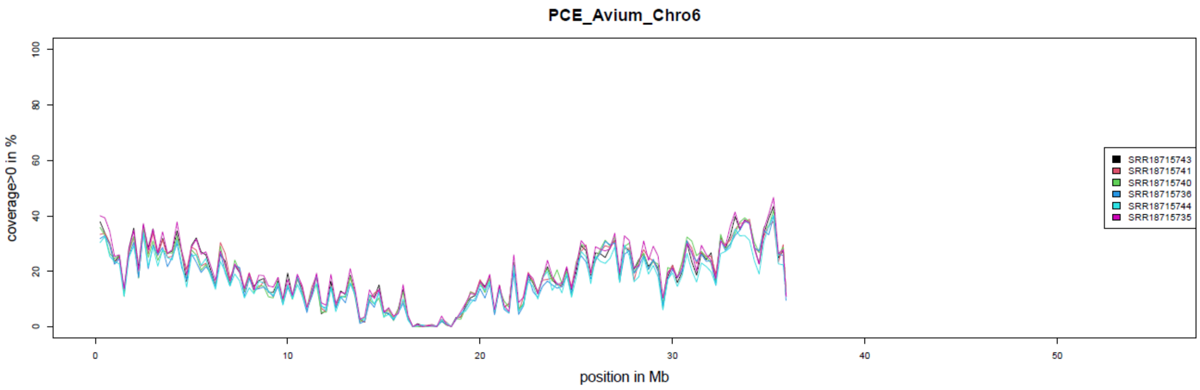
PCE_Avium_Chro6
coverage>0 in %
position in Mb
SRR18715743
SRR18715741
SRR18715740
SRR18715736
SRR18715744
SRR18715735

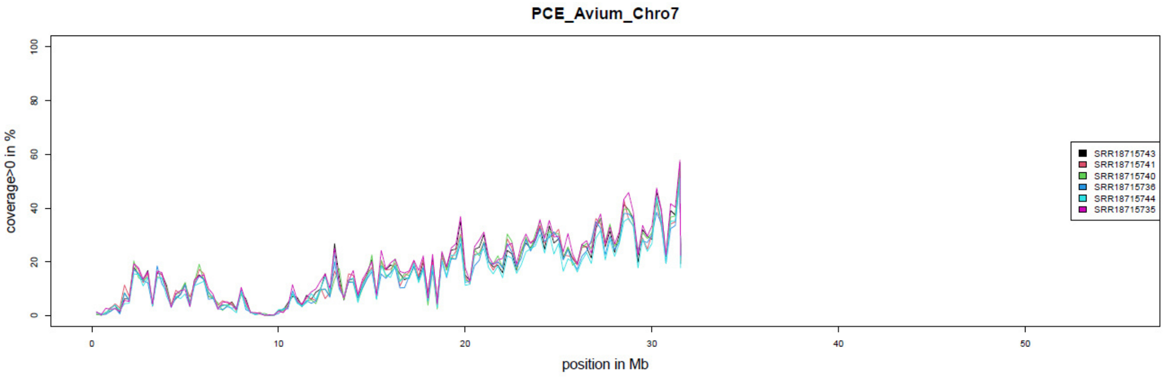
PCE_Avium_Chro7
coverage>0 in %
position in Mb
SRR18715743
SRR18715741
SRR18715740
SRR18715736
SRR18715744
SRR18715735

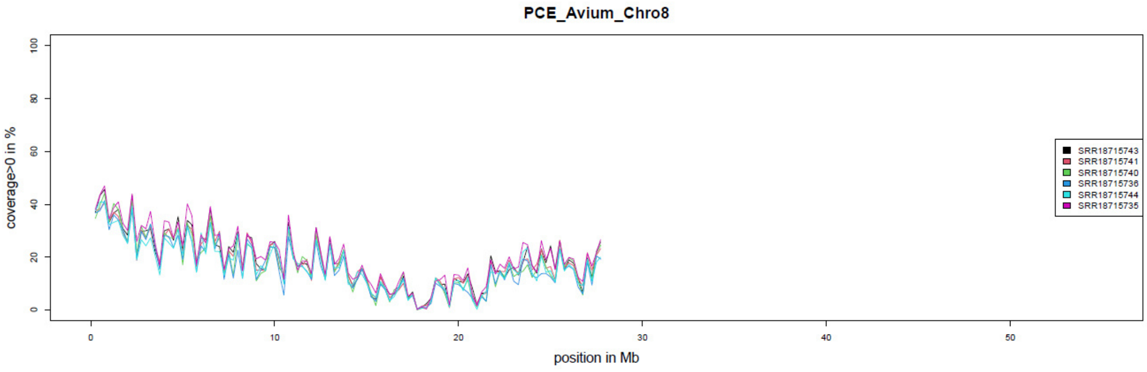
PCE_Avium_Chro8
coverage>0 in %
position in Mb
SRR18715743
SRR18715741
SRR18715740
SRR18715736
SRR18715744
SRR18715735
